# Detecting adaptive introgression in human evolution using convolutional neural networks

**DOI:** 10.1101/2020.09.18.301069

**Authors:** Graham Gower, Pablo Iáñez Picazo, Matteo Fumagalli, Fernando Racimo

## Abstract

Studies in a variety of species have shown evidence for positively selected variants introduced into one population via introgression from another, distantly related population—a process known as adaptive introgression. However, there are few explicit frameworks for jointly modelling introgression and positive selection, in order to detect these variants using genomic sequence data. Here, we develop an approach based on convolutional neural networks (CNNs). CNNs do not require the specification of an analytical model of allele frequency dynamics, and have outperformed alternative methods for classification and parameter estimation tasks in various areas of population genetics. Thus, they are potentially well suited to the identification of adaptive introgression. Using simulations, we trained CNNs on genotype matrices derived from genomes sampled from the donor population, the recipient population and a related non-introgressed population, in order to distinguish regions of the genome evolving under adaptive introgression from those evolving neutrally or experiencing selective sweeps. Our CNN architecture exhibits 95% accuracy on simulated data, even when the genomes are unphased, and accuracy decreases only moderately in the presence of heterosis. As a proof of concept, we applied our trained CNNs to human genomic datasets—both phased and unphased—to detect candidates for adaptive introgression that shaped our evolutionary history.

## Introduction

Ancient DNA studies have shown that human evolution during the Pleistocene was characterised by numerous episodes of interbreeding between distantly related groups (Green *et al*., 2010; Reich *et al*., 2010; Meyer *et al*., 2012; Prüfer *et al*., 2017; Kuhlwilm *et al*., 2016). We now know, for example, that considerable portions of the modern human gene pool derive from Neanderthals and Denisovans (Green *et al*., 2010; Reich *et al*., 2010; Prüfer *et al*., 2014). In the past few years, several methods have been developed to identify regions of present-day or ancient human genomes containing haplotypes that were introgressed from other groups of hominins. These include methods based on probabilistic models (Sankararaman *et al*., 2014, 2016; Steinrücken *et al*., 2018; Racimo *et al*., 2017a), on summary statistics (Vernot & Akey, 2014; Vernot *et al*., 2016; Racimo *et al*., 2017b) and on ancestral recombination graph reconstructions (Kuhlwilm *et al*., 2016; Hubisz *et al*., 2020; Speidel *et al*., 2019). Presumably, some of the introgressed material may have had fitness consequences in the recipient populations. While recent evidence suggests that a large proportion of Neanderthal ancestry was likely negatively selected (Harris & Nielsen, 2016; Juric *et al*., 2016), there is also support for positive selection on a smaller proportion of the genome—a phenomenon known as adaptive introgression (AI) (Whitney *et al*., 2006; Hawks & Cochran, 2006; Racimo *et al*., 2015).

Genomic evidence for AI has been found in numerous other species, including butterflies (Pardo-Diaz *et al*., 2012; Enciso-Romero *et al*., 2017), mosquitoes (Norris *et al*., 2015), hares (Jones *et al*., 2018), poplars (Suarez-Gonzalez *et al*., 2016) and monkeyflowers (Hendrick *et al*., 2016). A particularly striking example is AI in dogs, which appears to show strong parallels to AI in humans when occupying the same environmental niches. For example, a variant of the gene *EPAS1* has been shown to have introgressed from an archaic human population into the ancestors of Tibetans, and subsequently risen in frequency in the latter population, as a consequence of positive selection to high altitude (Huerta-Sánchez *et al*., 2014). A different high-frequency *EPAS1* variant is also uniquely found in Tibetan Mastiffs, and appears to also have introgressed into this gene pool via admixture with a different species, in this case Tibetan wolves (Miao *et al*., 2016), likely due to the same selective pressures.

To detect AI, researchers can look for regions of the genome with a particularly high frequency of introgressed haplotypes from a donor species or population into a recipient species or population. These haplotypes are often detected assuming neutrality of archaic alleles since the introgression event (Vernot *et al*., 2016; Vernot & Akey, 2014; Sankararaman *et al*., 2016, 2014). Other studies have designed statistics that are sensitive to characteristic patterns left by AI, using simulations incorporating both admixture and selection (Gittelman *et al*., 2016; Racimo *et al*., 2017b). More recently, Setter *et al*. (2020) developed a likelihood framework to look for local alterations to the site frequency spectrum that are consistent with adaptive introgression, using only data from the recipient species. The main challenge that these studies face is that it is hard to jointly model selection from material introduced via admixture (Racimo *et al*., 2015).

To overcome the need to compress data into summary statistics (which might miss important features) or solve complex analytical theory, deep learning techniques are increasingly becoming a popular solution to address problems in population genetics. These problems include the inference of demographic histories (Sheehan & Song, 2016; Flagel *et al*., 2018; Villanea & Schraiber, 2019; Mondal *et al*., 2019; Sanchez *et al*., 2020), admixture (Blischak *et al*., 2020), recombination (Chan *et al*., 2018; Flagel *et al*., 2018; Adrion *et al*., 2020b) and natural selection (Schrider & Kern, 2018; Sheehan & Song, 2016; Torada *et al*., 2019; Isildak *et al*., 2020). Deep learning is a branch of machine learning that relies on algorithms structured as multi-layered networks, which are trained using known relationships between the input data and the desired output. They can be used for classification, prediction or data compression (Aggarwal *et al*., 2018). Among the techniques in this field, convolutional neural networks (CNNs) are a family of methods originally designed for image recognition and segmentation (LeCun *et al*., 1995; Krizhevsky *et al*., 2012), which have been recently applied to population genetic data (Chan *et al*., 2018; Flagel *et al*., 2018; Torada *et al*., 2019; Isildak *et al*., 2020; Blischak *et al*., 2020; Sanchez *et al*., 2020). A CNN can learn complex spatial patterns from large datasets that may be informative for classification or prediction, using a series of linear operations known as convolutions, to compress the data into features that are useful for inference.

Despite the recent advances in deep learning for population genetics, no significant attempts have been proposed to identify AI from population genomic data. Here, we develop a deep learning method called genomatnn that jointly models archaic admixture and positive selection, in order to identify regions of the genome under adaptive introgression. We trained a CNN to learn relevant features directly from a genotype matrix at a candidate region, containing data from the donor population, the recipient population and a unadmixed outgroup. The method has *>*88 % precision to detect AI, and is effective on both ancient and recently selected introgressed haplotypes. We then applied our method to population genomic datasets where the donor population is either Neanderthals or Denisovans and the recipient populations are Europeans or Melanesians, respectively. In each case, we used the Yoruba population as a unadmixed outgroup and we were able to both recover previously identified AI regions and unveil new candidates for AI in human history.

## Results

### A CNN for detecting adaptive introgression

In our method, we assumed we have sequence data from multiple populations: the donor population and the recipient population in an admixture event, as well as an unadmixed population that is a sister group to the recipient (Fig. 1A). We constructed an *n × m* matrix for *n* haplotypes (or diploid genotypes, for unphased data), where each entry corresponds to the count of minor alleles in an individual’s haplotype (or diploid genotype), for a <u>^100^</u> kbp region of the genome. Within each population, we sorted these pseudo-haplotypes (or genotypes) according to similarity to the donor population, and concatenated the matrices for each of the populations into a single pseudo-genotype matrix (Fig. 1B).

**Figure 1:**
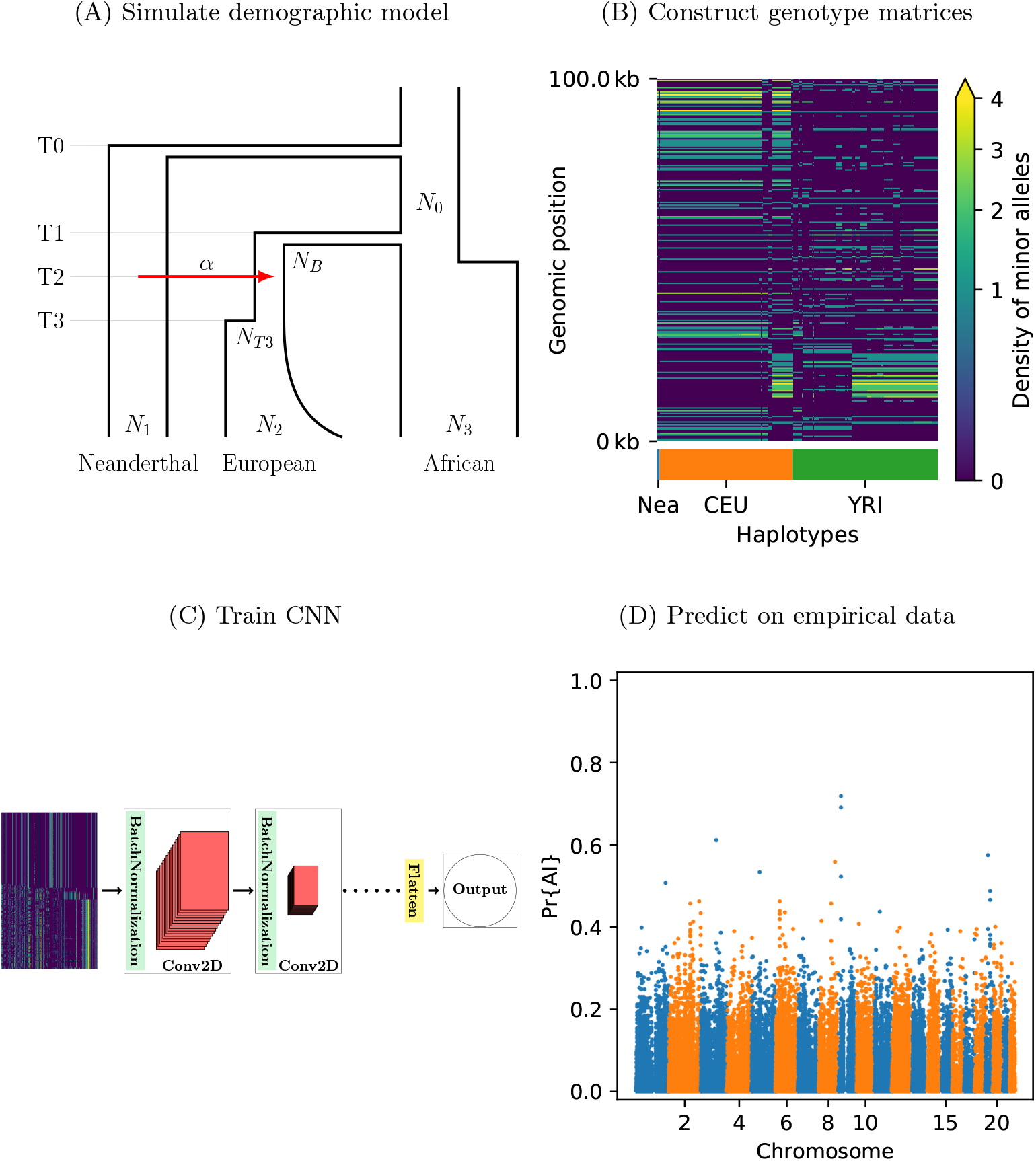
A schematic overview of how genomatnn detects adaptive introgression. We first simulate a demographic history, such as the HomininComposite_4G20 model shown in Fig. 1A, using the SLiM engine in stdpopsim. Parameter values for this model are given in Table S1. Three distinct scenarios are simulated for a given demographic model: neutral mutations only, a sweep in the recipient population, and adaptive introgression. The tree sequence file from each simulation is converted into a genotype matrix for input to the CNN. Fig. 1B shows a genotype matrix from an adaptive introgression simulation, where lighter pixels indicate a higher density of minor alleles, and haplotypes within each population are sorted left-to-right by similarity to the donor population (Nea). In this example, haplotype diversity is low in the recipient population (CEU), which closely resembles the donor (Nea). Thousands of simulations are produced for each simulation scenario, and their genotype matrices are used to train a binary-classification CNN (Fig. 1C). The CNN is trained to output Pr[AI], the probability that the input matrix corresponds to adaptive introgression. Finally, the trained CNN is applied to genotype matrices derived from a VCF/BCF file (Fig. 1D).

**Table 1:**
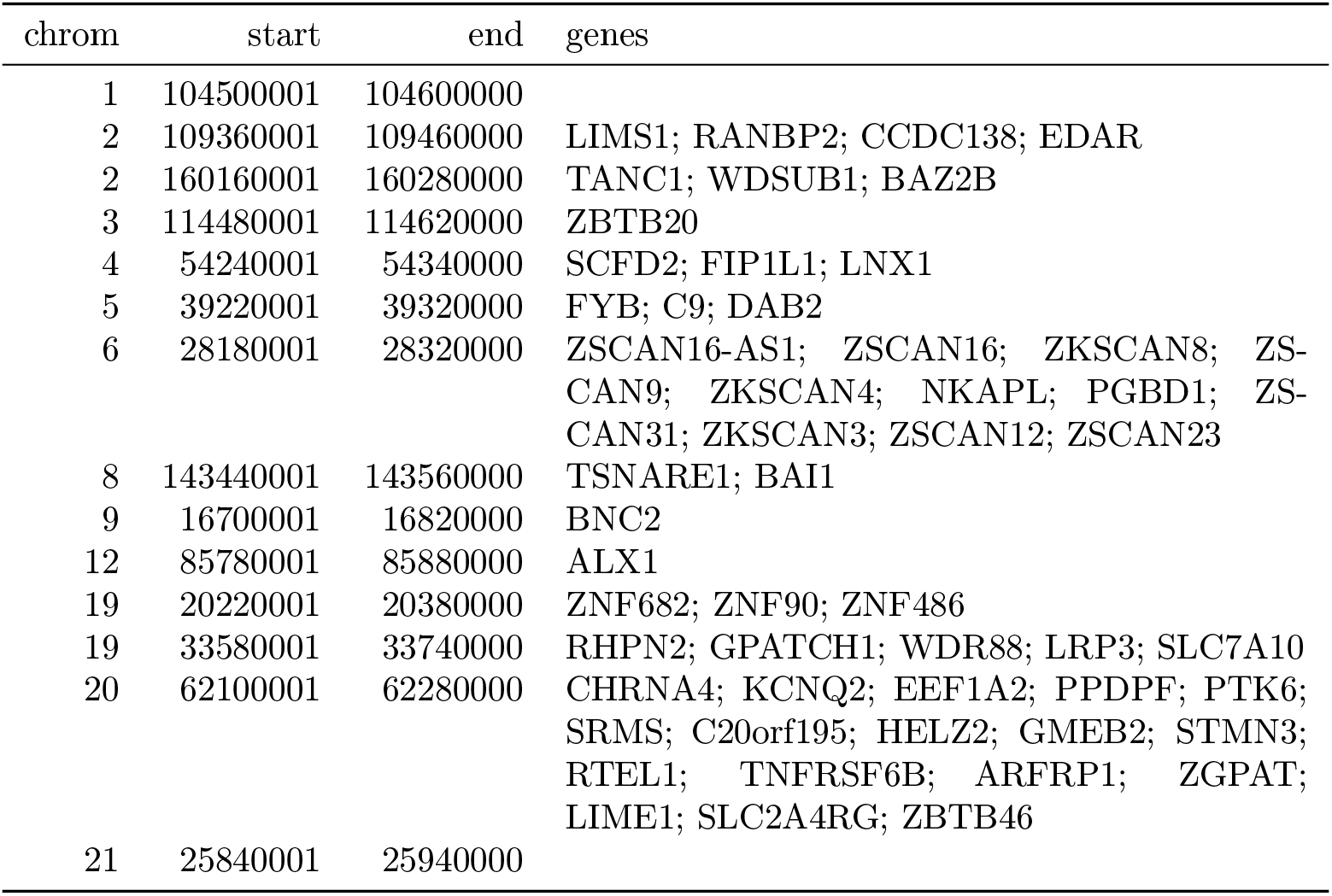
Top ranking gene candidates corresponding to Neanderthal AI in Europeans. We show genes which overlap, or are within 100 kbp of, the 30 highest ranked 100 kbp intervals. Adjacent intervals have been merged. The CNN was trained using only AI simulations with selected mutation having allele frequency > 0.25, and subsequently calibrated with resampled neutral:sweep:AI ratios of 1:0.1:0.02.

We designed a CNN (Fig. 1C) that takes this concatenated matrix as input to distinguish between adaptive introgression scenarios and other types of neutral or selection scenarios. The CNN was trained using simulations, and uses a series of convolution layers with successively smaller outputs, to extract increasingly higher-level features of the genotype matrices—features which are simultaneously informative of introgression and selection. The CNN outputs the probability that the input matrix comes from a genomic region that underwent adaptive introgression. As our simulations used a wide range of selection coefficients and times of selection onset, the network does not assume these parameters are known a priori, and is able to detect complete or incomplete sweeps at any time after gene flow.

Our method has several innovative features relative to previous population genetic implementations of CNNs (described extensively in the Methods section). For example, when loading the genotype matrices as input, we implemented an image resizing scheme that leads to fast training times, while avoiding the drawbacks of similar methods (Torada *et al*., 2019), by preserving inter-allele distances and thus the local density of segregating sites. Additionally, instead of using pooling layers, we used a 2×2 step size when performing convolutions. This has the same effect as pooling, in that the output size is smaller than the input, so the accuracy of the model is unaffected relative to traditional implementations of CNNs, but it has a much lower computational burden (Springenberg *et al*., 2015).

Furthermore, we incorporated a framework to visualise the features of the input data that draw the most attention from the CNN, by plotting saliency maps from the keras-vis library (Kotikalapudi & contributors, 2017). Saliency maps can help to understand which regions of the genotype matrix contribute the most toward the CNN prediction score (Fig. 3).

We also provide downloadable pre-trained CNNs as well as a pipeline for training new CNNs (see Methods). These interface with a new selection module that we designed and incorporated into the stdpopsim framework (Adrion *et al*., 2020a), using the forwards-in-time simulator SLiM (Haller & Messer, 2019). We believe this will facilitate the application of the method to other datasets, allowing users to modify its parameters according to the specific requirements of the biological system under study.

### Performance on simulations

We aimed to assess the performance of our method on simulations. We performed simulations under two different demographic models:

- Demographic model A: a three-population model including an African, a European and a Neanderthal population, with Neanderthal gene flow into Europeans (Fig. 1A)
- Demographic model B: a more complex model, including an African, a Melanesian, a Neanderthal and a Denisovan population, with two pulses of Denisovan gene flow into Melanesians, plus a pulse of Neanderthal gene flow into non-Africans, based on Jacobs *et al*. (2019) (Fig. S1).

When training a CNN on Demographic Model A using phased data, we obtained a precision of 90.2 % (proportion of AI predictions that were AI simulations) and 97.9 % negative predictive value (NPV; proportion of “not-AI” predictions that were either neutral or sweep simulations) (Figs. 2 and S8). The network output higher probabilities for AI simulations with larger selection coefficients, and for older times of onset of selection. We also observed that the network falsely classified neutral simulations as AI more frequently than it falsely classified sweep simulations. When the CNN was trained on this same demographic model assuming genotypes were unphased, the results were very similar, with 88.1 % precision and 98.7 % NPV (Fig. S7).

**Figure 2:**
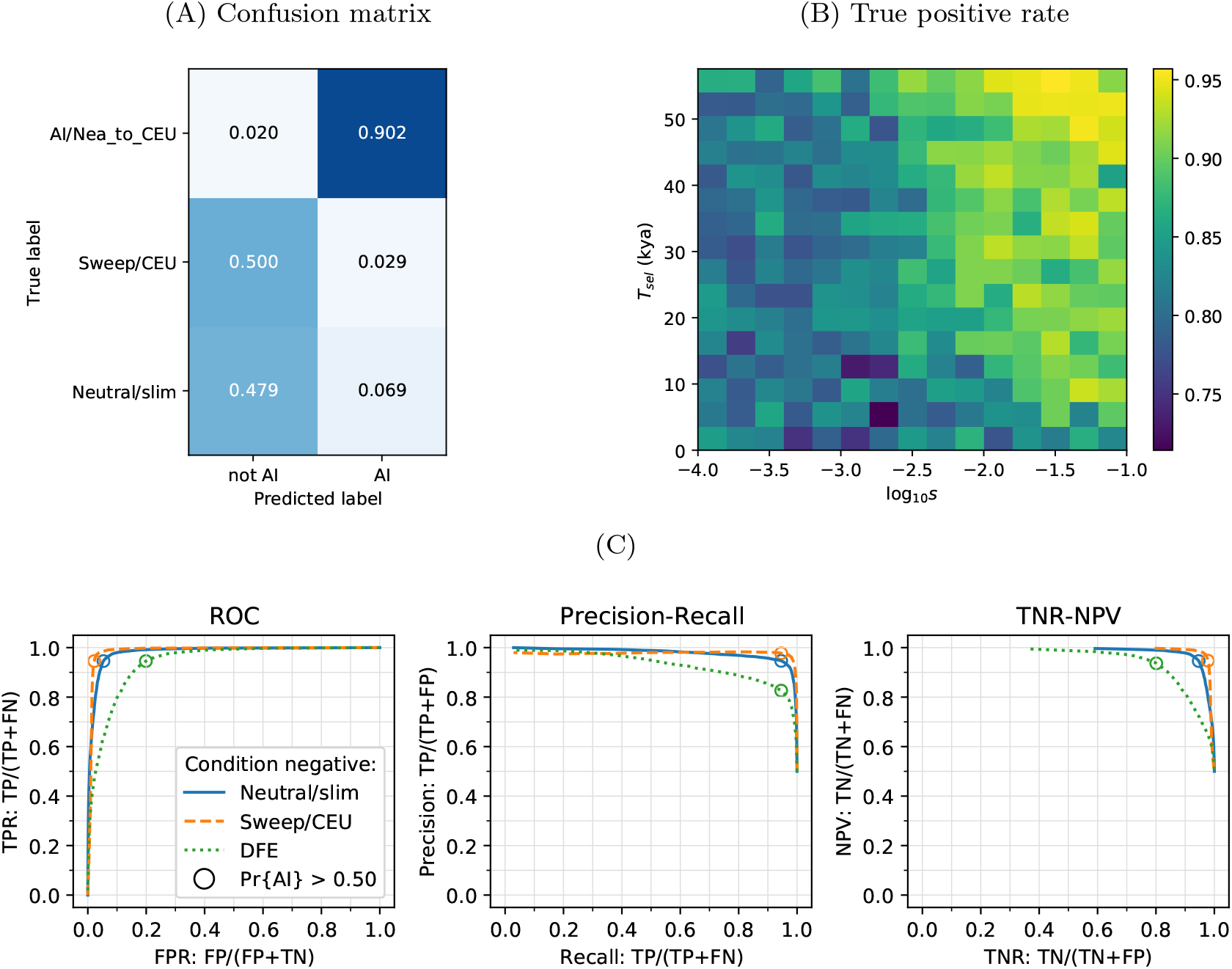
CNN performance on validation simulations for Demographic Model A. The CNN was trained using only AI simulations with selected mutation having allele frequency > 0.25. Fig. 2A: Confusion matrix. For the two prediction categories, either “not AI” or AI, we show the proportion attributed to each of the true (simulated) scenarios. Fig. 2B: Average CNN prediction for AI scenarios, binned by selection coefficient, *s*, and time of onset of selection *T_sel_*. Fig. 2C: ROC curves, precision-recall curves and True Negative Rate vs. Negative Predictive Value (TNR-NPV) curves. The positive condition is AI. The negative conditions are shown using different line styles/colours. The circles indicate the point in ROC-space (respectively Precision-Recall-space, and TNR-NPV-space) when using the threshold Pr[AI] > 0.5 for classifying a genotype matrix as AI. DFE: distribution of fitness effects. TP: true positives; FP: false positives; TPR: true positive rate; FPR: false positive rate; ROC: Receiver operating characteristics; TNR: true negative rate; TPR: true positive rate.

When training a CNN on Demographic Model B (assuming unphased genotypes, as accurately phased data is not readily available for Melanesian genomes), we obtained 88.8 % precision and 82.5 % NPV (Figs. S9 and S10). We note here that the network had greater precision when detecting AI derived from the more ancient pulse of Denisovan gene flow than the younger pulse.

Kim *et al*. (2018) and Zhang *et al*. (2020) recently suggested that introduced genetic material can mask deleterious recessive variation and produce a signal very similar to adaptive introgression. To assess whether heterosis following introgression affects the false positive rates in our CNN, we simulated a distribution of fitness effects (DFE) with recessive dominance for 70 % of derived mutations (the rest were simulated as neutral), and found this only slightly increases the false positive rate (Figs. 2 and S8).

We further tested whether the method was robust to demographic misspecification, by evaluating the CNN trained on Demographic Model A against simulations for Demographic Model B. As there are more Melanesian individuals than European individuals in our simulations (because we aimed to mimic the real number of genomes available in our data analysis below), we down-sampled the Melanesian genomes to match the number of European genomes, so as to perform a fair misspecification comparison. In this case, we found the precision dropped to 65.3 % and the NPV to 74.4 % (Fig. S6).

### Network attention

To understand which features of the input matrices were used by the CNN to make its predictions, we constructed saliency maps (Simonyan *et al*., 2014). This technique works by computing the gradient of a network’s output with respect to a single input. Thus, highlighted regions from the saliency map indicate where small changes in the input matrix have a relatively large influence over the CNN output prediction. We calculated an average saliency map for each output category predicted by the network (AI or not-AI), for a CNN trained on Demographic Model A (Fig. 3). Our results show that when the network was presented with an AI matrix, it focused most of the attention on the Neanderthal and European haplotypes, while not putting much emphasis on the African haplotypes. In non-AI scenarios, the network focused sharply on the Neanderthal and left-most European haplotypes. The saliency maps also show a concentration of attention in the central region of the genomic window, around where the selected mutation was drawn (even though this mutation was removed before constructing genotype matrices; see Methods).

**Figure 3:**
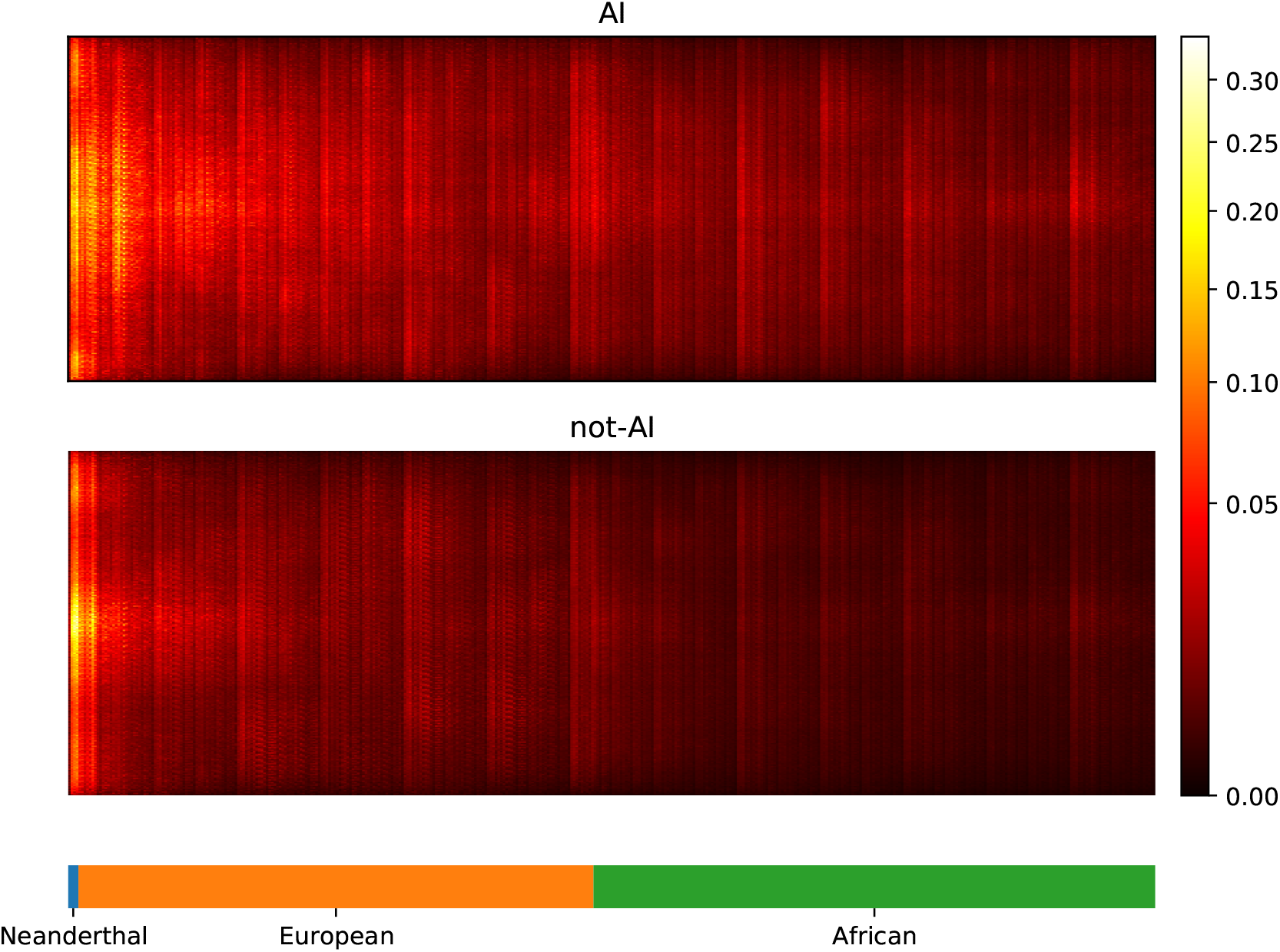
Saliency maps, showing the CNN’s attention across the input matrices for AI and not-AI inputs, calculated for the CNN trained on Demographic Model A, filtered for beneficial allele frequency > 0.25. Each panel shows the average gradient over input matrices encoding AI (top) or not-AI (bottom). Brighter colours indicate larger gradients, where small changes in the genotype matrix have a relatively larger influence over the CNN’s prediction. Columns in the input matrix correspond to haplotypes from the populations labelled at the bottom.

### Calibration

We implemented a score calibration scheme to account for the fact that our simulation categories (neutrality, sweep and AI) will be highly imbalanced in real data applications (Guo *et al*., 2017; Kull *et al*., 2017). CNN classifiers sometimes produce improperly calibrated probabilities (Guo *et al*., 2017). In our case, this occurs because the proportion of each category is not known in reality, and thus does not match the simulated proportion. For this reason, we fitted our calibration procedure using training data resampled with various ratios of neutral:sweep:AI simulations (Fig. 4). We tested different calibration methods by fitting the calibrator to the training dataset, and inspecting reliability plots and the sum of residuals on a validation dataset (see Methods).

**Figure 4:**
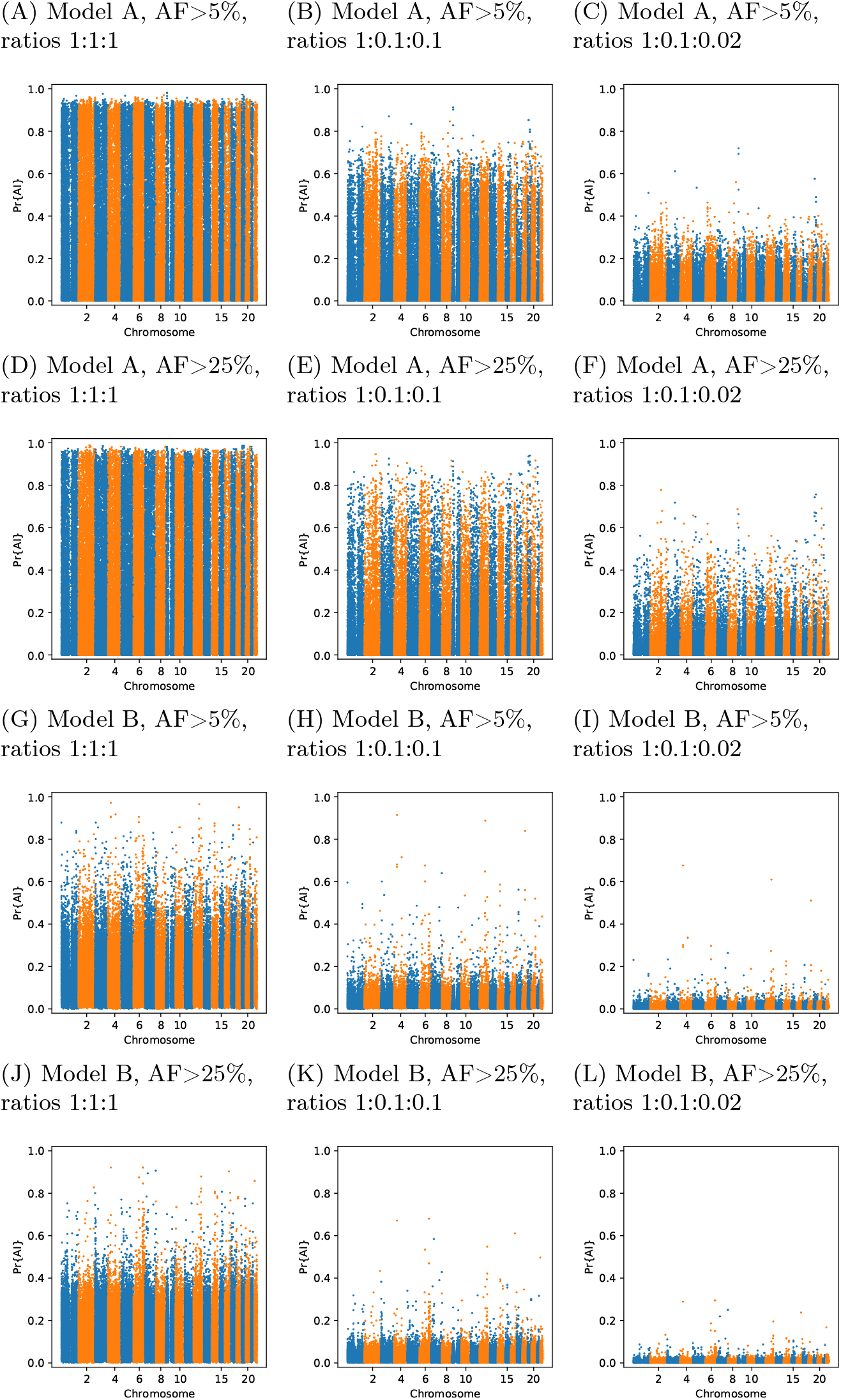
Comparison of Manhattan plots using beta-calibrated output probabilities for different class ratios. Each row indicates a single CNN, with equivalent data filtering. Each column indicates a different ratio of scenarios used for calibration. AF = Minimum beneficial allele frequency.

**Figure 5:**
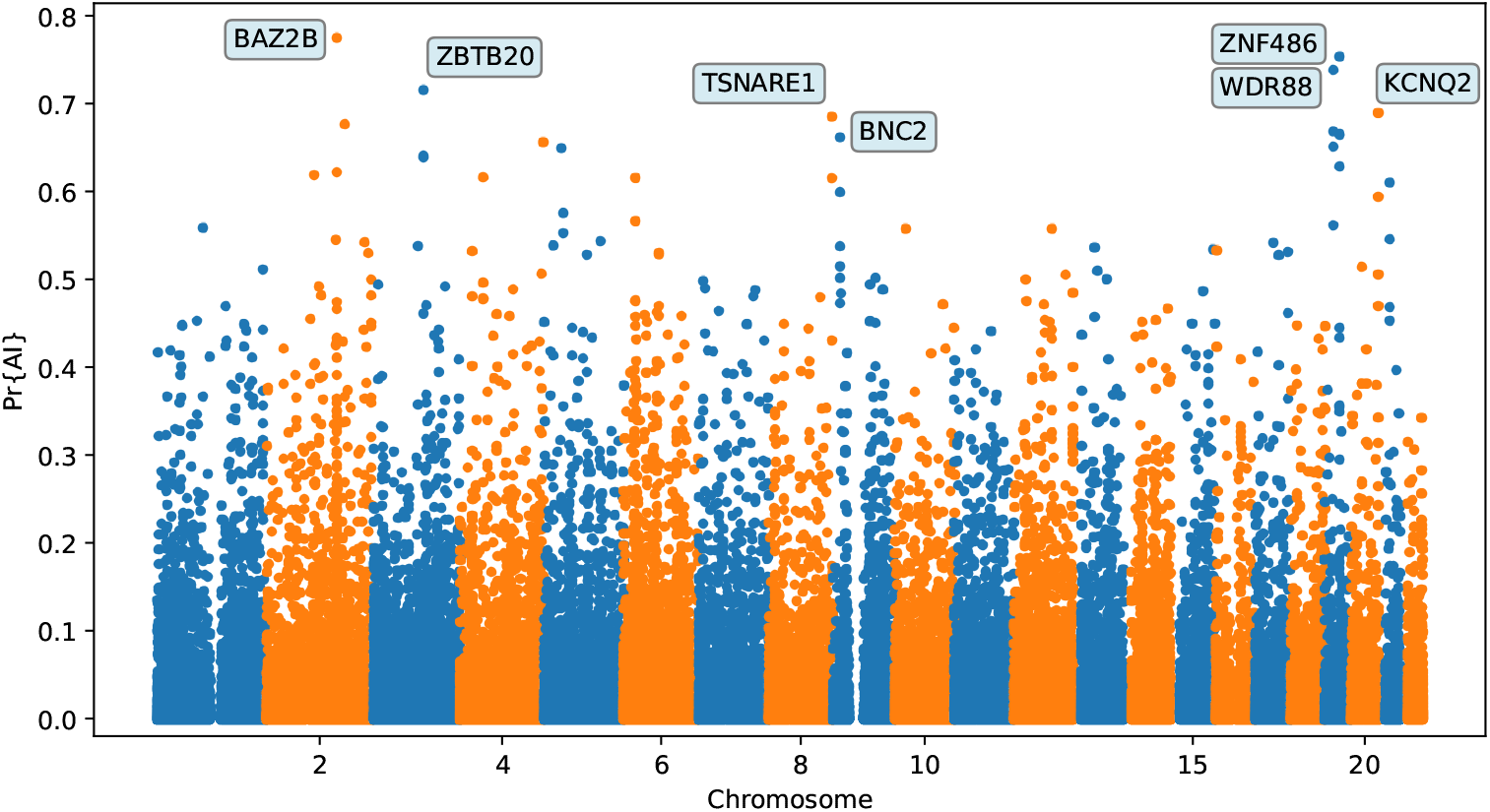
Application of the trained CNN to the Vindija and Altai Neanderthals, and 1000 genomes populations YRI and CEU. The CNN was applied to overlapping 100 kbp windows, moving along the chromosome in steps of size 20 kbp. The CNN was trained using only AI simulations with selected mutation having allele frequency > 25%, and subsequently calibrated with resampled neutral:sweep:AI ratios of 1:0.1:0.02.

**Figure 6:**
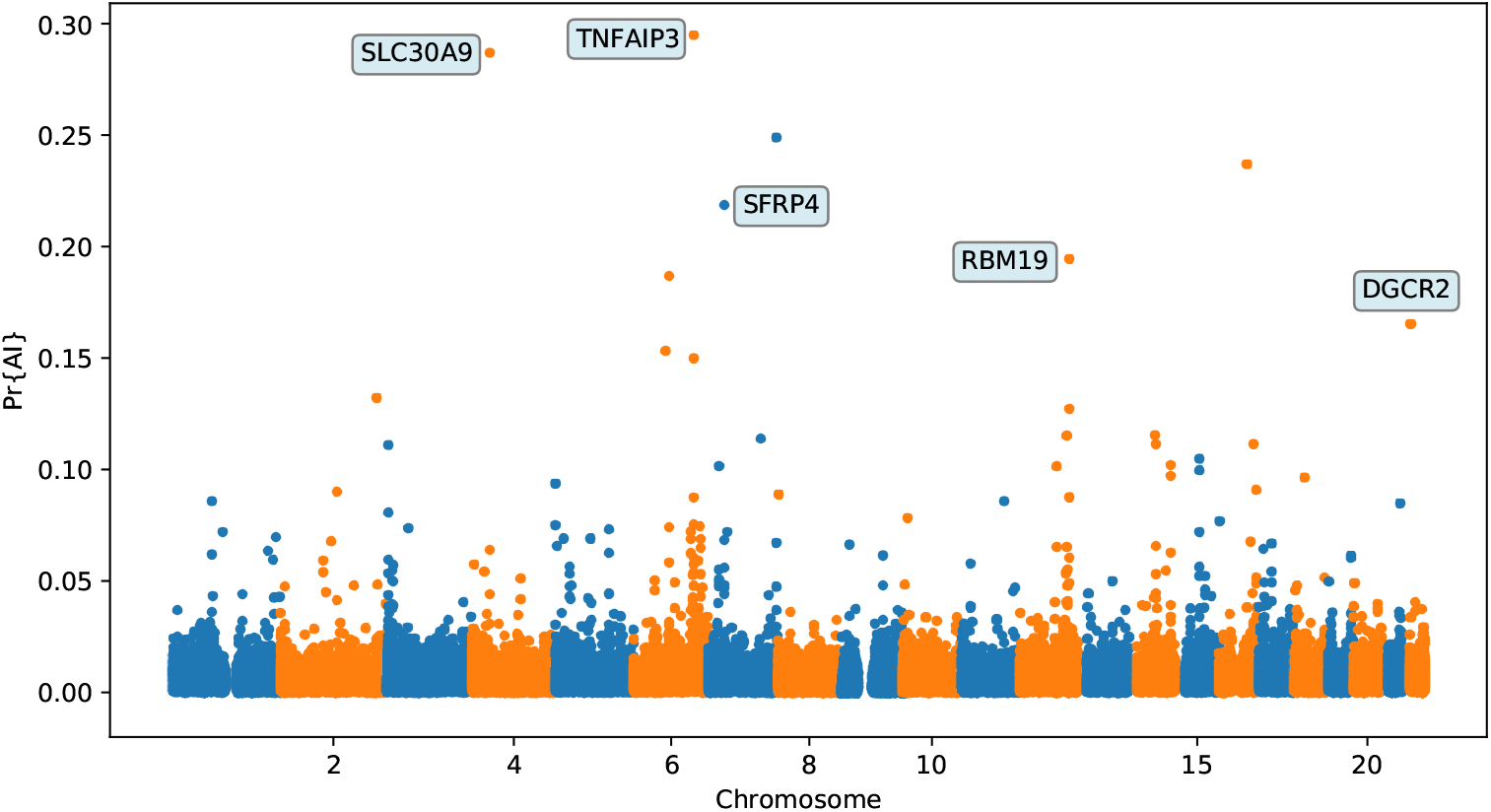
Application of the trained CNN to the Altai Denisovan and Altai Neanderthal, 1000 genomes YRI populations, and IGDP Melanesians. The CNN was applied to overlapping 100 kbp windows, moving along the chromosome in steps of size 20 kbp. The CNN was trained using only AI simulations with selected mutation having allele frequency > 25%, and subsequently calibrated with resampled neutral:sweep:AI ratios of 1:0.1:0.02.

### Candidates for Neanderthal adaptive introgression in European genomes

We applied our method to a combined genomic panel of archaic hominins (Prüfer *et al*., 2017, 2014; Meyer *et al*., 2012) and present-day humans (The 1000 Genomes Project Consortium, 2015; Jacobs *et al*., 2019), to look for regions of the genome where Non-African humans show signatures of AI from archaic hominins. First, we looked for Neanderthal introgression into the ancestors of Northwestern Europeans (CEU panel), using Yoruba Africans (YRI panel) as the unadmixed sister population. We used two different beneficial-allele frequency cutoffs for training: 5% and 25% (Tables 1 and S2). We focus here on describing the results from the 25% condition (Figs. S11 to S24). We found several candidate genes for AI that have been reported before (Sankararaman *et al*., 2014, 2016; Vernot & Akey, 2014; Gittelman *et al*., 2016; Racimo *et al*., 2017b), including *BNC2*, *KCNQ2/EEF1A2 WRD88/GPATCH1* and *TANC1*.

However, the candidate region we identify on chromosome 2 around *TANC1* extends farther downstream of this gene, also overlapping *BAZ2B* (Fig. S13). This codes for a protein related to chromatin remodelling, and may have a role in transcriptional activation. Mutations in BAZ2B have recently been associated with neurodevelopmental disorders, including developmental delay, autism spectrum disorder and intellectual disability (Scott *et al*., 2020). Additionally, we found two novel candidates for AI that have not been previously reported, spanning the regions chr6:28.18Mb–28.32Mb (Fig. S17) and chr20:62.1Mb–62.28Mb (Fig. S23), including multiple genes encoding zinc finger proteins. UK-biobank PHEWAS associations (Canela-Xandri *et al*., 2018) suggest both regions generally affect phenotypes related to blood, including platelet, erythrocyte and leukocyte counts (at the *p <* 10*^−^*^8^ level, the chr6 region has 91 hits, while the chr20 region has 19, with 10 of these traits in common).

### Candidates for Denisovan adaptive introgression in Melanesian genomes

We then looked for Denisovan AI in Melanesian genomes from the IGDP panel (Jacobs *et al*., 2019), also using Yoruba Africans as the unadmixed sister group, using two different beneficial-allele frequency cutoffs for training: 5% and 25% (Tables 2 and S3). Again, we focus on describing the results from the 25% condition (Figs. S25 to S46). Among the top candidates, we found a previously reported candidate for AI in Melanesians: *TNFAIP3* (Vernot *et al*., 2016; Gittelman *et al*., 2016). Denisovan substitutions carried by the introgressed haplotype in this gene have been found to enhance the immune response by tuning the phosphorylation of the encoded A20 protein, which is an immune response inhibitor (Zammit *et al*., 2019).

**Table 2:**
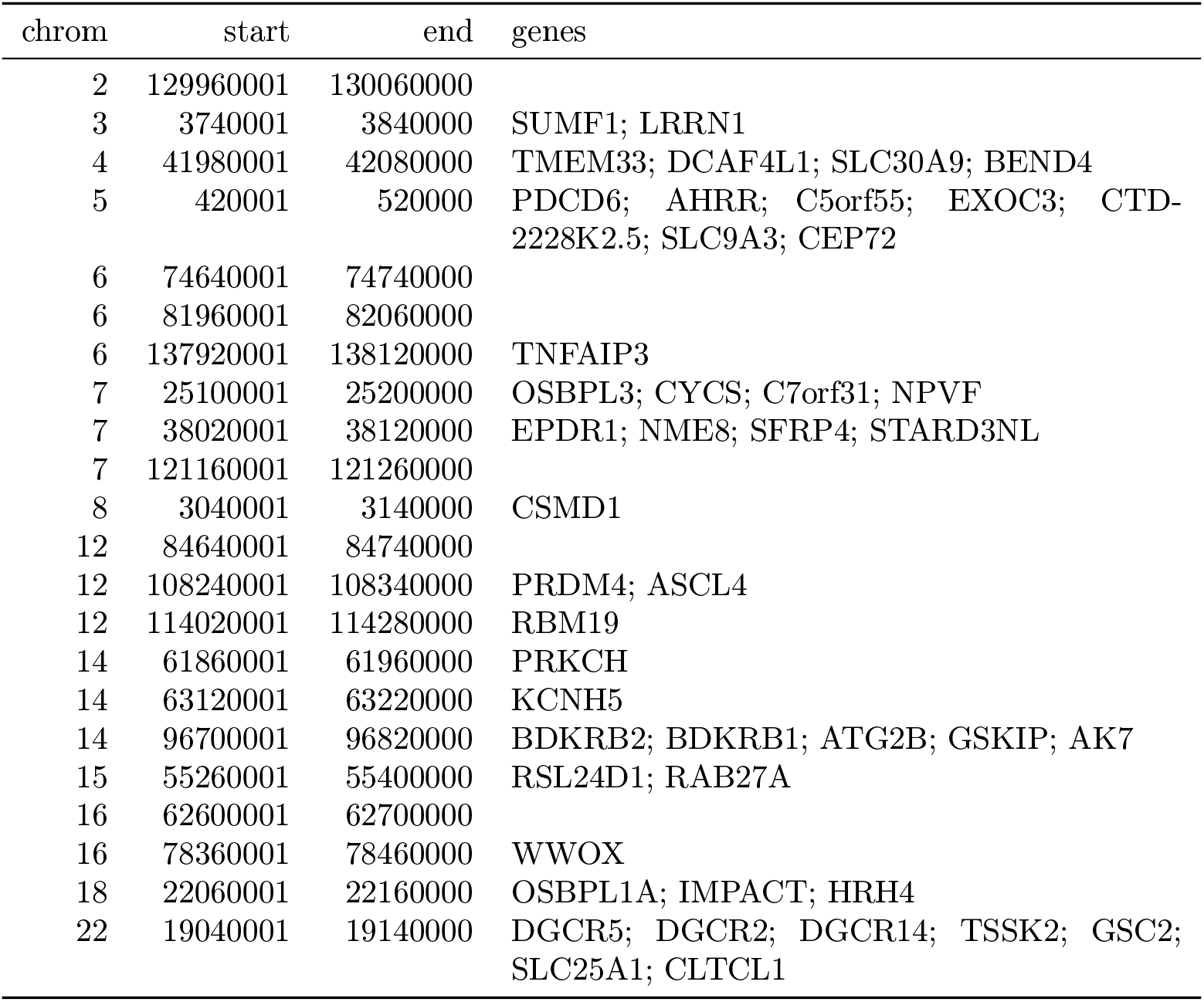
Top ranking gene candidates corresponding to Denisovan AI in Melanesians. We show genes which overlap, or are within 100 kbp of, the 30 highest ranked 100 kbp intervals. Adjacent intervals have been merged. The CNN was trained using only AI simulations with selected mutation having allele frequency > 0.25, and subsequently calibrated with resampled neutral:sweep:AI ratios of 1:0.1:0.02.

We found evidence for Denisovan AI in Melanesians at several other candidate regions. A few of these regions (or contiguous regions) were previously reported by Sankararaman *et al*. (2016) but not extensively described, possibly because the previously reported sections of those regions deemed to be introgressed were intergenic. One of the regions with strong evidence for AI (chr7:25.1Mb–25.2Mb; Fig. S32) overlaps the *CYCS* gene. This gene codes for cytochrome C: a small heme protein that plays a crucial role in the electron transport chain in mitochondria, and has been associated with various blood-related diseases, like thrombocytopenia (Morison *et al*., 2008; De Rocco *et al*., 2014; Uchiyama *et al*., 2018). Another top candidate region (chr12:108.24-108.34Mb, Fig. S37) is upstream of *PRDM4* and *ASCL4*. The former gene codes for a transcription factor that may be involved in the nerve growth factor cell survival pathway and play a role in tumour suppression (Yang & Huang, 1999). The latter gene codes for a different transcription factor that may be involved in skin development (Jonsson *et al*., 2004).

We detected signatures of Denisovan AI in a region in chromosome 3 near *SUMF1* and *LRNN1* (Fig. S26), which was also identified in Jacobs *et al*. (2019). *SUMF1* codes for an enzyme involved in the hydrolysis of sulfate esters, which has been associated with sulfatase deficiency (Cosma *et al*., 2003), while *LRNN1* encodes a protein involved in neuronal differentiation, which has been associated with neuroblastoma and Alzheimer’s disease (Bai *et al*., 2014; Hossain *et al*., 2012). Another candidate region is in chromosome 7 and is upstream of *SFRP4* (Fig. S33), which encodes a protein associated with diabetes (Mahdi *et al*., 2012) and Pyle’s disease (Simsek Kiper *et al*., 2016). Moreover, there is also a candidate region upstream of *RAB27A*, in chromosome 15 (Fig. S42). Mutations in this gene cause Griscelli syndrome, which results in pigmentary dilution in the hair and skin, as well as melanosome accumulation in melanocytes (Ménasché *et al*., 2000). Finally, we found evidence for Denisovan AI in two nearby regions in chromosome 14 (Figs. S39 and S40). One of these overlaps with *PRKCH* —encoding a protein kinase associated with cerebral infarction (Kubo *et al*., 2007). The other overlaps with *KCNH5* —coding for a potassium channel that may be associated with epileptic encephalopathy (Veeramah *et al*., 2013).

## Discussion

We have developed a new method to detect adaptive introgression along the genome using convolutional neural networks. The method has high precision when reporting candidate AI loci, and high negative predictive value when rejecting loci as not-AI: we obtain greater than 90 % accuracy under a variety of different selection scenarios (Table S4), with low false positive rates.

As reported previously (Kim *et al*., 2018; Zhang *et al*., 2020), heterosis following introgression can produce patterns very similar to AI, and we found this can inflate false positive detection of AI by our CNN. However, we simulated a DFE with recessive dominance for all mutations, which is not realistic in general, so our results in this regard represent a worse case scenario. A possible future improvement would be to train the CNN on simulations incorporating heterosis. We did not attempt this here because realistic DFE simulations represent a substantial computational burden.

The CNN took approximately 15 minutes to train on one NVIDIA Tesla T4 GPU, which amounts to 60 CPU hours for an equivalent CPU-only training procedure. All data were loaded into memory, which required approximately 120 GB RAM during training. The computational bottleneck lay in the generation of SLiM forward simulations: 300,000 simulations took approximately 80 weeks of CPU time for each of the demographic models. In the future, considerable speedups could potentially be obtained by optimising the simulation step, perhaps by implementing an adaptive introgression simulation framework that takes advantage of the backwards coalescent (e.g. building on the work by Setter *et al*., 2020).

We applied the method to human data, to look for adaptive introgression from archaic humans into the ancestors of present-day human genomes. When looking for Neanderthal AI in European genomes, we find previously found candidate genes (*BNC2*, *WRD88/GPATCH1*, *KCNQ2/EEF1A2*, *TANC1/BAZ2B*). We also recover candidates for adaptive introgression from Denisovans by applying our method to unphased Melanesian genomes. The top candidates include *TNFAIP3*, which has been reported before, but also include other, novel regions, containing genes involved in blood diseases (*CYCS*), neurological diseases (*PRKCH, KCNH5, LRNN1*), metabolism (*SFRP4, SUMF1*) and skin development (*ASCL4, RAB27A*).

We note, however, that, as with previous methods, visual inspection of the haplotypes or genotypes of the top candidate regions remains a necessary criterion to accurately assess whether a region may have been under adaptive introgression. For example, in the scans we performed, we found a few candidate regions for Neanderthal AI in Europeans that are likely to be false positives, e.g. chr2:109360001–109460000; chr4:54240001–54340000; chr8:143440001–143540000. These appear to be the result of shared ancestral variation with African populations, and yet are classified as having high probability of being under AI. Thus, our method allows for a rapid scan and prioritisation of potential targets, but these need to be further assessed for veracity and any functional consequence. Inclusion of more complex selection scenarios, involving positive or balancing selection on ancestral variation, as well as linked selection, might serve to ameliorate the rate of false positives in the future.

Furthermore, our simulation procedure does not model genotype errors or data missingness. Not explicitly accounting for this may negatively impact the robustness of the minor allele density computation and the subsequent haplotype sorting procedure, and, in turn, affect the accuracy of the CNN.

The precision of our method necessarily depends upon the demographic history of the populations involved. We found it more challenging to detect AI when the timing of gene flow is younger or the introgressing population is more diverged from the panel that is used to represent it. This is apparent when comparing results for the Neanderthal-into-European demographic scenario and the Denisovan-into-Melanesian demographic scenario. In the former, gene flow is more recent (55 kya versus 50 kya and 30 kya) (Sankararaman *et al*., 2016; Jacobs *et al*., 2019) and sequences are available for a population closely related to the putative source, which increases power. Furthermore, for the two putative pulses of Denisovan gene flow (Jacobs *et al*., 2019), we find our model has greater precision with AI for the more ancient pulse (94 % versus 83.6 %; Fig. S10), likely because haplotypes from the older pulse have more time to rise in frequency. We also found that distinguishing AI from a selective sweep (hard or soft), is relatively easier than distinguishing AI from neutral variation, and that the time of onset of selection in an AI scenario has little bearing on accuracy unless the onset is very recent.

Our method requires sequencing data from the population from which the introgression event originated. This may be problematic in cases where the source of introgression may be distantly related to the population genomic panel that is used to represent it. Future work could involve developing a CNN that can detect adaptive introgression from a ghost (unsampled) population, for cases in which genomic data from the source is unavailable (e.g. see Setter *et al*., 2020).

The method can take either phased or unphased data as input. This flexibility allows for its application to a range of study systems in the future, in which phasing may not be financially or methodologically feasible. It does, however, require called genotypes and is therefore not yet suitable for genomes sequenced at low coverage. One could envision extending the framework developed here to low-coverage genomes by working with matrices of genotype likelihoods (Korneliussen *et al*., 2014) rather than matrices of genotypes or haplotypes. A CNN could learn the relationship between the observed likelihoods under a given model and the model parameters that generated those likelihoods, but we leave that to a future work.

Future studies could also address the fact that we must use simulations to train the network, which involves an implicit amount of supervision by the user. The range of parameters and models that are simulated during training are necessarily hand-selected a priori, and misspecification does negatively affect CNN performance. Progress in this regard could involve the use of generative adversarial networks (GANs), which appears to be a fruitful way to address this. Indeed, recent work suggests that one can train a GAN to learn to generate realistic population genomic data for any population (Wang *et al*., 2020).

The attention analyses performed here allowed only a posteriori reasoning on how the network learned to predict AI, so further work is encouraged in this area. For instance, interpretability of neural networks can be assessed using symbolic metamodelling (Alaa & van der Schaar, 2019) with reinforcement learning algorithms deployed to identify the subset of most informative features of input data (Yoon *et al*., 2019). In this context, such approaches should be able to pinpoint the important characteristics of genomic data, and possibly derive more informative summary statistics to predict complex evolutionary events.

In summary, we have shown that CNNs are a powerful approach to detecting adaptive introgression and can recover both known and novel selection candidates that were introduced via admixture. As in previous applications to other problems in the field (Sheehan & Song, 2016; Flagel *et al*., 2018; Schrider & Kern, 2018; Villanea & Schraiber, 2019; Mondal *et al*., 2019; Torada *et al*., 2019; Isildak *et al*., 2020), this exemplifies how deep learning can serve as a very powerful tool for population genetic inference. This type of technique may thus be a useful resource for future studies aiming to unravel our past history and that of other species, as statistical methodologies and computational resources continue to improve.

## Methods

### Simulations

For CNN training, we performed simulations under three scenarios: neutral mutations only; positive selection of a de novo mutation in the recipient population (selective sweep); and positive selection of a derived mutation that was transferred via gene flow from the donor population to the recipient population (adaptive introgression, AI). In the sweep and AI scenarios, the selection coefficient was drawn log-uniformly from between 0.0001 and 0.1 for Europeans and between 0.001 and 0.1 for Melanesians. The uniformly distributed time of mutation was decoupled from the uniformly distributed time of selection onset (thus allowing for soft sweeps). For the selective sweep scenario, the mutation and selection times could occur at any time older than 1 kya but more recent than the split between the recipient population and its unadmixed sister population, with the constraint that the mutation must be introduced before the onset of selection. For the AI scenario, a neutrally evolving mutation was introduced to the donor population any time more recent than the split between the donor and the ancestor of recipient and unadmixed sister population, but older than 1 kya before the introgression event. Then, this mutation was transmitted to the recipient population, whereupon selection could start to act on it at any time after introgression but before 1 kya.

We further evaluated our Demographic Model A CNNs using an additional 10,000 simulations that incorporated a DFE using the parameters estimated for Europeans in Kim *et al*. (2017) and used in Kim *et al*. (2018). We considered two mutation types: 30 % neutral and 70 % deleterious. The deleterious portion of introduced mutations had a selection coefficient drawn from a reflected gamma distribution with shape parameter 0.186, and expected value −0.01314833. We approximated the dominance scheme from Kim *et al*. (2018), using a fixed dominance coefficient for deleterious mutations of 0.5*/*(1 7071.07 *E*[*s*]) where *E*[*s*] is the expected value from the gamma distribution (i.e. all deleterious mutations were effectively recessive).

To incorporate selection, we implemented a new module in stdpopsim (Adrion *et al*., 2020a), which leverages the forwards-in-time simulator SLiM (Haller & Messer, 2019) for simulating selection. For consistency, we also used stdpopsim’s SLiM engine for neutral simulations. stdpopsim uses SLiM’s ability to output tree sequences (Haller *et al*., 2019; Kelleher *et al*., 2018), which retains complete information about the samples’ marginal genealogies. Further, stdpopsim recapitates the tree sequences (ensuring that all sampled lineages have a single common ancestor), and applies neutrally evolving mutations to the genealogies, using the coalescent framework of msprime (Kelleher *et al*., 2016).

We simulated 100 kbp regions, with a mutation rate of 1.29 10*^−^*^08^ per site per generation (Tian *et al*., 2019), an empirical recombination map drawn uniformly at random from the HapMapII genetic map (Frazer *et al*., 2007), and the selected mutation introduced at the region’s midpoint. For both the sweep scenario and the AI scenario, we used a rejection-sampling approach to condition on the selected allele’s frequency being ≥ 1% in the recipient population at the end of the simulation. This was done by saving the simulation state prior to the introduction of the selected mutation (and saving again after successful transmission to the recipient population, for the AI scenario), then restoring simulations to the most recent save point if the mutation was lost, or if the allele frequency threshold was not met at the end of the simulation.

To speed up simulations, we applied a scaling factor of *Q* = 10. Scaling divides population sizes (*N*) and event times (*T*) by *Q*, and multiplies the mutation rate *µ*, recombination rate *r* and selection coefficient *s* by *Q*, such that the population genetic parameters *θ* = 4*Nµ*, *ρ* = 4*Nr*, and *Ns* remain approximately invariant to the applied scaling factor (Haller & Messer, 2019). After simulating, we further filtered our AI scenario simulations to exclude those that ended with a minor beneficial allele frequency less than a specific cutoff. We tried two cutoffs—5% and 25%—and present results for both. Rejection sampling within SLiM was not possible at these higher thresholds, as simulations often had low probability of reaching the threshold, particularly for recently introduced mutations.

To investigate Neanderthal gene flow into Europeans, we simulated an out-of-Africa demographic model with a single pulse of Neanderthal gene flow into Europeans but not into African Yoruba (Fig. 1A), using a composite of previously published model parameters (Table S1). The number of samples to simulate for each population was chosen to match the YRI and CEU panels in the 1000 Genomes dataset (The 1000 Genomes Project Consortium, 2015), and the two high coverage Neanderthal genomes (Prüfer *et al*., 2014, 2017). The two simulated Neanderthals were sampled at times corresponding to the estimated ages of the samples as reported in Prüfer *et al*. (2017).

To investigate Denisovan gene flow into Melanesian populations, we simulated an out-of-Africa demographic history incorporating two pulses of Denisovan gene flow (Malaspinas *et al*., 2016; Jacobs *et al*., 2019) implemented as the PapuansOutOfAfrica_10J19 model in stdpopsim (Adrion *et al*., 2020a). For this demographic model we sampled a single Denisovan and a single Neanderthal (with sampling time of the latter corresponding to the Altai Neanderthal’s estimated age). The number of Melanesian samples was chosen to match a subset of the IGDP panel (Jacobs *et al*., 2019). The Baining population of New Britain was excluded at the request of the IGDP data access committee, and we also excluded first degree relatives, resulting in a total of 139 Melanesian individuals used in the analysis. As this demographic model includes two pulses of Denisovan admixture, we simulated half of our AI simulations to correspond with gene flow from the first pulse, and half from the second pulse.

### Conversion of simulations to genotype matrices

We converted the tree sequence files from the simulations into genotype matrices using the tskit Python API (Kelleher *et al*., 2016). Major alleles (those with sample frequency greater than 0.5 after merging all individuals) were encoded in the matrix as 0, while minor alleles were encoded as 1. In the event of equal counts for both alleles, the major allele was chosen at random. Only sites with a minor allele frequency *>* 5% were retained. For sweep and AI simulations, we excluded the site of the selected mutation.

We note that different simulations result in different numbers of segregating sites, but a requirement for CNN training is that each datum in a batch must have the same dimensions. Existing approaches to solve this problem are to use only a fixed number of segregating sites (Chan *et al*., 2018), to pad the matrix out to the maximum number of observed segregating sites (Flagel *et al*., 2018), or to use an image-resize function to constrain the size of the input data (Torada *et al*., 2019). Each approach discards spatial information about the local density of segregating sites, although this may be recovered by including an additional vector of inter-site distances as input to the network (Flagel *et al*., 2018).

To obtain the benefits of image resizing (fast training times for reduced sizes and easy application to genomic windows of a fixed size), while avoiding its drawbacks, we chose to resize our input matrices differently, and only along the dimension corresponding to sites. To resize the genomic window to have length *m*, the window was partitioned into *m* bins, and for each individual haplotype we counted the number of minor alleles observed per bin. Compared with interpolation-based resizing (Torada *et al*., 2019), binning is qualitatively similar, but preserves inter-allele distances and thus the local density of segregating sites. Furthermore, as we do not resize along the dimension corresponding to individuals, this also permits the use of permutation-invariant networks (Chan *et al*., 2018), although we do not pursue that network architecture here. We report results for *m* = 256, but also tried *m* = 32, 64, and 128 bins. Preliminary results indicated greater training and validation accuracy for CNNs trained with more bins, around 1% difference between both 32 and 64, and 64 and 128, although only marginal improvement for 256 compared with 128 bins. When matching unphased data, we combined genotypes by summing minor allele counts between the chromosomes of each individual. We note that all data were treated as either phased, or unphased, and no mixed phasing was considered.

We then partitioned the resized genotype matrix into submatrices by population. Submatrices were ordered left-to-right according to the donor, recipient, and unadmixed populations respectively. For genotype matrices including both Neanderthals and Denisovans, we placed the non-donor archaic population to the left of the donor. To ensure that a non-permutation-invariant CNN could learn the structure in our data, we sorted the haplotypes (Flagel *et al*., 2018; Torada *et al*., 2019). The resized haplotypes/individuals within each submatrix were ordered left-to-right by decreasing similarity to the donor population, calculated as the Euclidean distance to the average minor-allele density of the donor population (analogous to a vector of the donor allele frequencies). An example (phased) genotype matrix image for an AI simulation is shown in Fig. 1B.

### Conversion of empirical data to genotype matrices

Using bcftools (Li, 2011), we performed a locus-wise intersection of the following VCFs: 1000 Genomes (The 1000 Genomes Project Consortium, 2015), IGDP (Jacobs *et al*., 2019), the high coverage Denisovan genome (Meyer *et al*., 2012), and the Altai and Vindija Neanderthal genomes (Prüfer *et al*., 2014, 2017). All VCFs corresponded to the GRCh37/hg19 reference sequence. Genotype matrices were constructed by parsing the output of bcftools query over 100 kbp windows, filtering out sites with sample allele frequency < 5% or with more than 10 % of genotypes missing, then excluding windows with fewer than 20 segregating sites. Each genotype matrix was then resized and sorted as described for simulations. When data were considered to be phased, as for the CEU/YRI populations, we also treated the Neanderthal genotypes as if they were phased according to REF/ALT columns in the VCF. While this is equivalent to random phasing, both high-coverage Neanderthal individuals are highly inbred, so this is unlikely to be problematic in practice.

### CNN model architecture and training

We implemented the CNN model in Keras (Chollet *et al*., 2015), configured to use the Tensorflow backend (Abadi *et al*., 2015). To save disk space and memory, the input matrices were stored as 8 bit integers rather than floating point numbers, and were not mean-centred or otherwise normalised prior to input into the network. We instead made the first layer of our network a batch normalisation layer, which converts the input layer to floating point numbers and learns the best normalisation of the data for the network.

The CNN architecture (Fig. 1C) consists of *k* convolution blocks each comprised of a batch normalisation layer followed by a 2D convolution layer with 2×2 stride, 16 filters of size 4×4, and leaky ReLU activation. The *k* blocks are followed by a single fully-connected output node of size one, with sigmoid activation. We do not include pooling layers, as is common in a CNN architecture (e.g. Torada *et al*., 2019), and instead use a 2×2 stride size to reduce the output size of successive blocks (Springenberg *et al*., 2015). This is computationally cheaper and had no observable difference in network performance. We sought to maximise the depth of the network, but the size of the input layer constrains the maximum number of blocks in the network due to successive halving of the dimensionality in each block. For *m* = 256 resizing bins, we used *k* = 7 blocks.

We partitioned 100,000 independent simulations for each of the three selection scenarios into training and validation sets (approximate 90%/10% split). The model was trained for three epochs, with model weights updated after batches of 64, using the Adam optimiser and cross-entropy for the loss function. We evaluated model fit by inspecting loss and accuracy terms at end of training (Table S4). Preliminary analyses indicated three epochs were sufficient for approximate convergence between training and validation metrics, but we did not observe divergence (likely indicating overfitting) even when training for additional epochs.

### Calibration

CNNs may produce improperly calibrated probabilities (Guo *et al*., 2017). For a well calibrated output, we expect proportion *x* of the output probabilities with Pr[AI] ∼ *x* to be true positives. To calibrate our CNN output, we applied beta calibration (Kull *et al*., 2017) by fitting a logistic regression model to our validation data after model training. Beta and other calibration methods were assessed by fitting the calibrator to the training dataset and inspecting reliability plots on a validation dataset (Figs. S2 to S5). We also checked if the sum of the residuals was normally distributed, following the approach of Turner *et al*. (2019). Both beta calibration and isotonic regression gave well-calibrated probabilities compared with uncalibrated model outputs, and we chose to apply beta calibration due to its relative simplicity (Kull *et al*., 2017).

The proportion of predictions which are false positives or false negatives depends upon the relative ratios of AI versus not-AI windows of the genome. This ratio is not known, so we fitted our calibration procedure using resampled training data with multiple ratios for neutral:sweep:AI (Fig. 4).

### Saliency maps

Saliency maps were computed on, and then averaged over, a set of 100 simulated genotype matrices for each simulated scenario, using keras-vis (Kotikalapudi & contributors, 2017). We applied the visualize_saliency function on a a pre-trained CNN, and we configured it to use the guided backpropagation modifier. A sharper image was obtained by exchanging the CNN output layer’s sigmoid activation with linear activation, as recommended in the keras-vis documentation.

### Application of trained CNN to empirical datasets

We show Manhattan plots where each data point is a 100 kbp window that moves along the genome in steps of size 20 kbp. Gene annotations were extracted from the Ensembl release 87 GFF3 file (with GRCh37/hg19 coordinates), obtained via ensembl’s ftp server. We extracted the columns with source=“ensembl_havana” and type=“gene”, and report the genes which intersected with the 30 top ranking CNN predictions or a 100 kbp flanking region. Adjacent regions were merged together prior to intersection, so that genes were reported only once.

### Compute resources

All simulations and results reported here were obtained on an compute server with two Intel Xeon 6248 CPUs (80 cores total), 768 GB RAM, and five NVIDIA Tesla T4 GPUs. 300,000 SLiM simulations took approximately 80 weeks of CPU time for each of Demographic Model A and B. Each simulation executes independently, and is readily distributed across cores or compute nodes. This produced 450 GB of tree sequence files. The resized genotype matrices were compressed into a Zarr cache (Zarr Development Team, 2020) with size 2.8 GB, for faster loading. Training a single CNN on one GPU took approximately 15 minutes, or 60 CPU hours for an equivalent CPU-only training procedure. We did not attempt to optimise memory usage, and thus all data were loaded into memory, requiring approximately 120 GB RAM during training. Predicting AI for all genomic windows on an empirical dataset (22 single-chromosome BCF files) took 1 CPU hour. However, our prediction pipeline uses multiprocessing and efficiently scales to 80 cores.

### Code availability

The source code for performing simulations, training and evaluating a CNN, and applying a CNN to empirical VCF data, were developed in a new Python application called genomatnn, available at https://github.com/grahamgower/genomatnn. Python code for visualising the trained models can be found at https://github.com/pabloswfly/CNN-vis.

## Acknowledgements

We thank Andrew Kern, Martin Sikora, Flora Jay and Anders Albrechtsen, as well as members of the Racimo group and the PopSim consortium, for helpful advice and discussions. We also thank Murray Cox and Georgi Hudjashov for facilitating access to the IGDP data. FR and GG were funded by a Villum Fonden Young Investigator award to FR (project no. 00025300). MF was funded by a Leverhulme Research Project grant (RPG-2018-208).

## Supplementary Tables

**Table S1:**
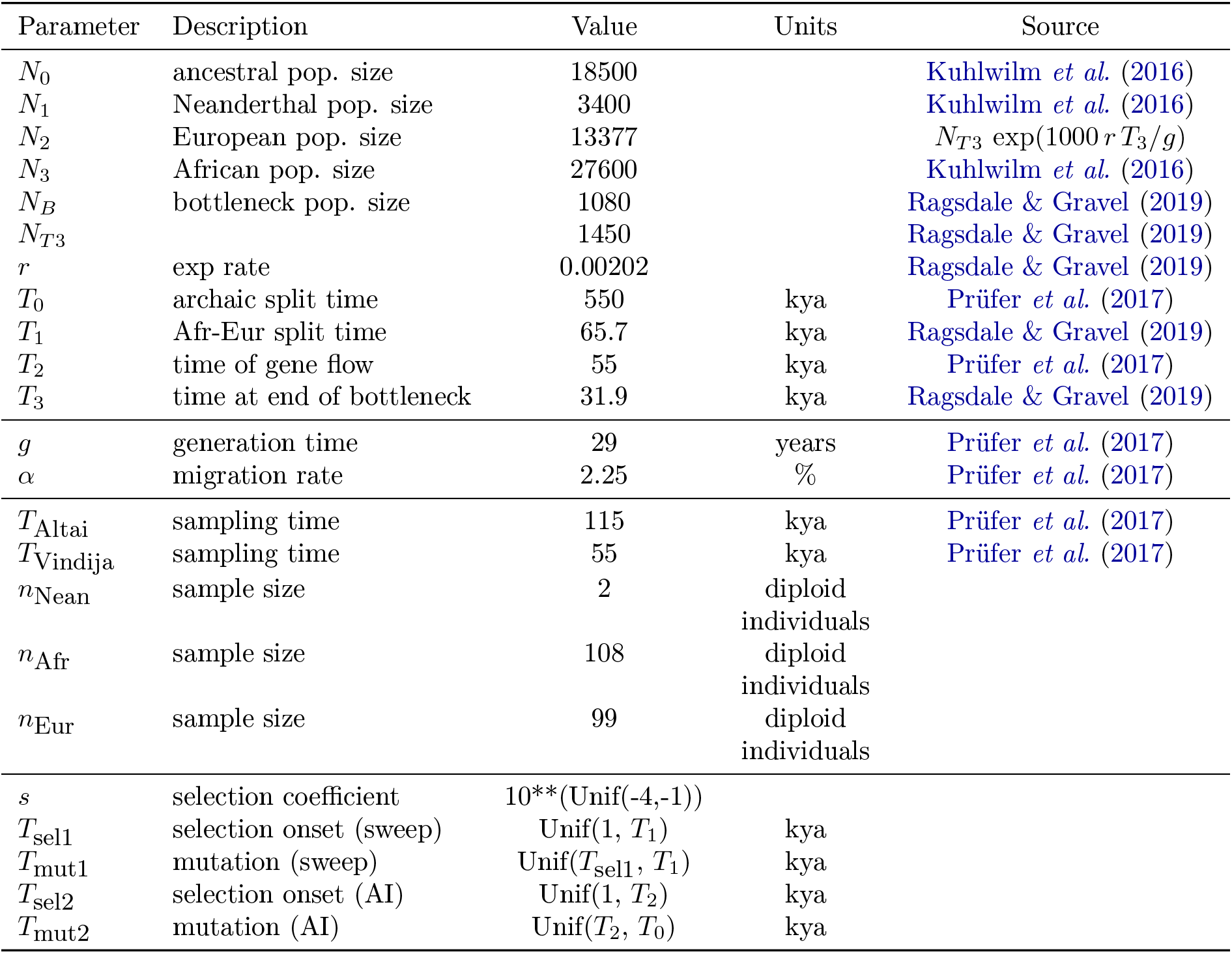
Parameter values used for simulating the HomininComposite_4G20 demographic model, with parameters corresponding to Fig. 1A.

**Table S2:**
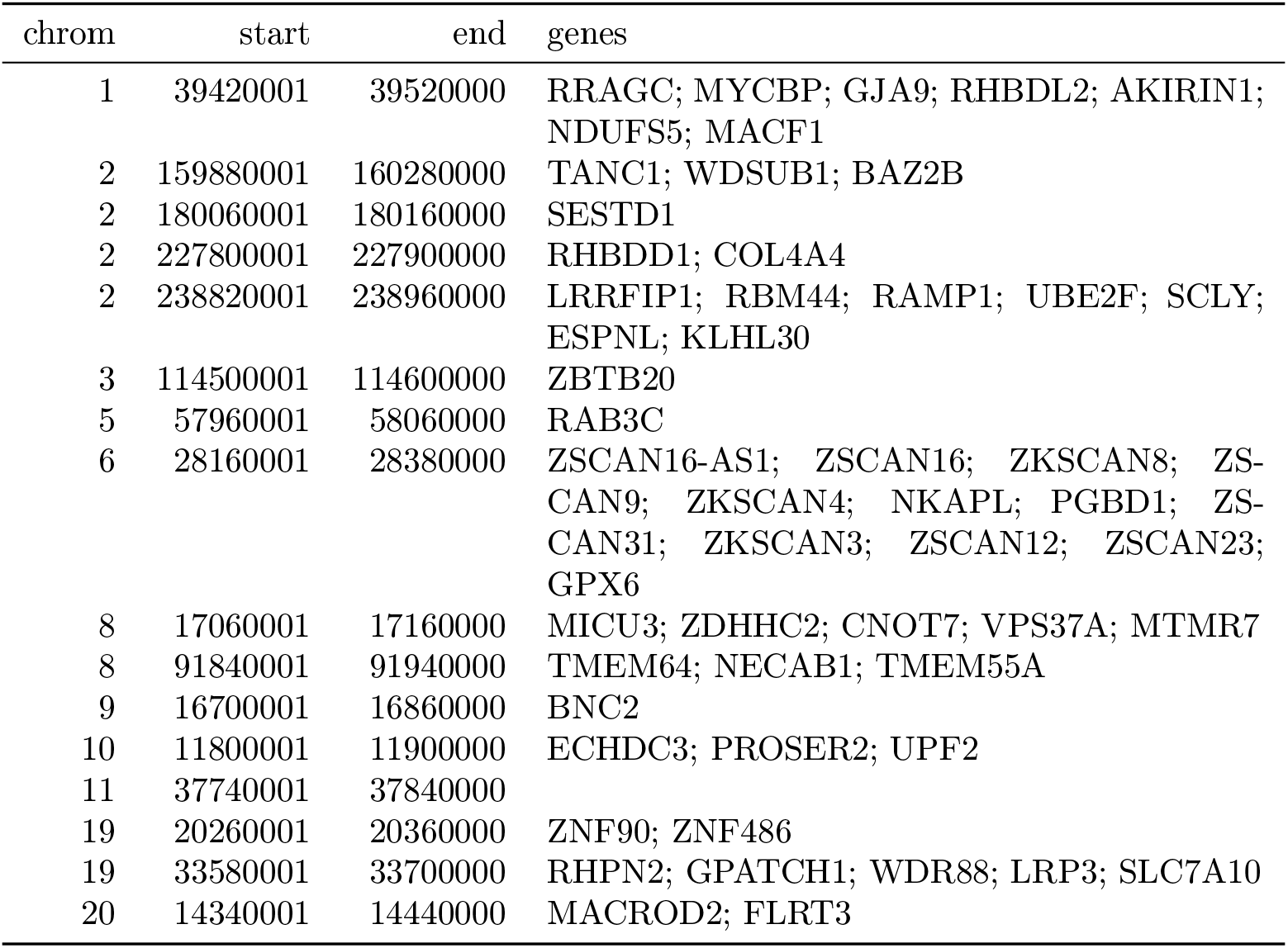
Top ranking gene candidates corresponding to Neanderthal AI in Europeans. We show genes which overlap, or are within 100 kbp of, the 30 highest ranked 100 kbp intervals. Adjacent intervals have been merged. The CNN was trained using only AI simulations with selected mutation having allele frequency > 5%, and subsequently calibrated with resampled neutral:sweep:AI ratios of 1:0.1:0.02.

| chrom | start | end | genes |
| --- | --- | --- | --- |
| 1 | 39420001 | 39520000 | RRAGC; MYCBP; GJA9; RHBDL2; AKIRIN1; NDUFS5; MACF1 |
| 2 | 159880001 | 160280000 | TANC1; WDSUB1; BAZ2B |
| 2 | 180060001 | 180160000 | SESTD1 |
| 2 | 227800001 | 227900000 | RHBDD1; COL4A4 |
| 2 | 238820001 | 238960000 | LRRFIP1; RBM44; RAMP1; UBE2F; SCLY; ESPNL; KLHL30 |
| 3 | 114500001 | 114600000 | ZBTB20 |
| 5 | 57960001 | 58060000 | RAB3C |
| 6 | 28160001 | 28380000 | ZSCAN16-AS1; ZSCAN16; ZKSCAN8; ZSCAN9; ZKSCAN4; NKAPL; PGBD1; ZSCAN31; ZKSCAN3; ZSCAN12; ZSCAN23; GPX6 |
| 8 | 17060001 | 17160000 | MICU3; ZDHHC2; CNOT7; VPS37A; MTMR7 |
| 8 | 91840001 | 91940000 | TMEM64; NECAB1; TMEM55A |
| 9 | 16700001 | 16860000 | BNC2 |
| 10 | 11800001 | 11900000 | ECHDC3; PROSER2; UPF2 |
| 11 | 37740001 | 37840000 |  |
| 19 | 20260001 | 20360000 | ZNF90; ZNF486 |
| 19 | 33580001 | 33700000 | RHPN2; GPATCH1; WDR88; LRP3; SLC7A10 |
| 20 | 14340001 | 14440000 | MACROD2; FLRT3 |

**Table S3:** Top ranking gene candidates corresponding to Denisovan AI in Melanesians. We show genes which overlap, or are within 100 kbp of, the 30 highest ranked 100 kbp intervals. Adjacent intervals have been merged. The CNN was trained using only AI simulations with selected mutation having allele frequency > 5%, and subsequently calibrated with resampled neutral:sweep:AI ratios of 1:0.1:0.02.

| chrom | start | end | genes |
| --- | --- | --- | --- |
| 1 | 2880001 | 2980000 | ACTRT2; LINC00982; PRDM16 |
| 1 | 220080001 | 220180000 | SLC30A10; EPRS; BPNT1; IARS2 |
| 2 | 221040001 | 221140000 |  |
| 3 | 15400001 | 15500000 | SH3BP5; METTL6; EAF1; COLQ |
| 4 | 41960001 | 42100000 | TMEM33; DCAF4L1; SLC30A9; BEND4 |
| 5 | 135440001 | 135540000 | TGFBI; SMAD5-AS1; SMAD5; TRPC7 |
| 6 | 81980001 | 82120000 | FAM46A |
| 7 | 121160001 | 121260000 |  |
| 9 | 95500001 | 95600000 | IPPK; BICD2; ZNF484 |
| 10 | 59660001 | 59760000 |  |
| 12 | 80780001 | 80880000 | OTOGL; PTPRQ |
| 12 | 84620001 | 84740000 |  |
| 14 | 57620001 | 57760000 | EXOC5; AP5M1; NAA30 |
| 17 | 29480001 | 29720000 | NF1; OMG; EVI2B; EVI2A; RAB11FIP4 |
| 18 | 38180001 | 38320000 |  |
| 20 | 54340001 | 54440000 |  |

**Table S4:**
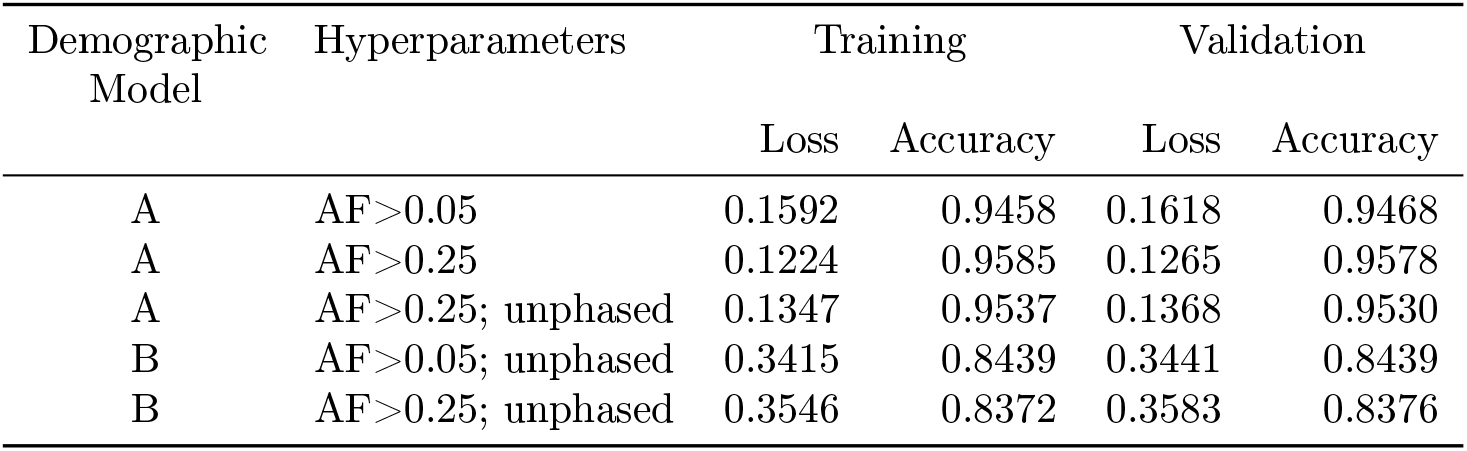
Loss and accuracy for CNNs after training for three epochs, as reported by Keras/Tensorflow, for the training and validation datasets. Binary cross-entropy was used for the loss function.

| Demographic Model | Hyperparameters | Training |  | Validation |  |
| --- | --- | --- | --- | --- | --- |
|  |  | Loss | Accuracy | Loss | Accuracy |
| A | AF$>0.05$ | 0.1592 | 0.9458 | 0.1618 | 0.9468 |
| A | AF$>0.25$ | 0.1224 | 0.9585 | 0.1265 | 0.9578 |
| A | AF$>0.25$; unphased | 0.1347 | 0.9537 | 0.1368 | 0.9530 |
| B | AF$>0.05$; unphased | 0.3415 | 0.8439 | 0.3441 | 0.8439 |
| B | AF$>0.25$; unphased | 0.3546 | 0.8372 | 0.3583 | 0.8376 |

## Supplementary Figures

**Figure S1:**
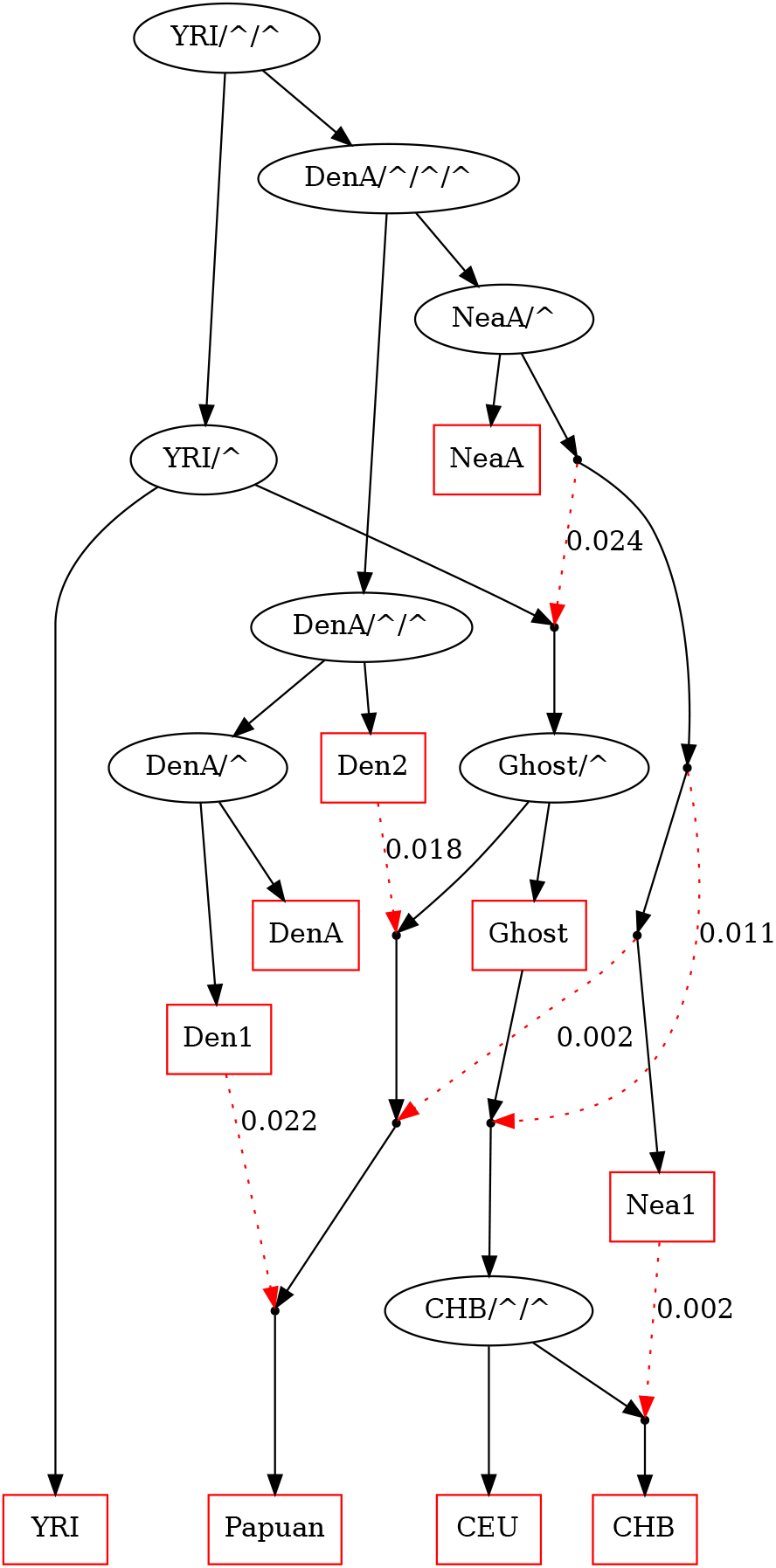
Overview of the Jacobs *et al.* (2019) demographic model, featuring two pulses of Denisovan gene flow into Papuans, which we implemented as the PapuansOutOfAfrica_10J19 model in stdpopsim. Black lines show ancestor/descendent relations and red dotted lines show pulses of admixture with the indicated proportion. DenA and NeaA are the sampled populations corresponding to Altai Denisovan and Altai Neanderthal, while Den1, Den2, and Nea1 correspond to introgressing lineages.

**Figure S2:**
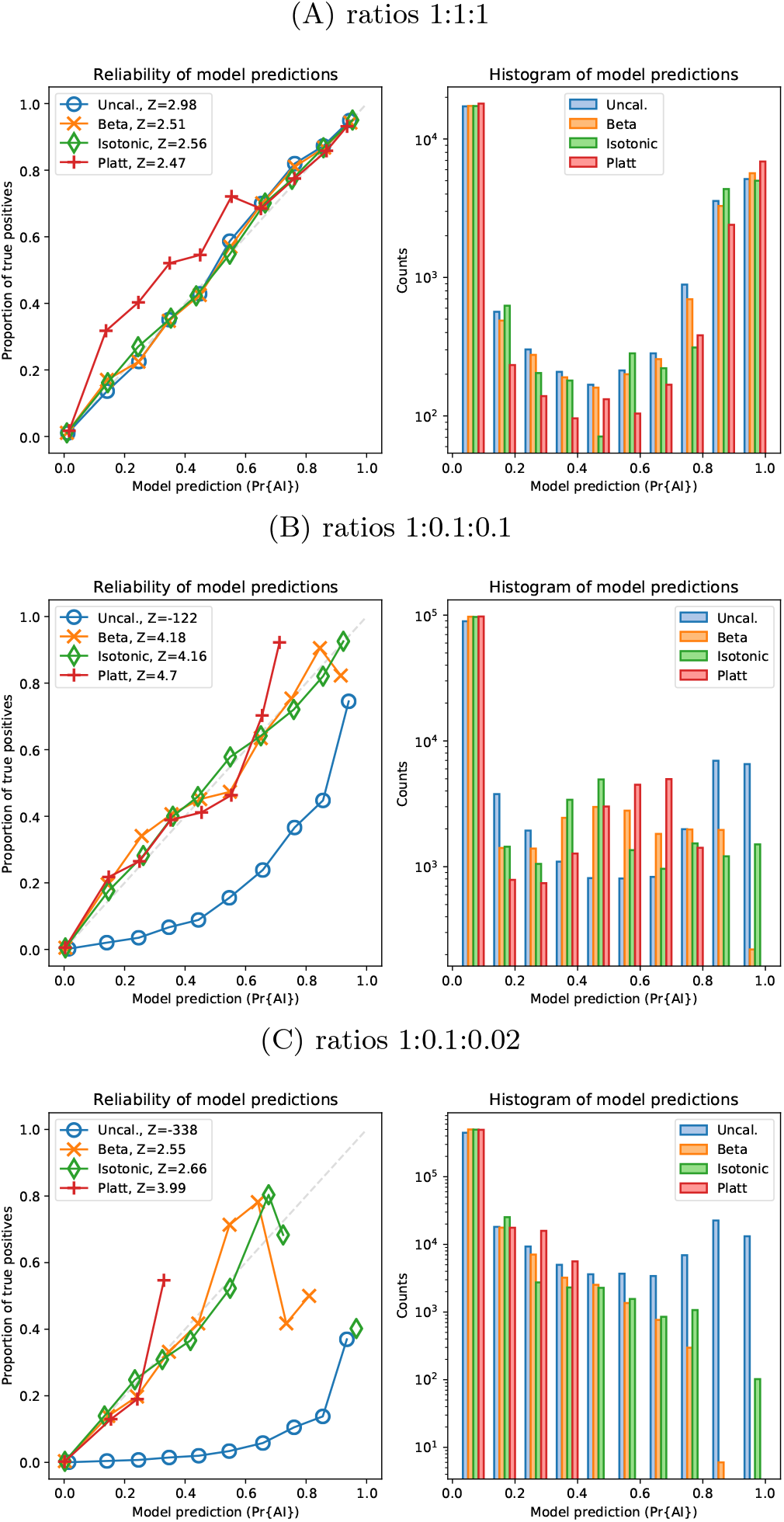
Reliability of probabilities produced by the CNN, for the validation dataset, with and without calibration, for Demographic Model A with a minimum beneficial allele frequency of 5%. The variance-normalised sum of residuals is inset in the upper left corner of each of the reliability plots (*Z*), which for well-calibrated predictions is approximately normally distributed (Turner *et al.*, 2019).

**Figure S3:**
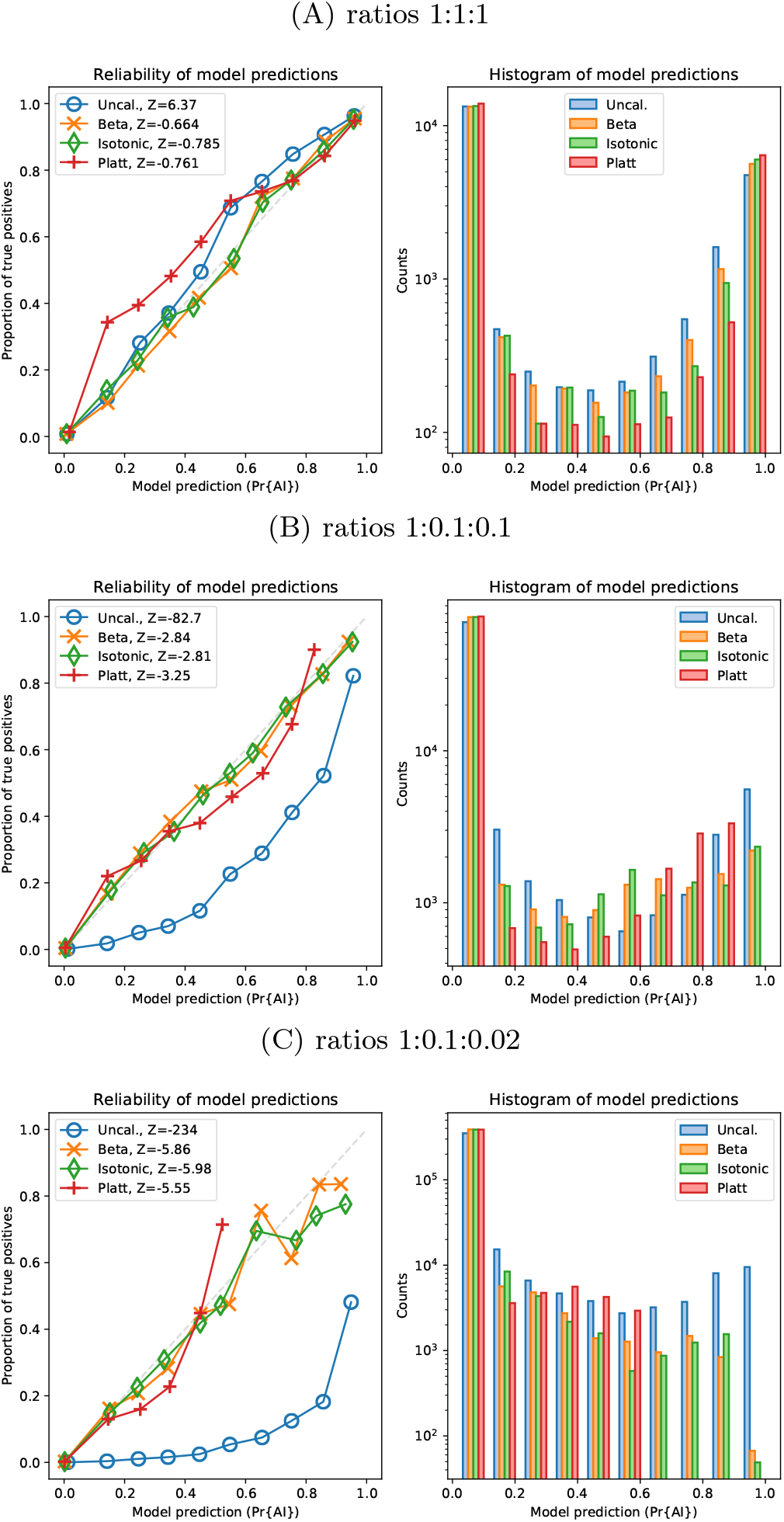
Reliability of probabilities produced by the CNN, for the validation dataset, with and without calibration, for Demographic Model A with a minimum beneficial allele frequency of 25%. The variance-normalised sum of residuals is inset in the upper left corner of each of the reliability plots (*Z*), which for well-calibrated predictions is approximately normally distributed (Turner *et al.*, 2019).

**Figure S4:**
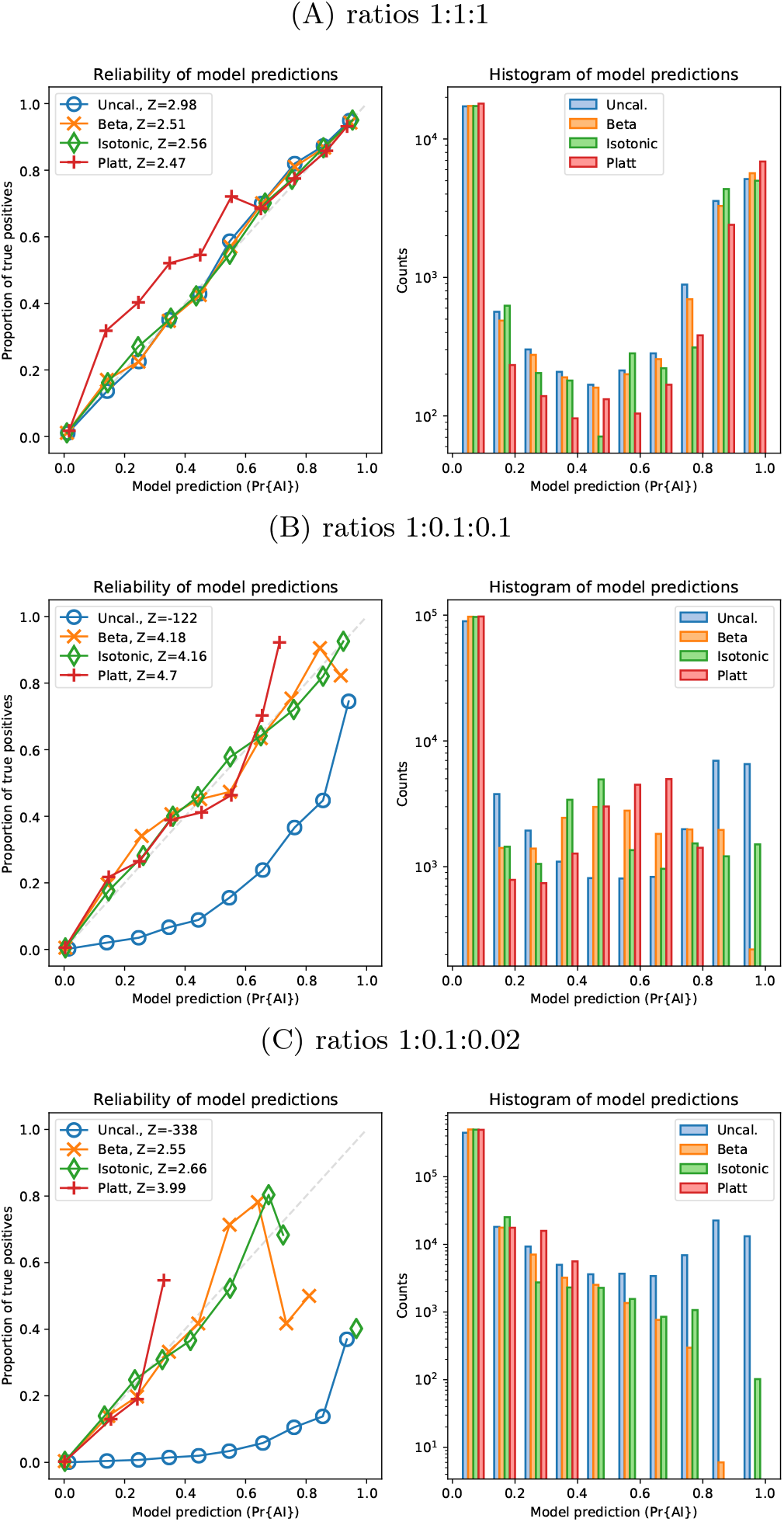
Reliability of probabilities produced by the CNN, for the validation dataset, with and without calibration, for Demographic Model B with a minimum beneficial allele frequency of 5%. The variance-normalised sum of residuals is inset in the upper left corner of each of the reliability plots (*Z*), which for well-calibrated predictions is approximately normally distributed (Turner *et al.*, 2019).

**Figure S5:**
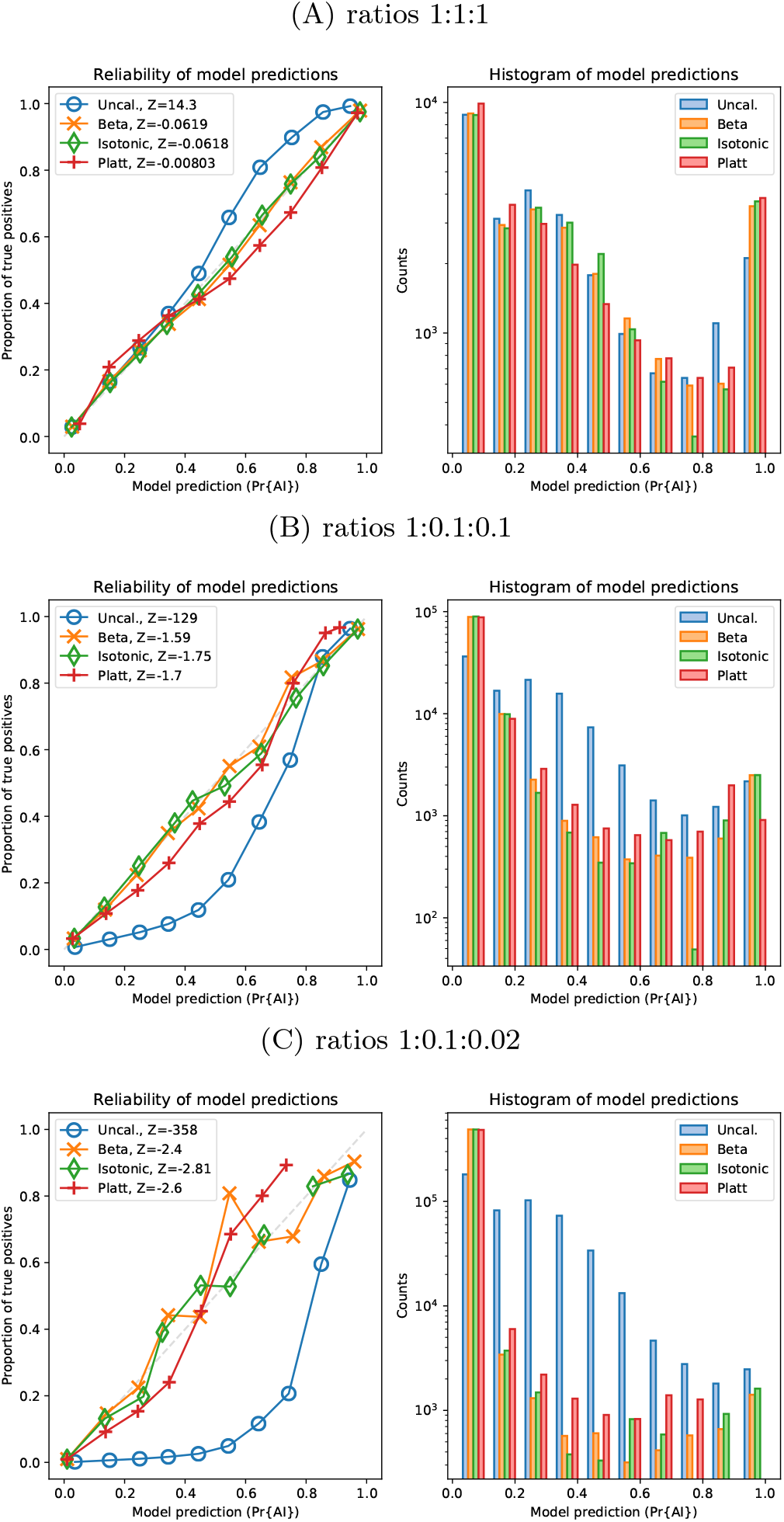
Reliability of probabilities produced by the CNN, for the validation dataset, with and without calibration, for Demographic Model B with a minimum beneficial allele frequency of 25%. The variance-normalised sum of residuals is inset in the upper left corner of each of the reliability plots (*Z*), which for well-calibrated predictions is approximately normally distributed (Turner *et al.*, 2019).

**Figure S6:**
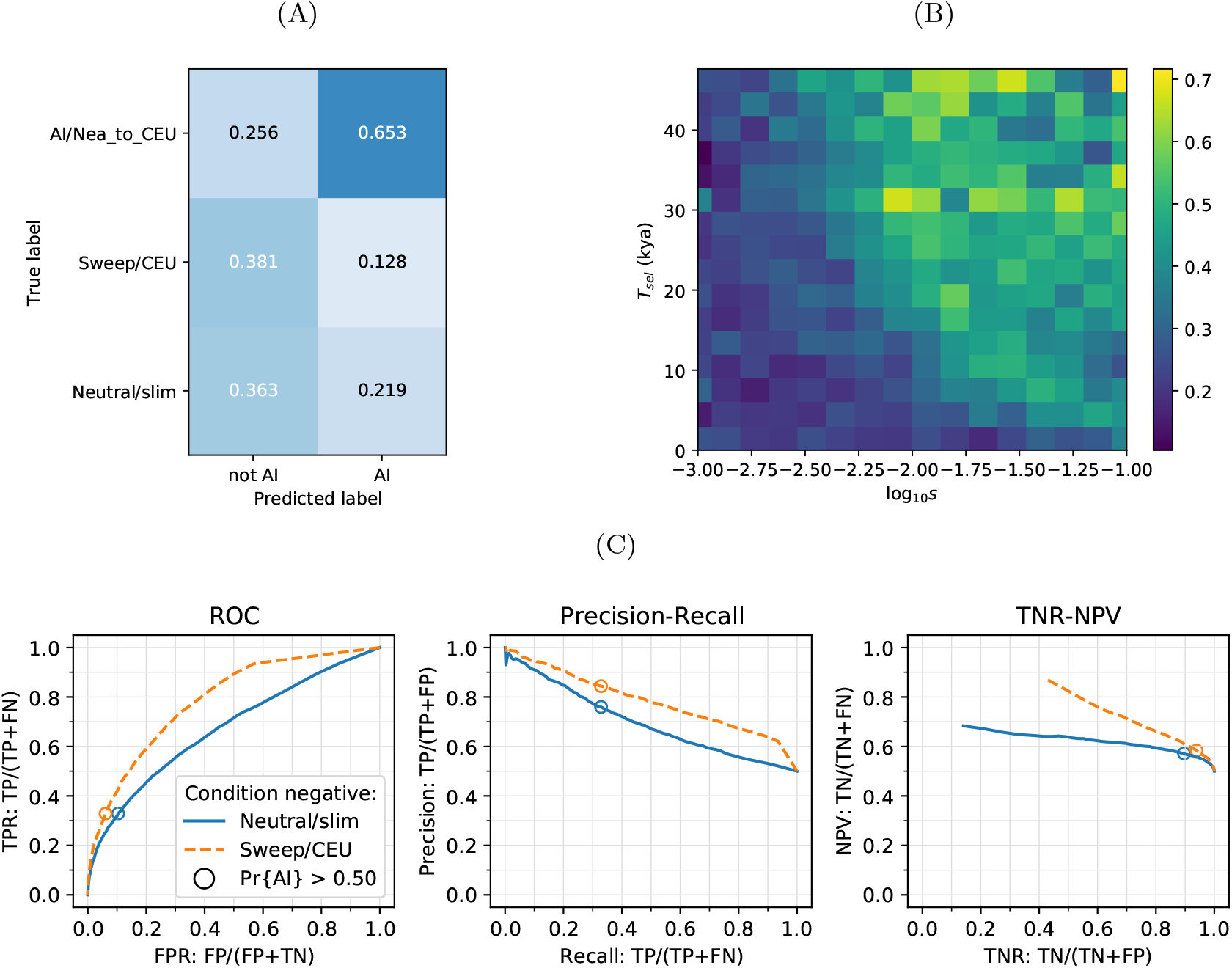
CNN performance on validation simulations for Demographic Model B, after training using Demographic Model A. The CNN was trained using only AI simulations with selected mutation having allele frequency > 0.25. A) Confusion matrix. For the two prediction categories, either “not AI” or AI, we show the proportion attributed to each of the true (simulated) scenarios. B) Average CNN prediction for AI scenarios, binned by selection coefficient, *s*, and time of onset of selection *T_sel_*. C) ROC curves, precision recall curves and True Negative Rate vs. Negative Predictive Value (TNR-NPV) curves. The positive condition is AI. The negative conditions are shown using different line styles/colours. The circles indicate the point in ROC-space (respectively Precision-Recall-space, and TNR-NPV-space) when using the threshold Pr[AI] > 0.5 for classifying a genotype matrix as AI. TP: true positives; FP: false positives; TPR: true positive rate; FPR: false positive rate; ROC: Receiver operating characteristics; TNR: true negative rate; TPR: true positive rate.

**Figure S7:**
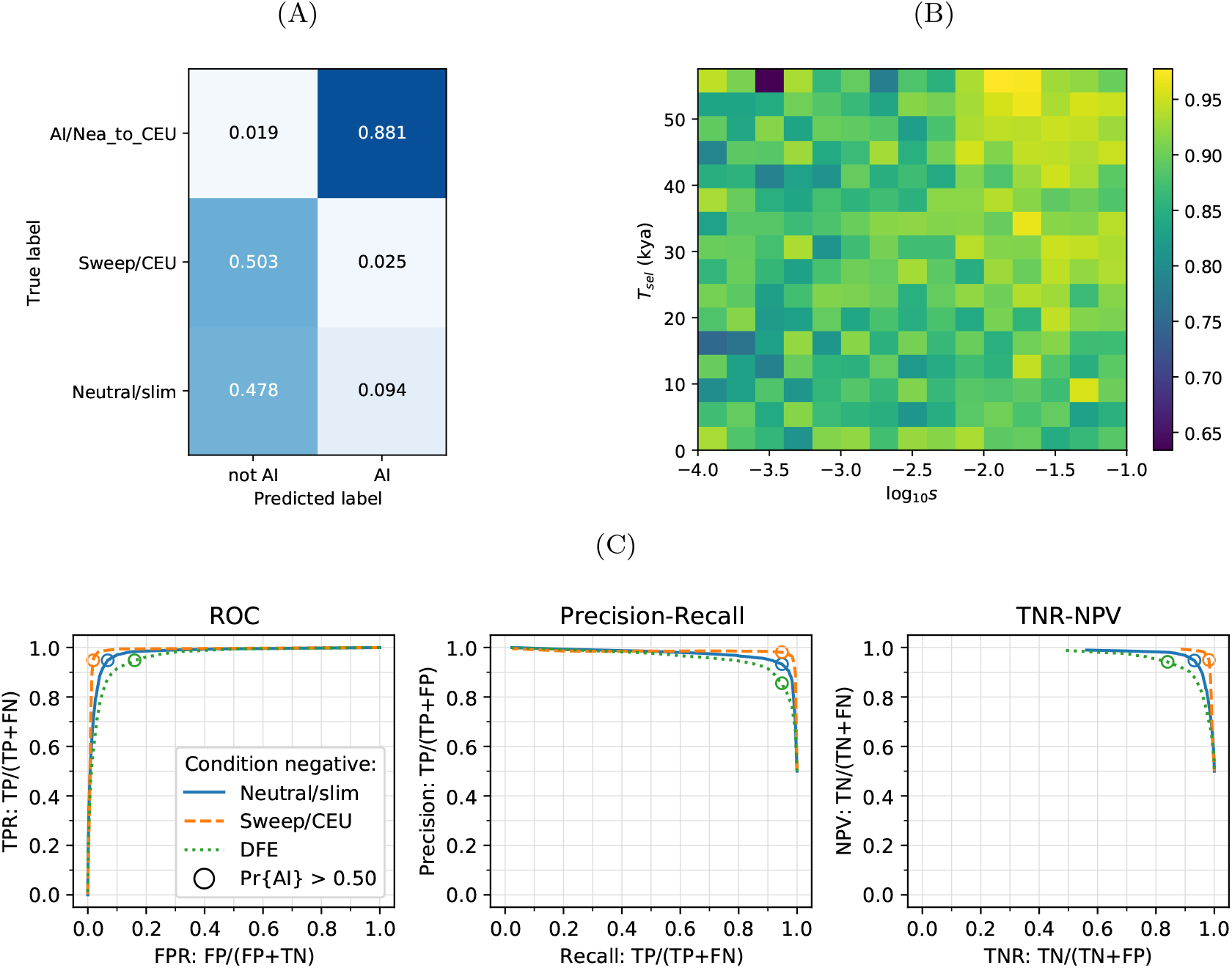
CNN performance on validation simulations for Demographic Model A with unphased data. The CNN was trained using only AI simulations with selected mutation having allele frequency > 25%. A) Confusion matrix. For the two prediction categories, either “not AI” or AI, we show the proportion attributed to each of the true (simulated) scenarios. B) Average CNN prediction for AI scenarios, binned by selection coefficient, *s*, and time of onset of selection *T_sel_*. C) ROC curves, precision recall curves and True Negative Rate vs. Negative Predictive Value (TNR-NPV) curves. The positive condition is AI. The negative conditions are shown using different line styles/colours. The circles indicate the point in ROC-space (respectively Precision-Recall-space, and TNR-NPV-space) when using the threshold Pr[AI] > 0.5 for classifying a genotype matrix as AI. DFE: distribution of fitness effects. TP: true positives; FP: false positives; TPR: true positive rate; FPR: false positive rate; ROC: Receiver operating characteristics; TNR: true negative rate; TPR: true positive rate.

**Figure S8:**
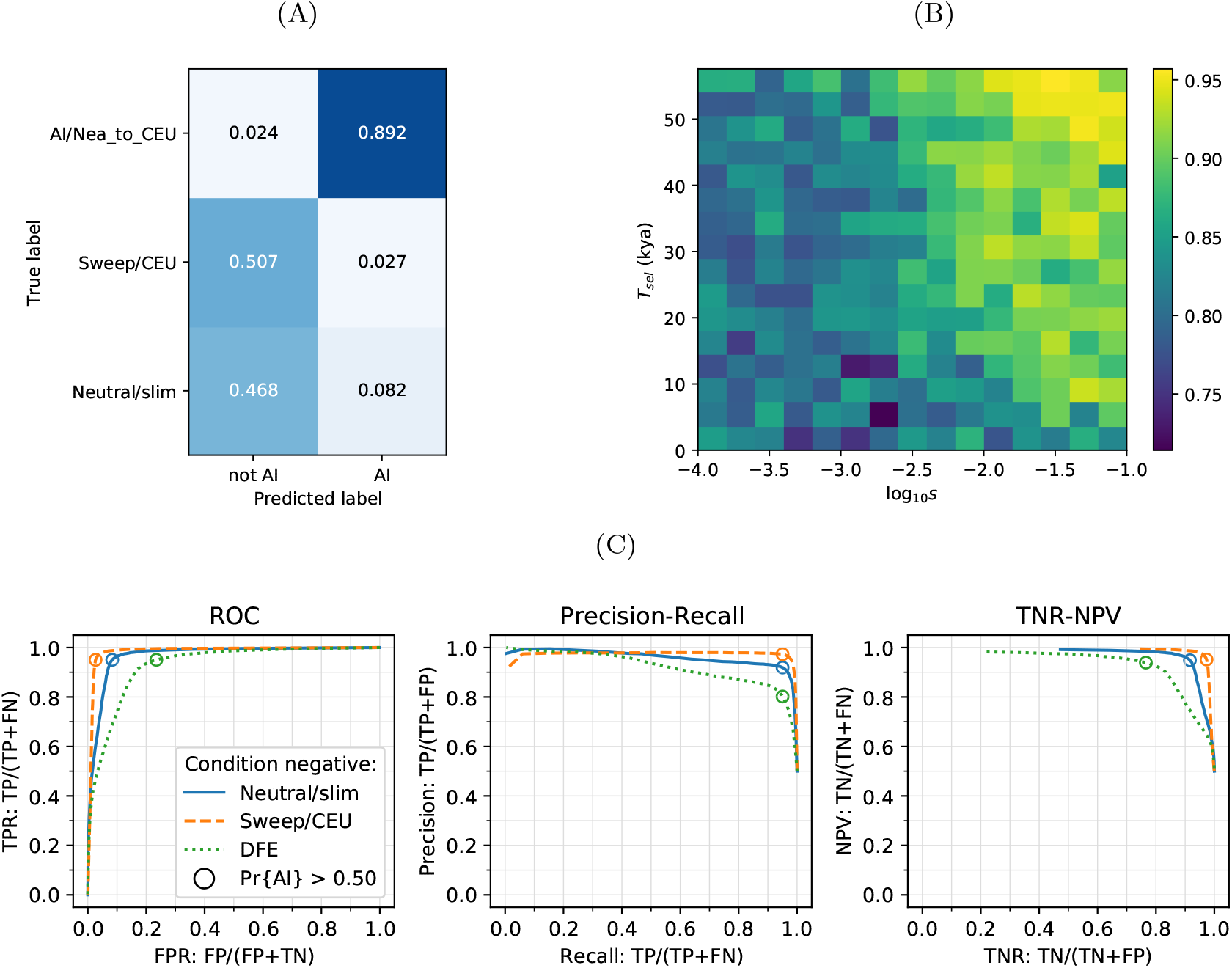
CNN performance on validation simulations for Demographic Model A with phased data. The CNN was trained using only AI simulations with selected mutation having allele frequency > 5%. A) Confusion matrix. For the two prediction categories, either “not AI” or AI, we show the proportion attributed to each of the true (simulated) scenarios. B) Average CNN prediction for AI scenarios, binned by selection coefficient, *s*, and time of onset of selection *T_sel_*. C) ROC curves, precision recall curves and True Negative Rate vs. Negative Predictive Value (TNR-NPV) curves. The positive condition is AI. The negative conditions are shown using different line styles/colours. The circles indicate the point in ROC-space (respectively Precision-Recall-space, and TNR-NPV-space) when using the threshold Pr[AI] > 0.5 for classifying a genotype matrix as AI. DFE: distribution of fitness effects. TP: true positives; FP: false positives; TPR: true positive rate; FPR: false positive rate; ROC: Receiver operating characteristics; TNR: true negative rate; TPR: true positive rate.

**Figure S9:**
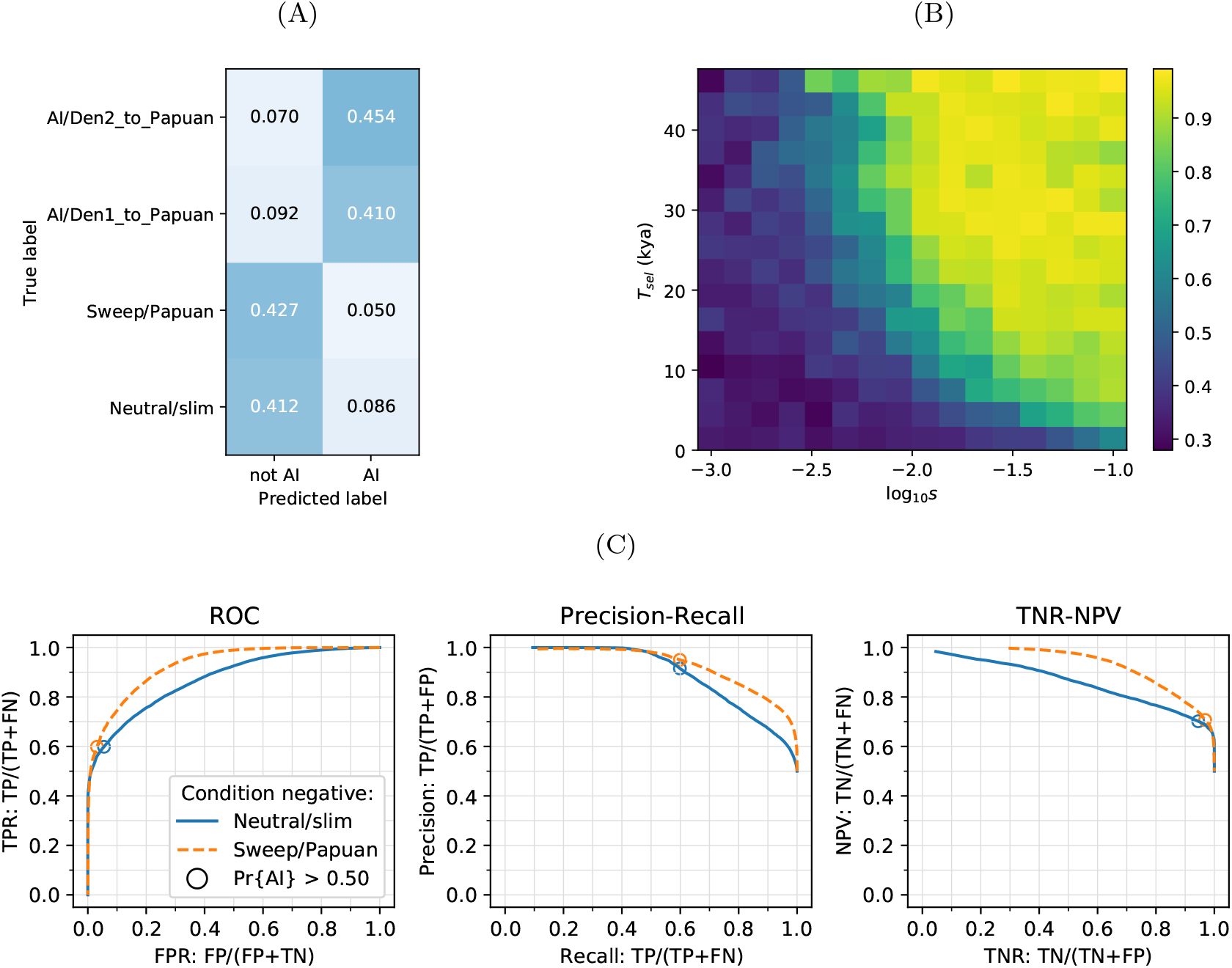
CNN performance on validation simulations for Demographic Model B with unphased data. The CNN was trained using only AI simulations with selected mutation having allele frequency > 5%. A) Confusion matrix. For the two prediction categories, either “not AI” or AI, we show the proportion attributed to each of the true (simulated) scenarios. B) Average CNN prediction for AI scenarios, binned by selection coefficient, *s*, and time of onset of selection *T_sel_*. C) ROC curves, precision recall curves and True Negative Rate vs. Negative Predictive Value (TNR-NPV) curves. The positive condition is AI. The negative conditions are shown using different line styles/colours. The circles indicate the point in ROC-space (respectively Precision-Recall-space, and TNR-NPV-space) when using the threshold Pr[AI] > 0.5 for classifying a genotype matrix as AI. TP: true positives; FP: false positives; TPR: true positive rate; FPR: false positive rate; ROC: Receiver operating characteristics; TNR: true negative rate; TPR: true positive rate.

**Figure S10:**
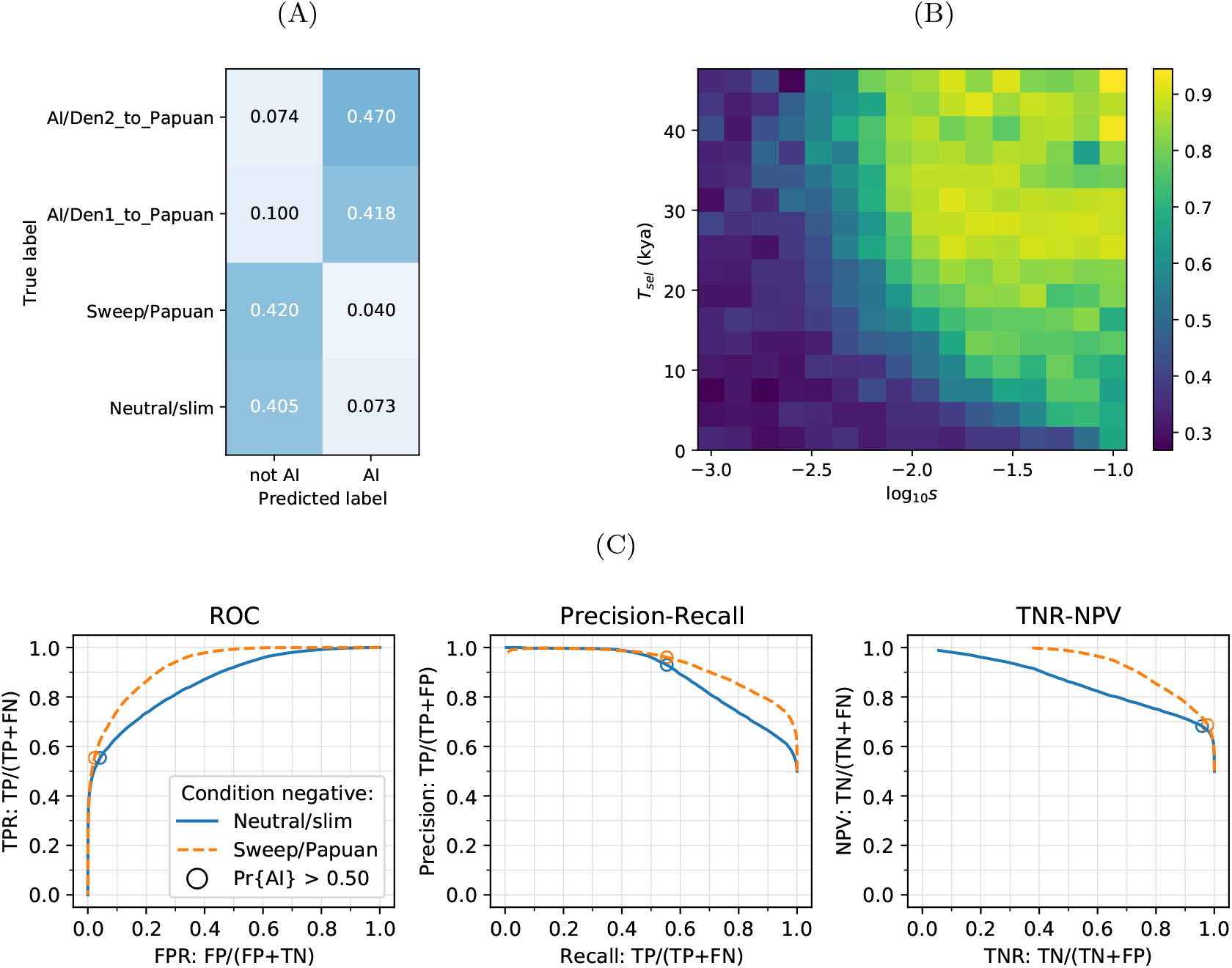
CNN performance on validation simulations for Demographic Model B with unphased data. The CNN was trained using only AI simulations with selected mutation having allele frequency > 25%. A) Confusion matrix. For the two prediction categories, either “not AI” or AI, we show the proportion attributed to each of the true (simulated) scenarios. B) Average CNN prediction for AI scenarios, binned by selection coefficient, *s*, and time of onset of selection *T_sel_*. C) ROC curves, precision recall curves and True Negative Rate vs. Negative Predictive Value (TNR-NPV) curves. The positive condition is AI. The negative conditions are shown using different line styles/colours. The circles indicate the point in ROC-space (respectively Precision-Recall-space, and TNR-NPV-space) when using the threshold Pr[AI] > 0.5 for classifying a genotype matrix as AI. TP: true positives; FP: false positives; TPR: true positive rate; FPR: false positive rate; ROC: Receiver operating characteristics; TNR: true negative rate; TPR: true positive rate.

**Figure S11:**
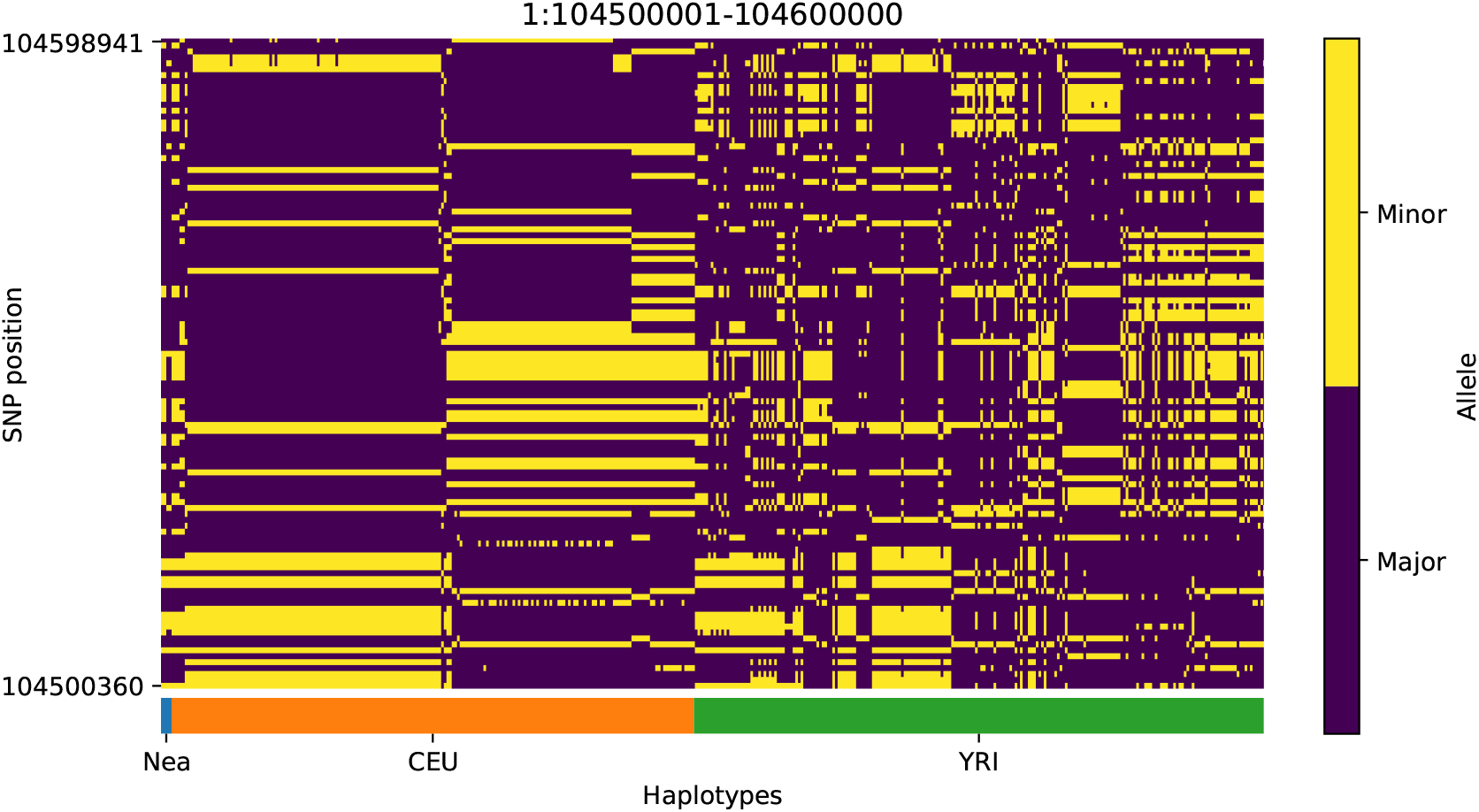
Haplotype plot for the candidate region chr1:104500001-104600000 in the Neanderthal-into-European AI scan. Bright yellow indicates minor allele, dark blue indicates major allele. Haplotypes within populations are sorted left-to-right by similarity to Neanderthals.

**Figure S12:**
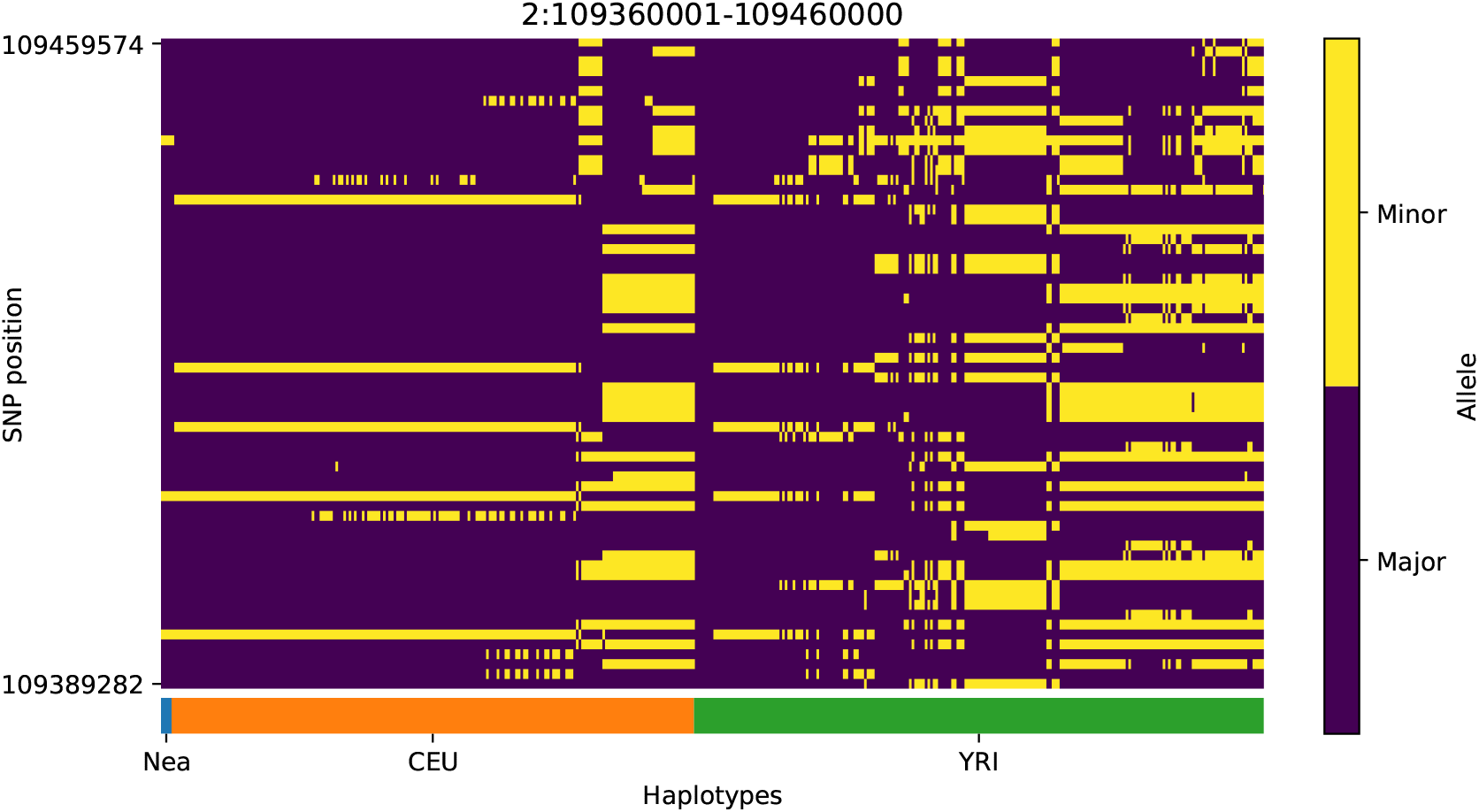
Haplotype plot for the candidate region chr2:109360001-109460000 in the Neanderthal-into-European AI scan. Bright yellow indicates minor allele, dark blue indicates major allele. Haplotypes within populations are sorted left-to-right by similarity to Neanderthals.

**Figure S13:**
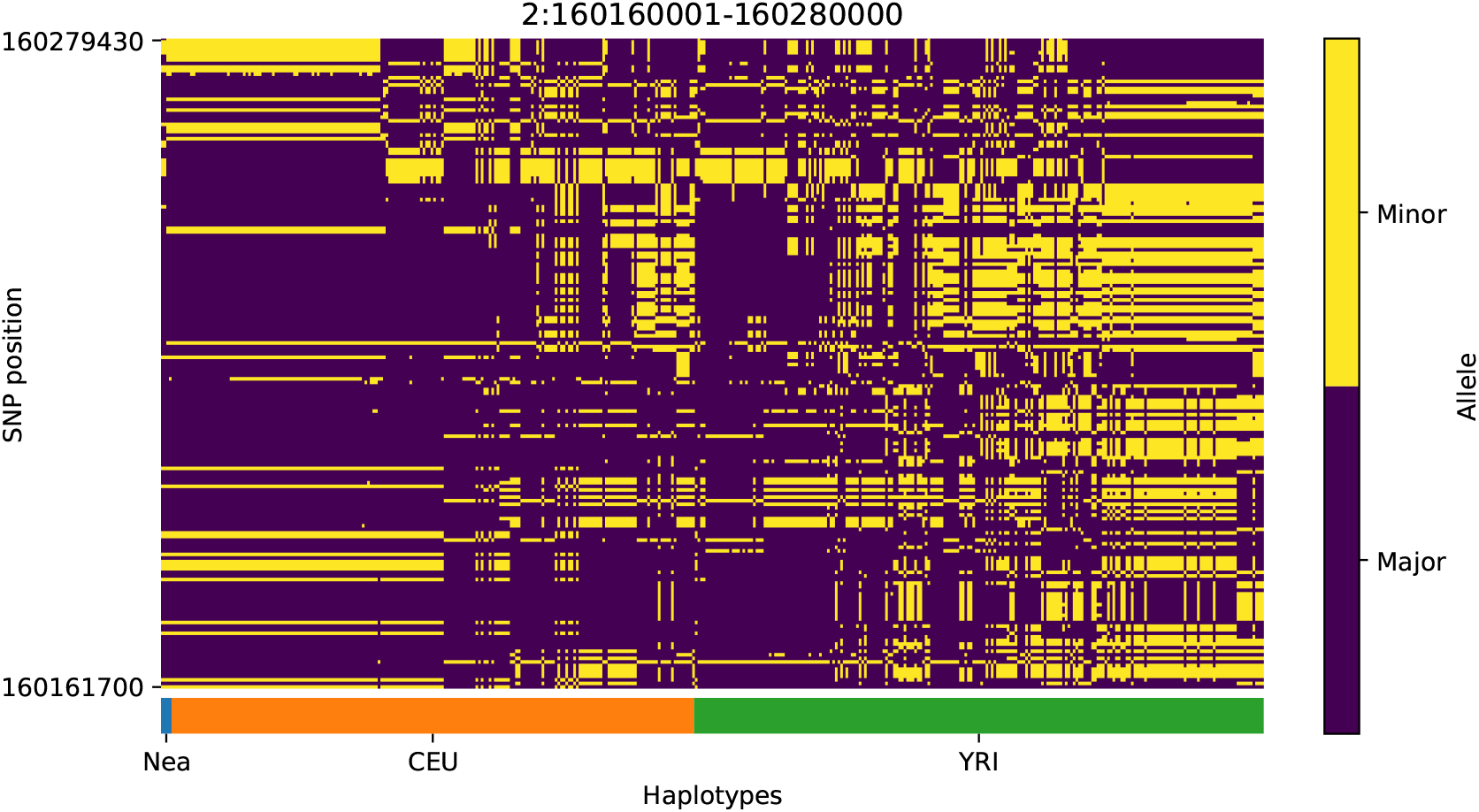
Haplotype plot for the candidate region chr2:160160001-160280000 in the Neanderthal-into-European AI scan. Bright yellow indicates minor allele, dark blue indicates major allele. Haplotypes within populations are sorted left-to-right by similarity to Neanderthals.

**Figure S14:**
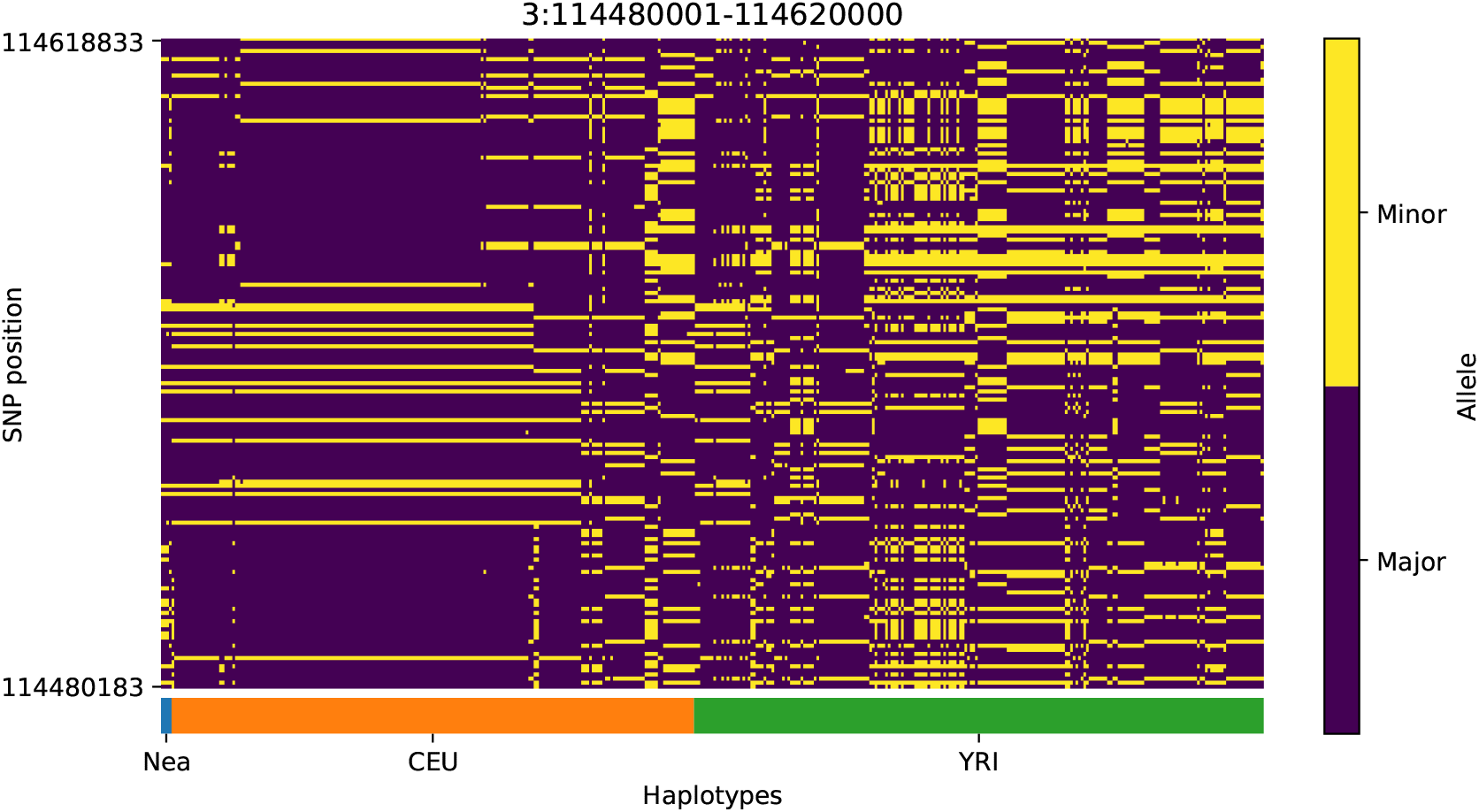
Haplotype plot for the candidate region chr3:114480001-114620000 in the Neanderthal-into-European AI scan. Bright yellow indicates minor allele, dark blue indicates major allele. Haplotypes within populations are sorted left-to-right by similarity to Neanderthals.

**Figure S15:**
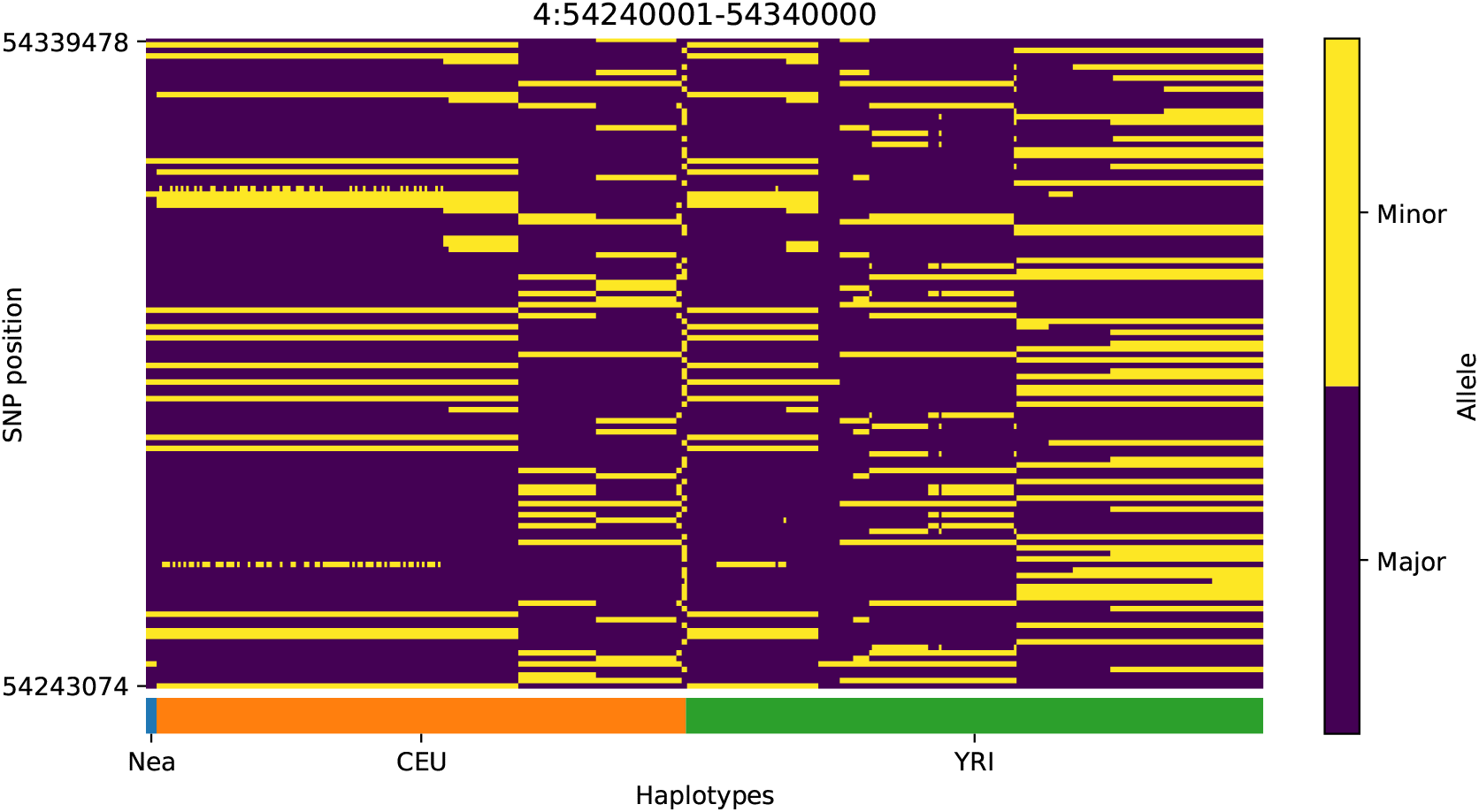
Haplotype plot for the candidate region chr4:54240001-54340000 in the Neanderthal-into-European AI scan. Bright yellow indicates minor allele, dark blue indicates major allele. Haplotypes within populations are sorted left-to-right by similarity to Neanderthals.

**Figure S16:**
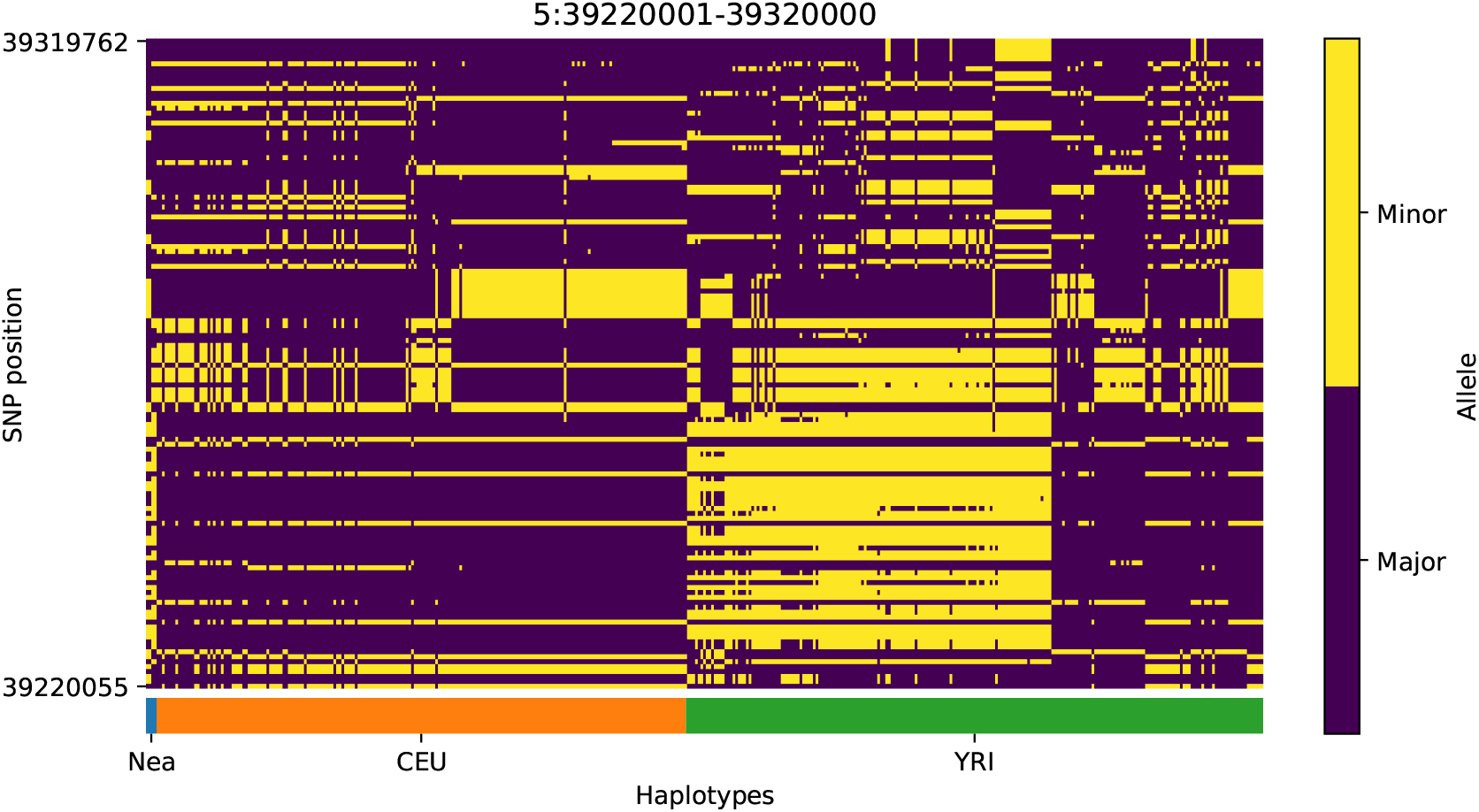
Haplotype plot for the candidate region chr5:39220001-39320000 in the Neanderthal-into-European AI scan. Bright yellow indicates minor allele, dark blue indicates major allele. Haplotypes within populations are sorted left-to-right by similarity to Neanderthals.

**Figure S17:**
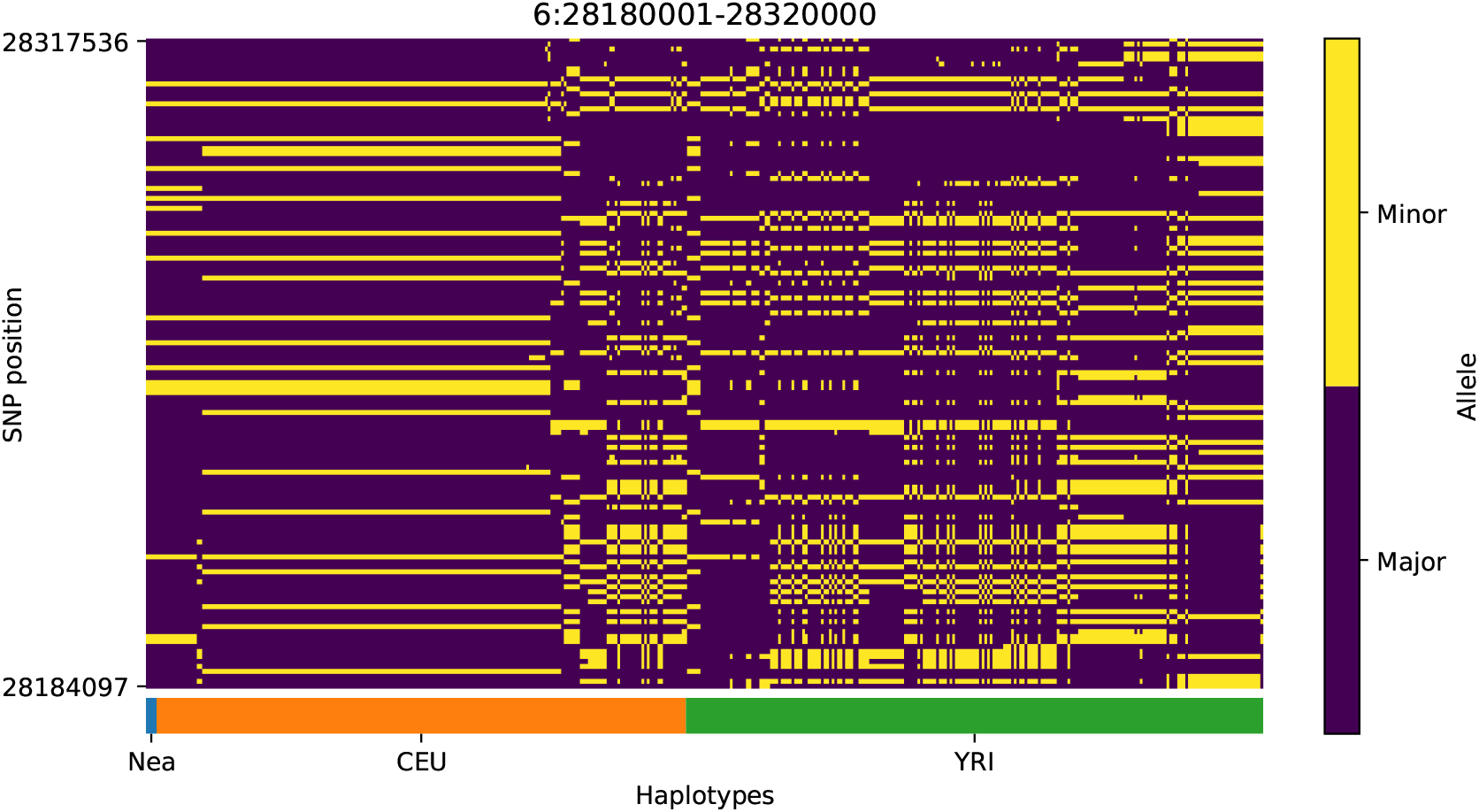
Haplotype plot for the candidate region chr6:28180001-28320000 in the Neanderthal-into-European AI scan. Bright yellow indicates minor allele, dark blue indicates major allele. Haplotypes within populations are sorted left-to-right by similarity to Neanderthals.

**Figure S18:**
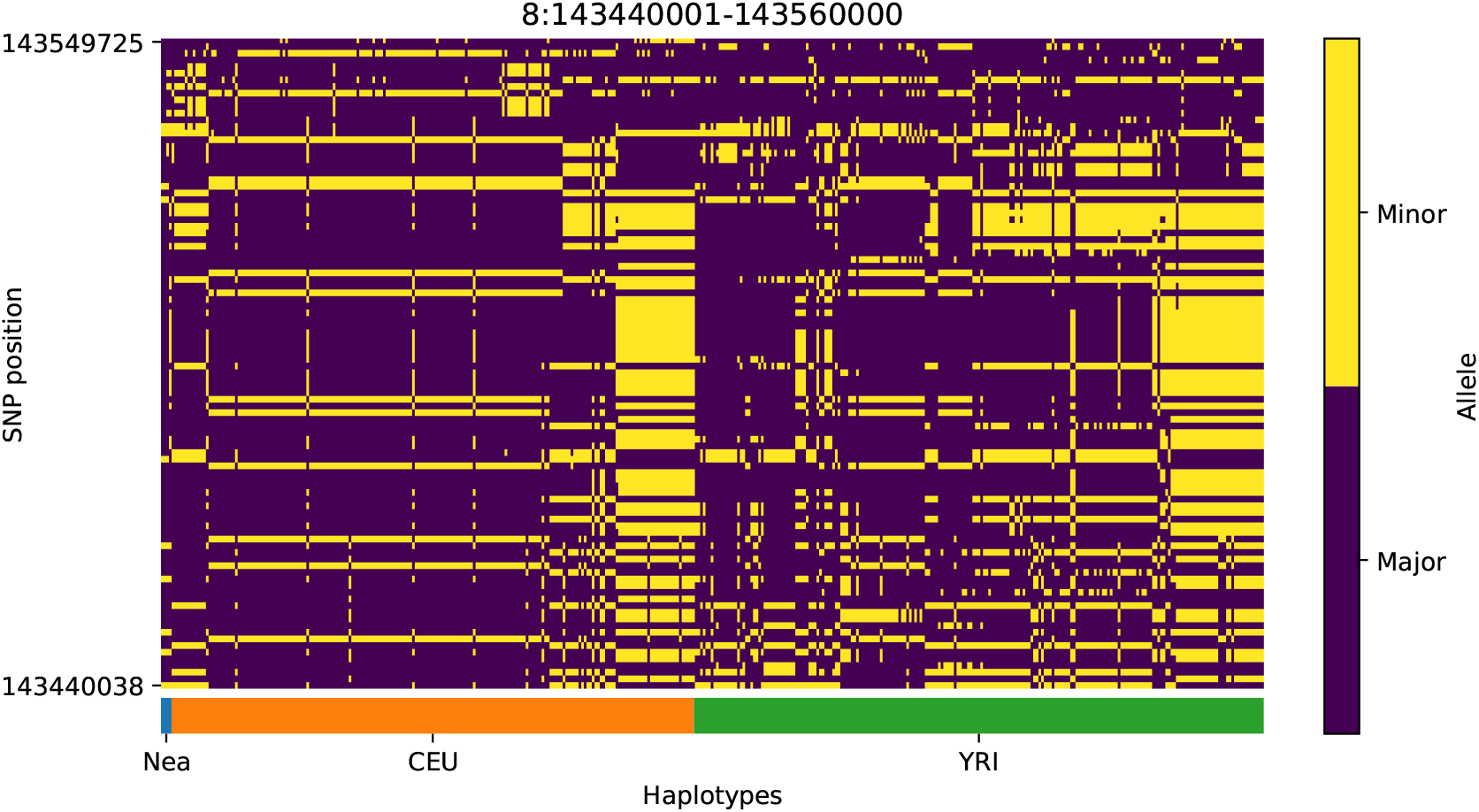
Haplotype plot for the candidate region chr8:143440001-143560000 in the Neanderthal-into-European AI scan. Bright yellow indicates minor allele, dark blue indicates major allele. Haplotypes within populations are sorted left-to-right by similarity to Neanderthals.

**Figure S19:**
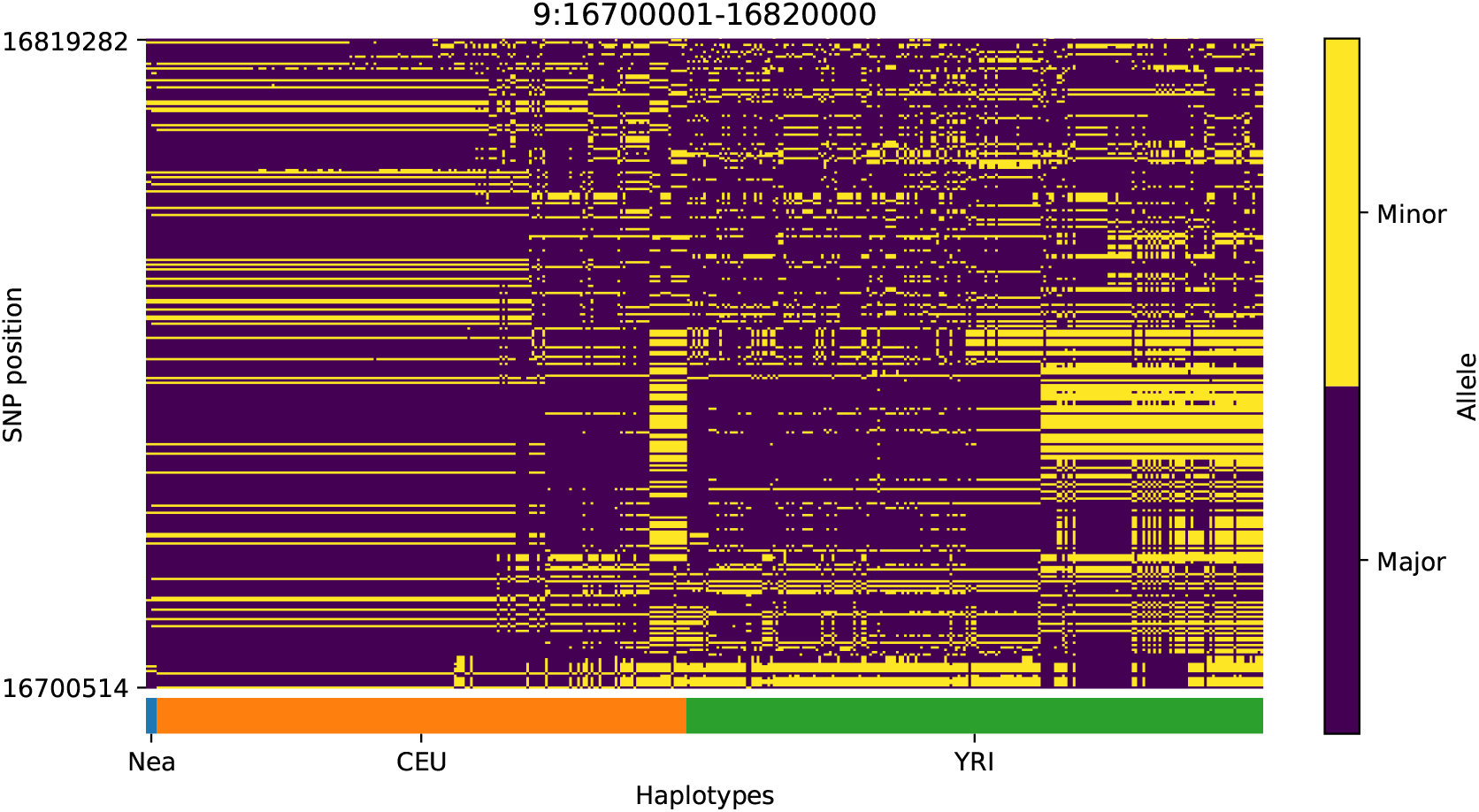
Haplotype plot for the candidate region chr9:16700001-16820000 in the Neanderthal-into-European AI scan. Bright yellow indicates minor allele, dark blue indicates major allele. Haplotypes within populations are sorted left-to-right by similarity to Neanderthals.

**Figure S20:**
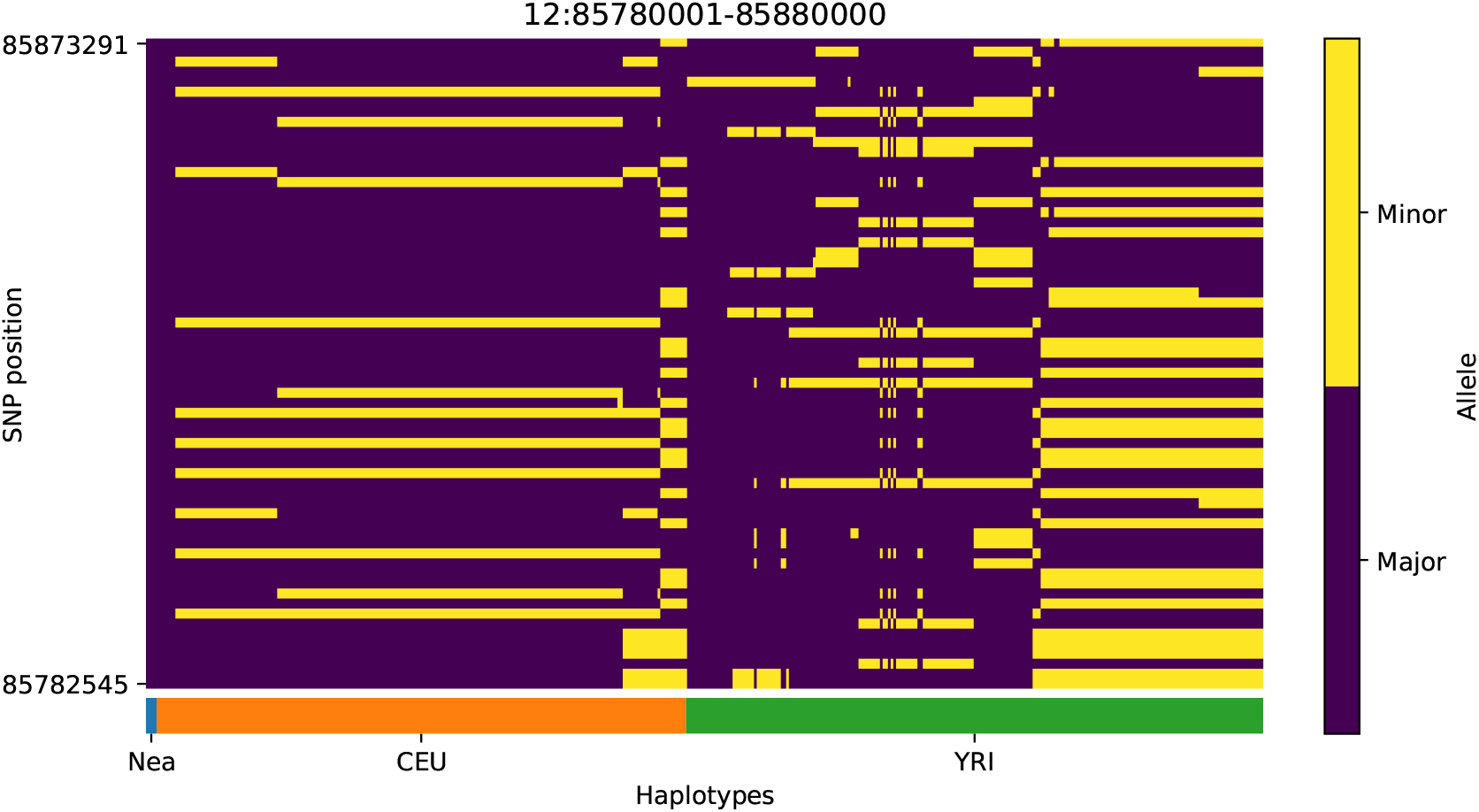
Haplotype plot for the candidate region chr12:85780001-85880000 in the Neanderthal-into-European AI scan. Bright yellow indicates minor allele, dark blue indicates major allele. Haplotypes within populations are sorted left-to-right by similarity to Neanderthals.

**Figure S21:**
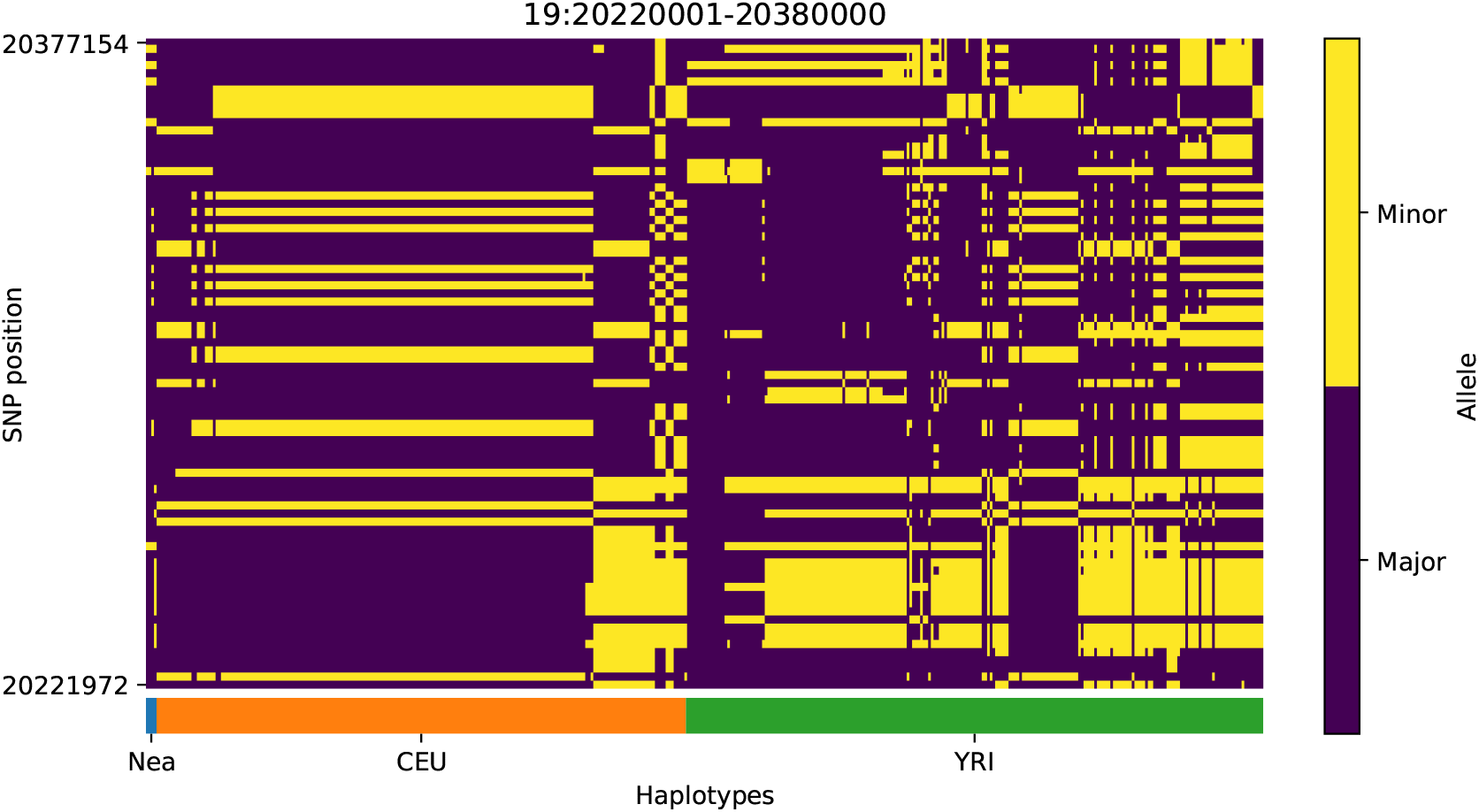
Haplotype plot for the candidate region chr19:20220001-20380000 in the Neanderthal-into-European AI scan. Bright yellow indicates minor allele, dark blue indicates major allele. Haplotypes within populations are sorted left-to-right by similarity to Neanderthals.

**Figure S22:**
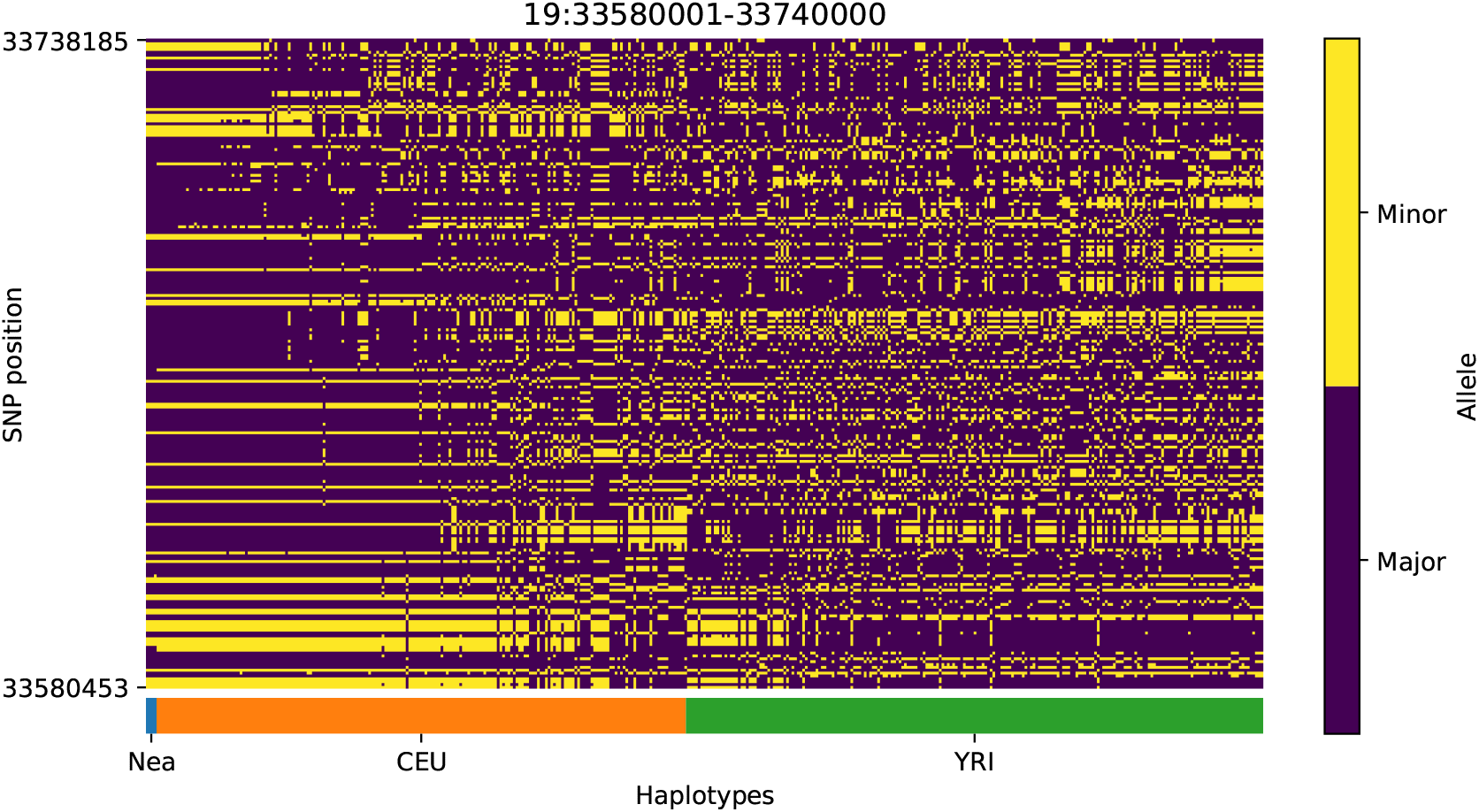
Haplotype plot for the candidate region chr19:33580001-33740000 in the Neanderthal-into-European AI scan. Bright yellow indicates minor allele, dark blue indicates major allele. Haplotypes within populations are sorted left-to-right by similarity to Neanderthals.

**Figure S23:**
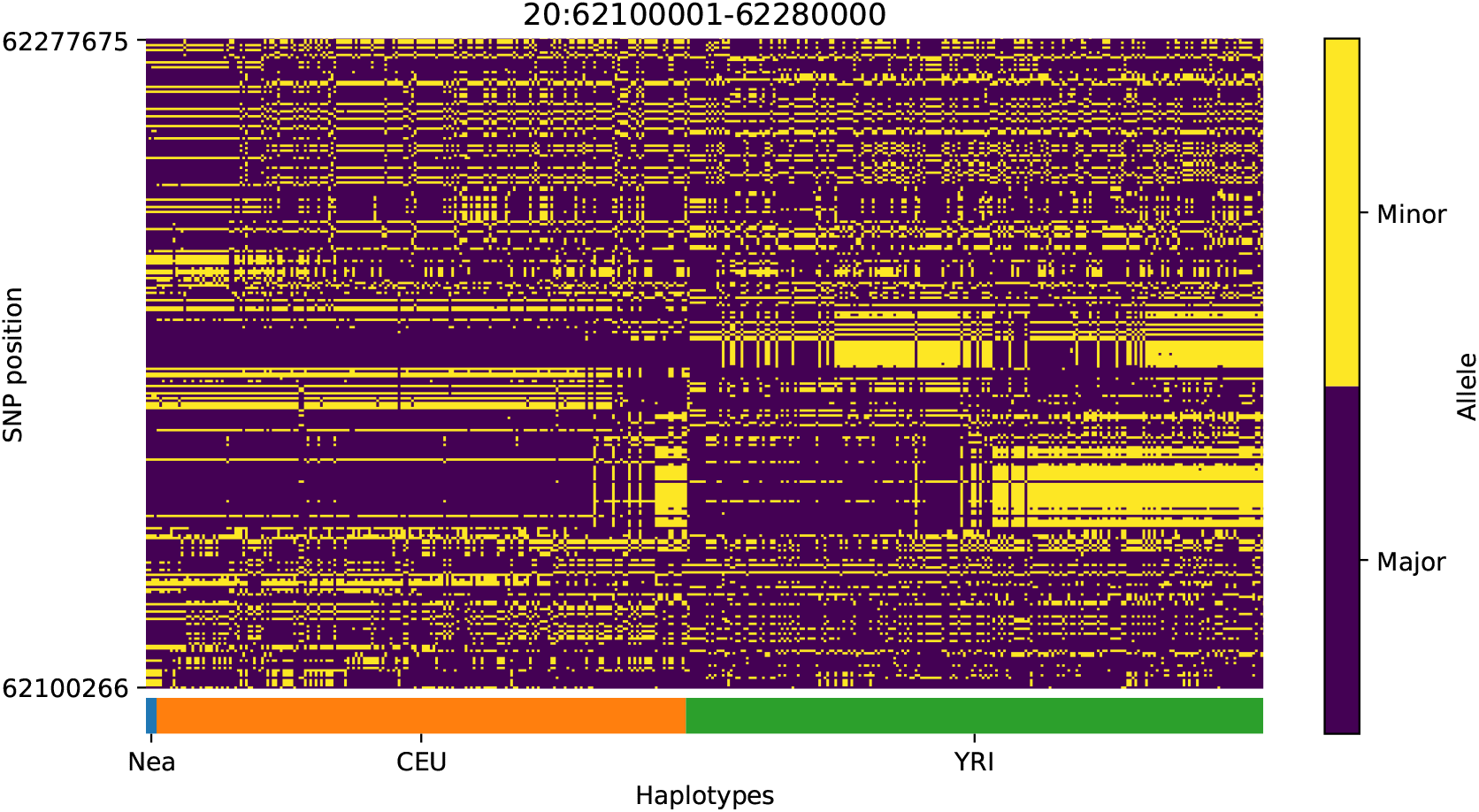
Haplotype plot for the candidate region chr20:62100001-62280000 in the Neanderthal-into-European AI scan. Bright yellow indicates minor allele, dark blue indicates major allele. Haplotypes within populations are sorted left-to-right by similarity to Neanderthals.

**Figure S24:**
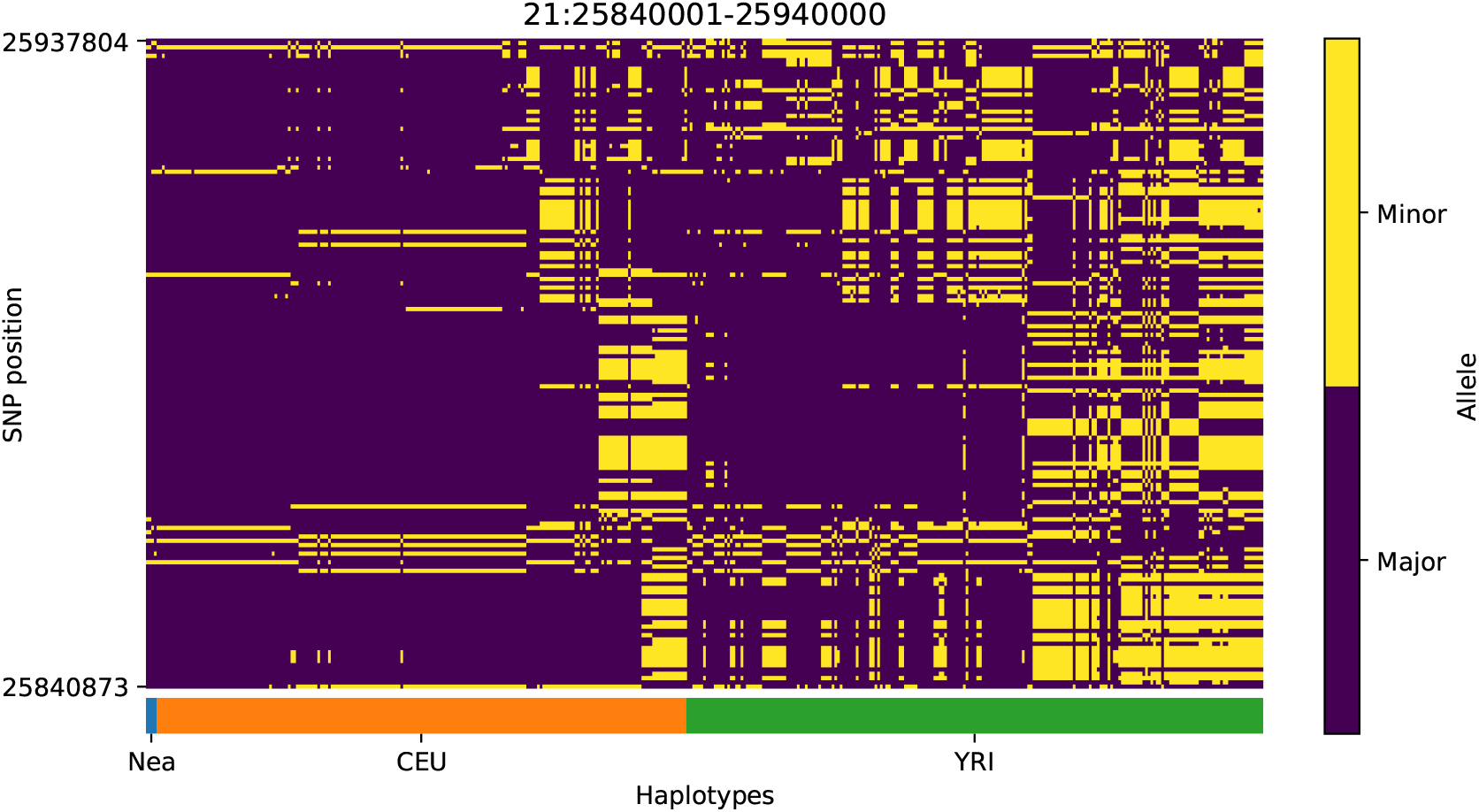
Haplotype plot for the candidate region chr21:25840001-25940000 in the Neanderthal-into-European AI scan. Bright yellow indicates minor allele, dark blue indicates major allele. Haplotypes within populations are sorted left-to-right by similarity to Neanderthals.

**Figure S25:**
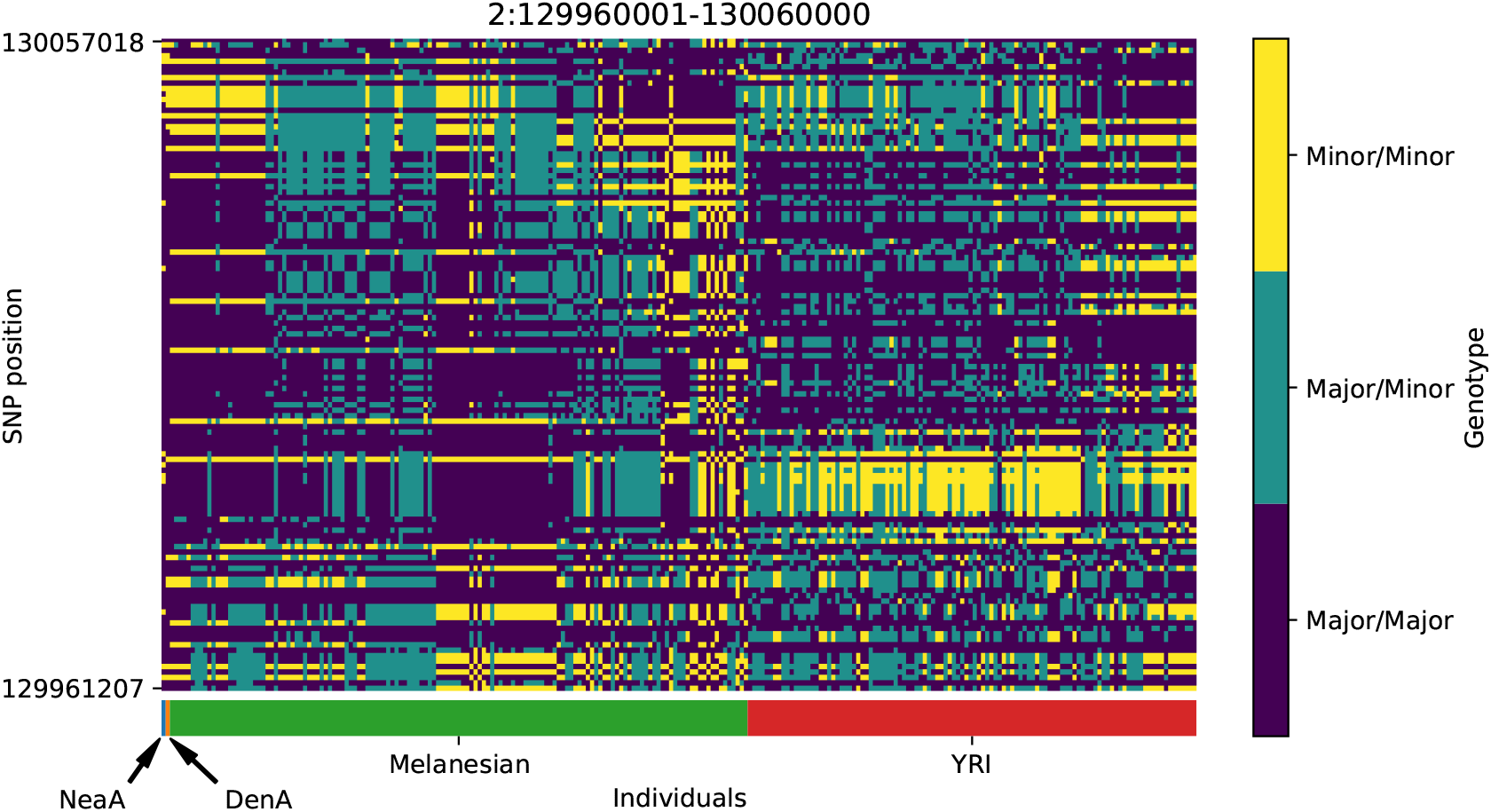
Genotype plot for the candidate region chr2:129960001-130060000 in the Denisovan-into-Melanesian AI scan. Dark blue = homozygote major allele, light blue = heterozygote, yellow = homozygote minor allele. Genotypes within populations are sorted left-to-right by similarity to the Denisovan.

**Figure S26:**
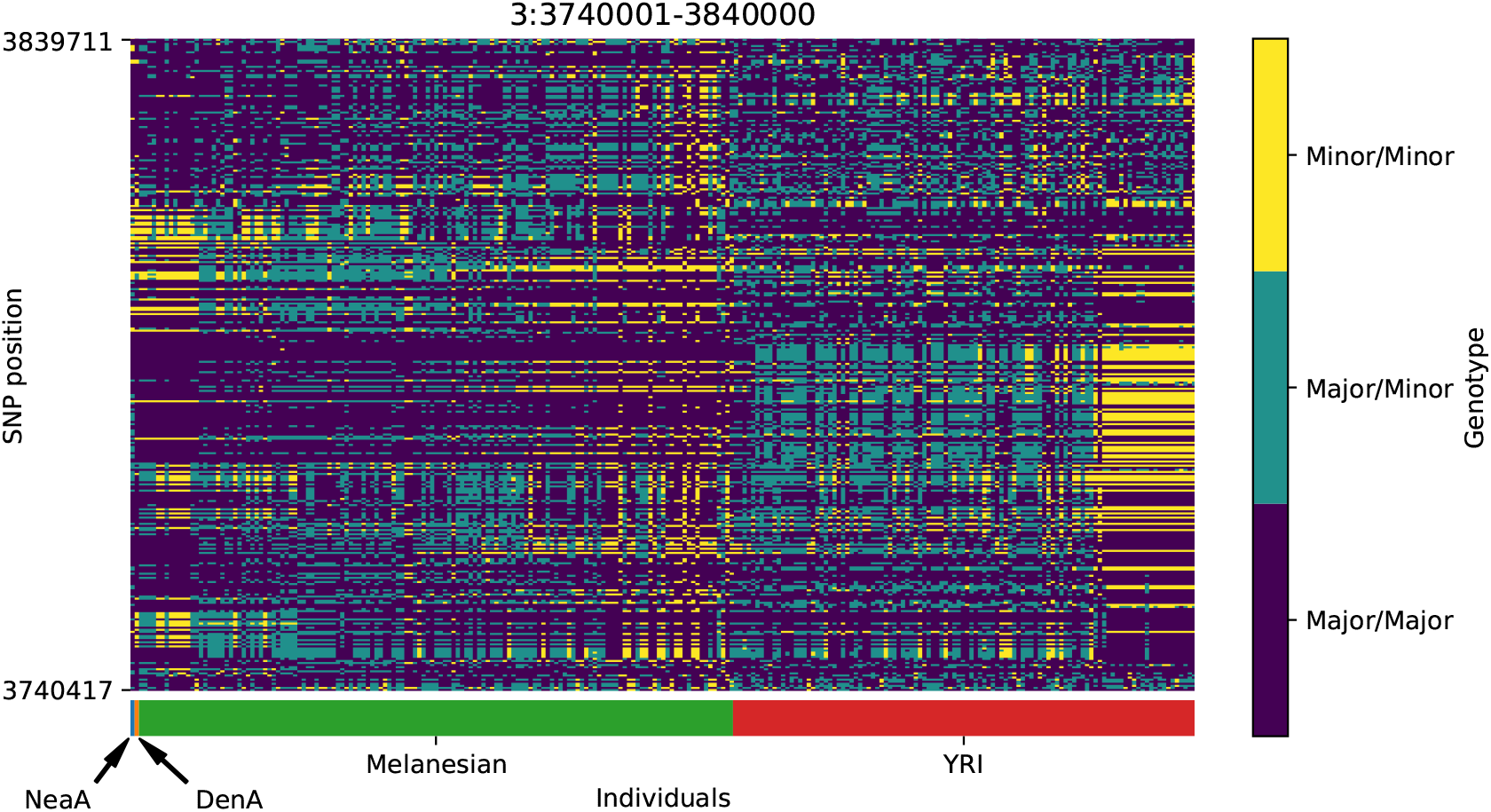
Genotype plot for the candidate region chr3:3740001-3840000 in the Denisovan-into-Melanesian AI scan. Dark blue = homozygote major allele, light blue = heterozygote, yellow = homozygote minor allele. Genotypes within populations are sorted left-to-right by similarity to the Denisovan.

**Figure S27:**
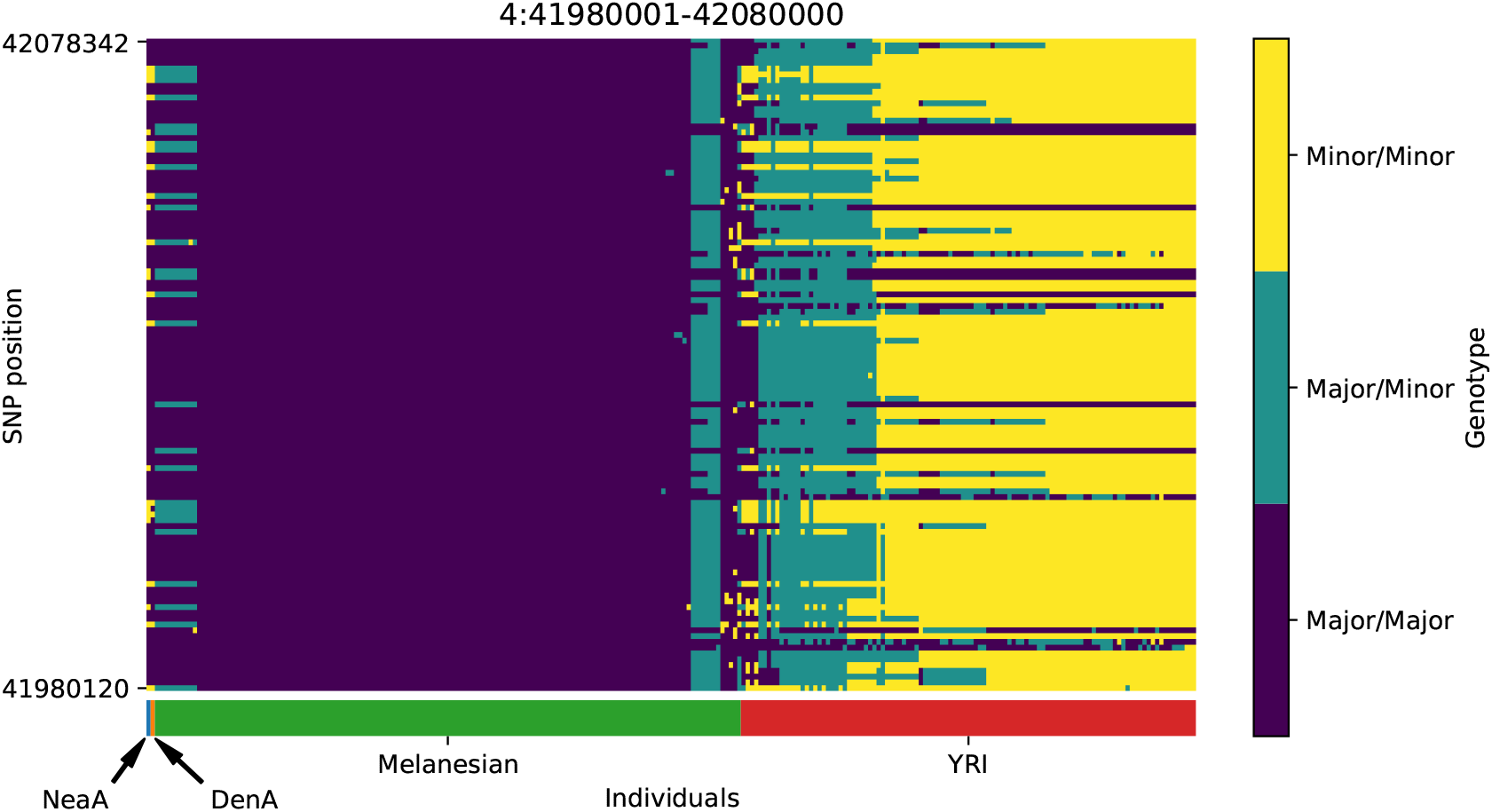
Genotype plot for the candidate region chr4:41980001-42080000 in the Denisovan-into-Melanesian AI scan. Dark blue = homozygote major allele, light blue = heterozygote, yellow = homozygote minor allele. Genotypes within populations are sorted left-to-right by similarity to the Denisovan.

**Figure S28:**
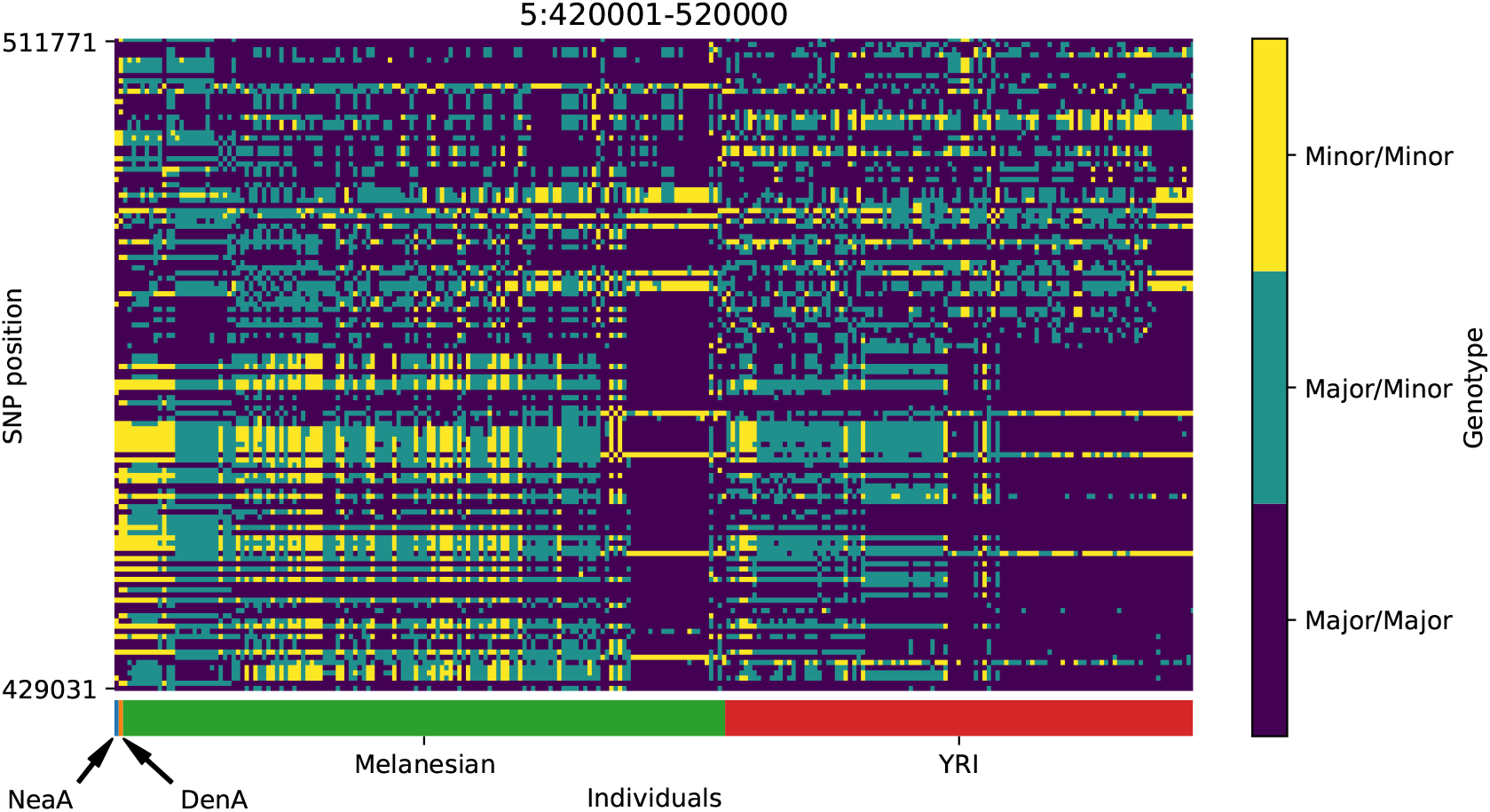
Genotype plot for the candidate region chr5:420001-520000 in the Denisovan-into-Melanesian AI scan. Dark blue = homozygote major allele, light blue = heterozygote, yellow = homozygote minor allele. Genotypes within populations are sorted left-to-right by similarity to the Denisovan.

**Figure S29:**
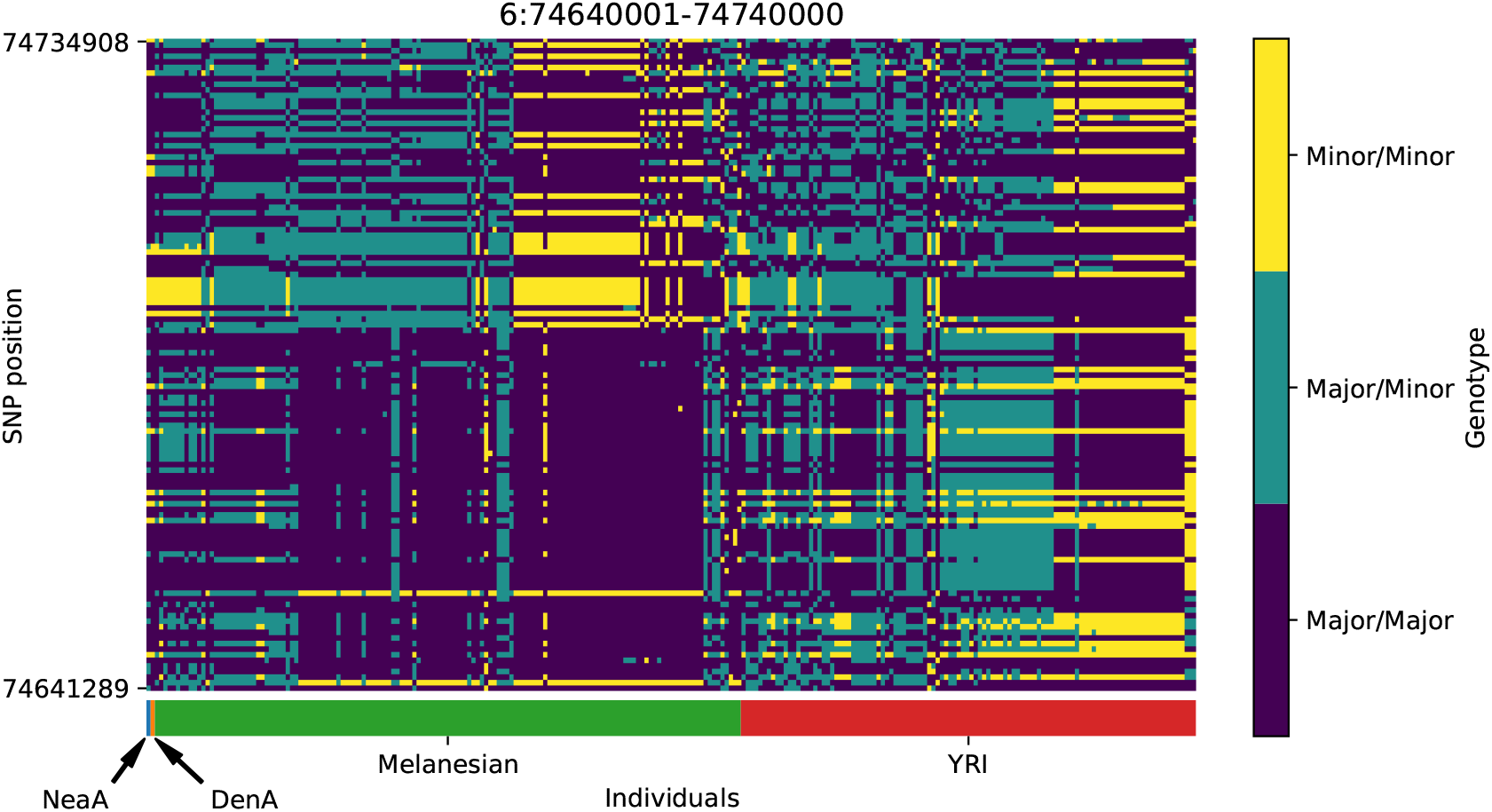
Genotype plot for the candidate region chr6:74640001-74740000 in the Denisovan-into-Melanesian AI scan. Dark blue = homozygote major allele, light blue = heterozygote, yellow = homozygote minor allele. Genotypes within populations are sorted left-to-right by similarity to the Denisovan.

**Figure S30:**
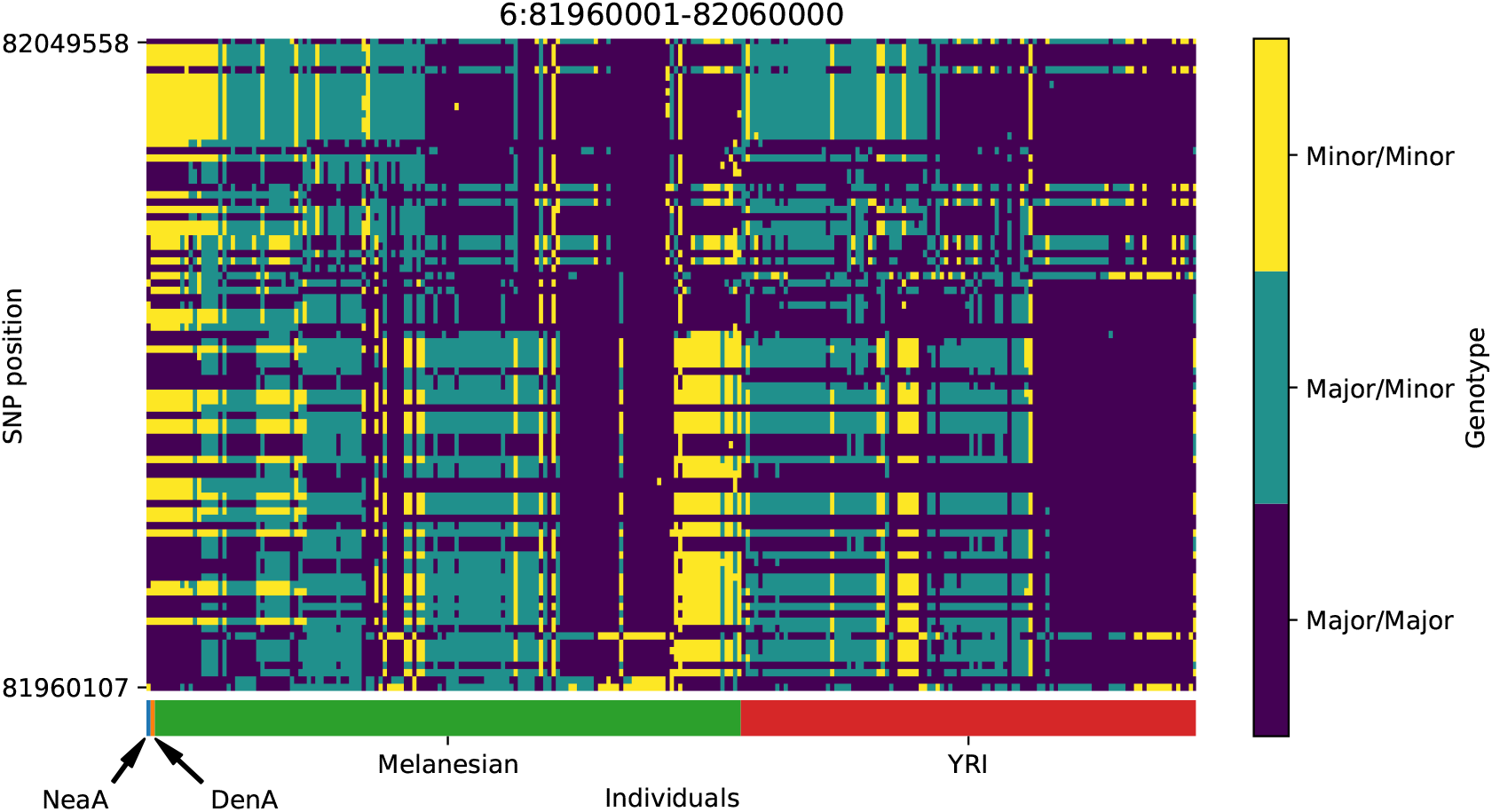
Genotype plot for the candidate region chr6:81960001-82060000 in the Denisovan-into-Melanesian AI scan. Dark blue = homozygote major allele, light blue = heterozygote, yellow = homozygote minor allele. Genotypes within populations are sorted left-to-right by similarity to the Denisovan.

**Figure S31:**
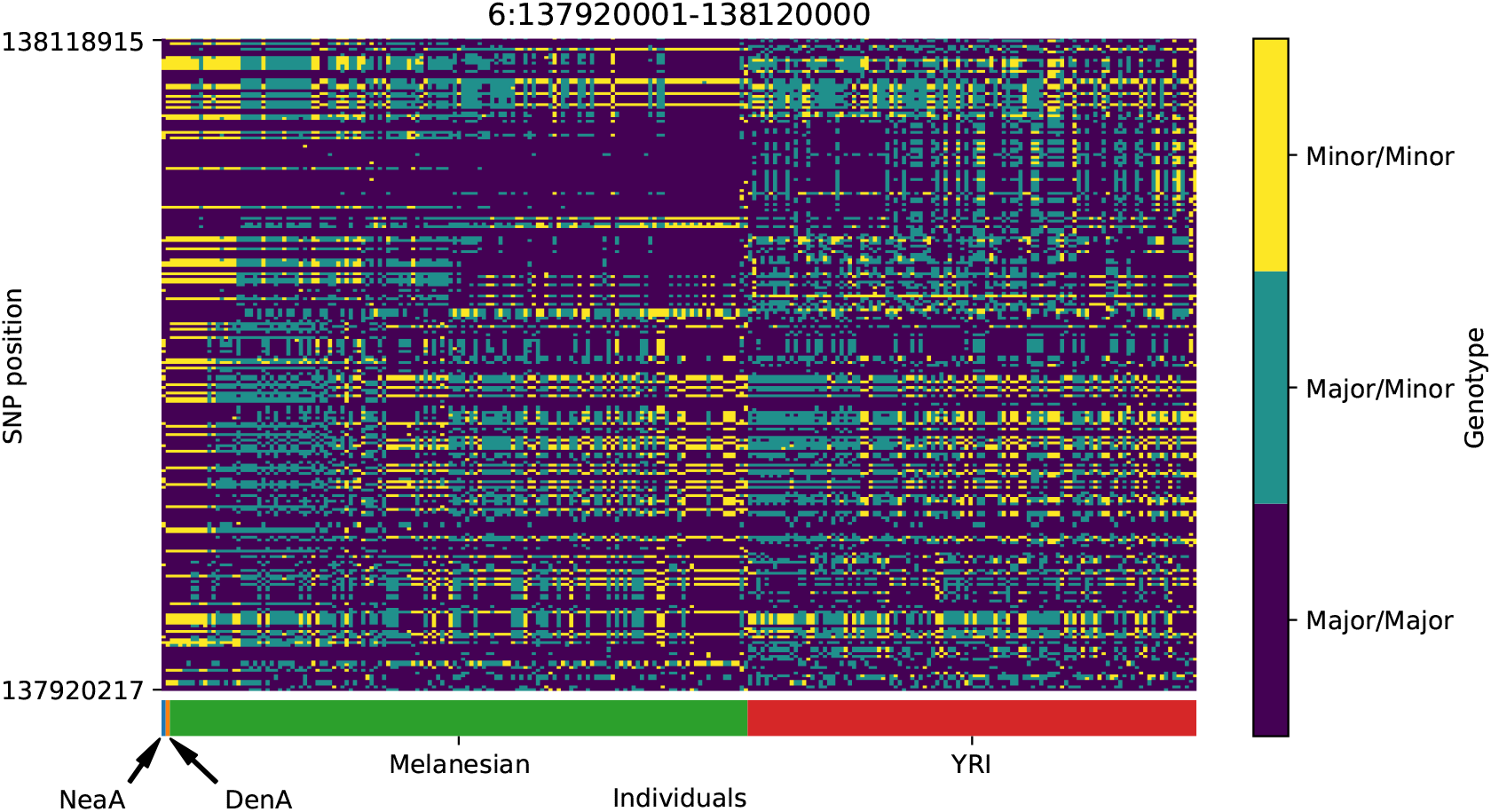
Genotype plot for the candidate region chr6:137920001-138120000 in the Denisovan-into-Melanesian AI scan. Dark blue = homozygote major allele, light blue = heterozygote, yellow = homozygote minor allele. Genotypes within populations are sorted left-to-right by similarity to the Denisovan.

**Figure S32:**
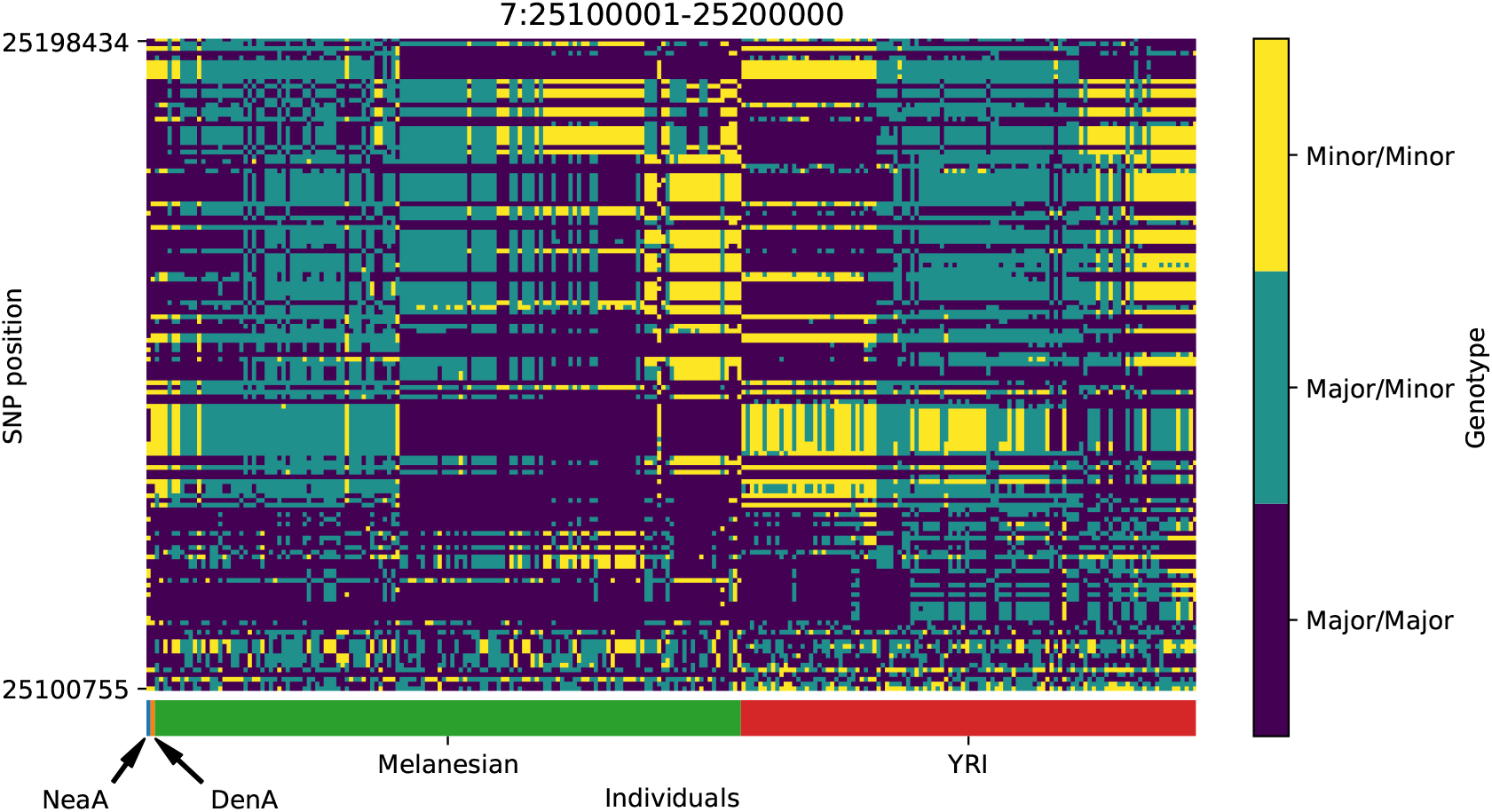
Genotype plot for the candidate region chr7:25100001-25200000 in the Denisovan-into-Melanesian AI scan. Dark blue = homozygote major allele, light blue = heterozygote, yellow = homozygote minor allele. Genotypes within populations are sorted left-to-right by similarity to the Denisovan.

**Figure S33:**
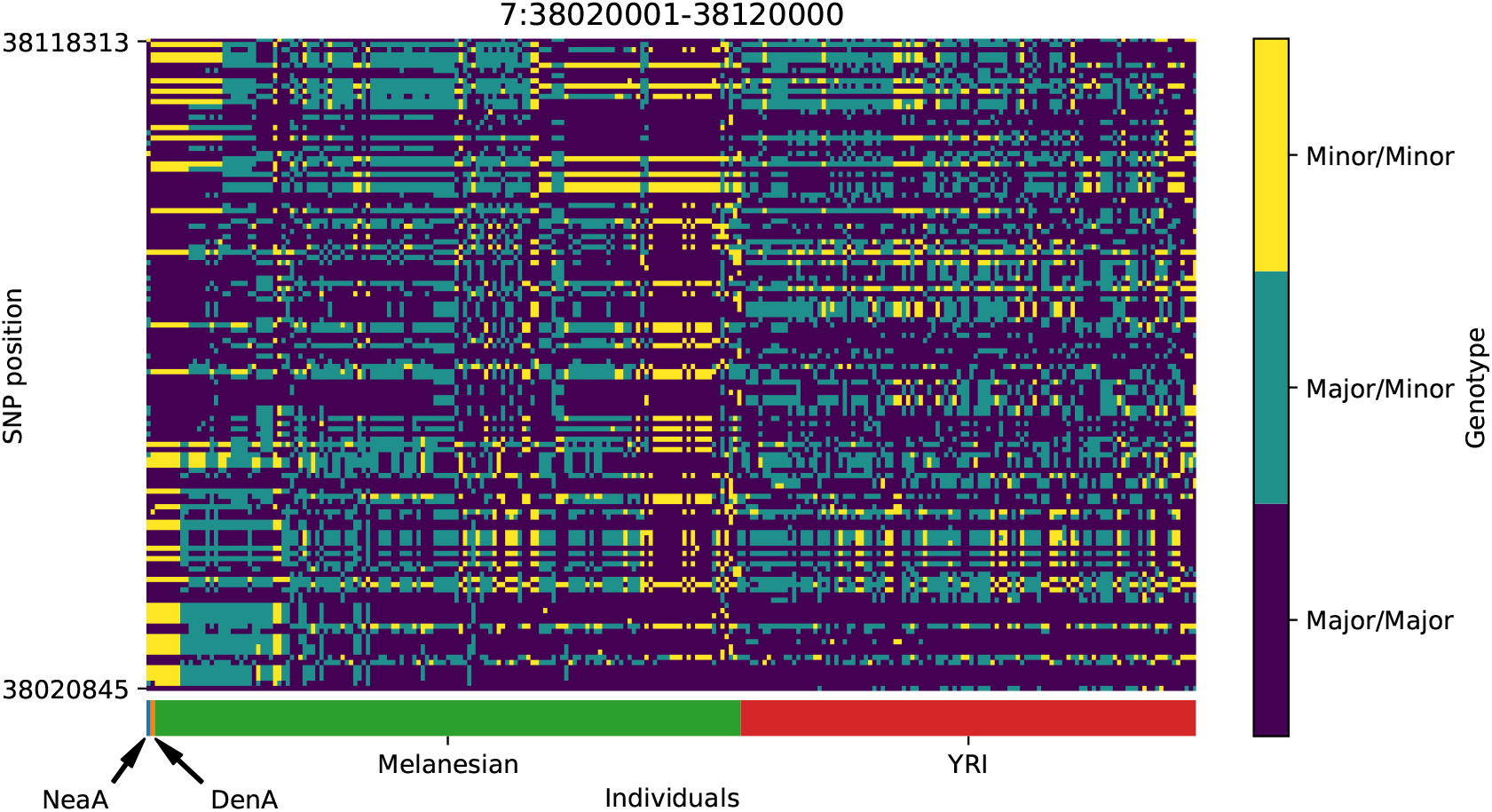
Genotype plot for the candidate region chr7:38020001-38120000 in the Denisovan-into-Melanesian AI scan. Dark blue = homozygote major allele, light blue = heterozygote, yellow = homozygote minor allele. Genotypes within populations are sorted left-to-right by similarity to the Denisovan.

**Figure S34:**
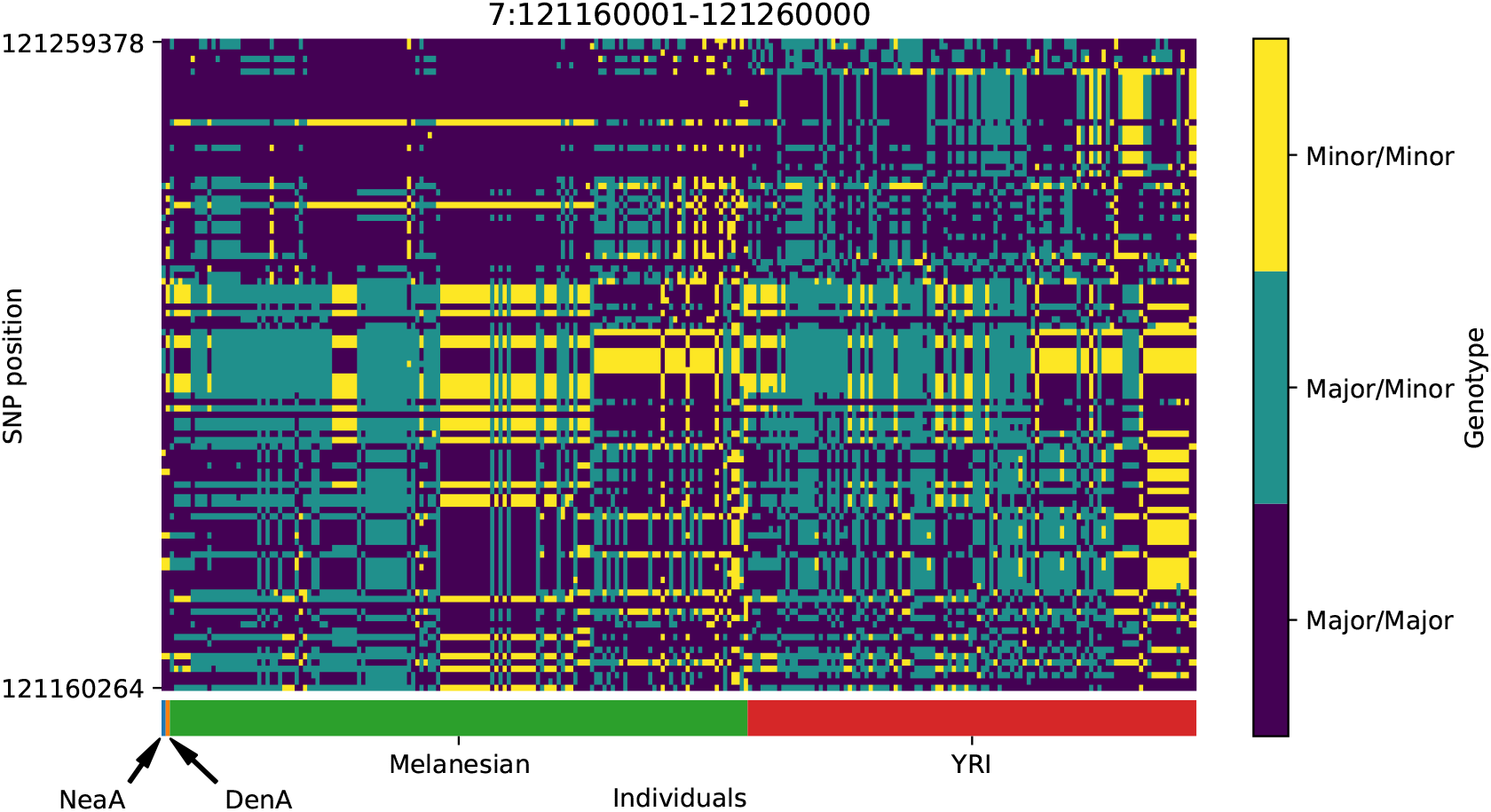
Genotype plot for the candidate region chr7:121160001-121260000 in the Denisovan-into-Melanesian AI scan. Dark blue = homozygote major allele, light blue = heterozygote, yellow = homozygote minor allele. Genotypes within populations are sorted left-to-right by similarity to the Denisovan.

**Figure S35:**
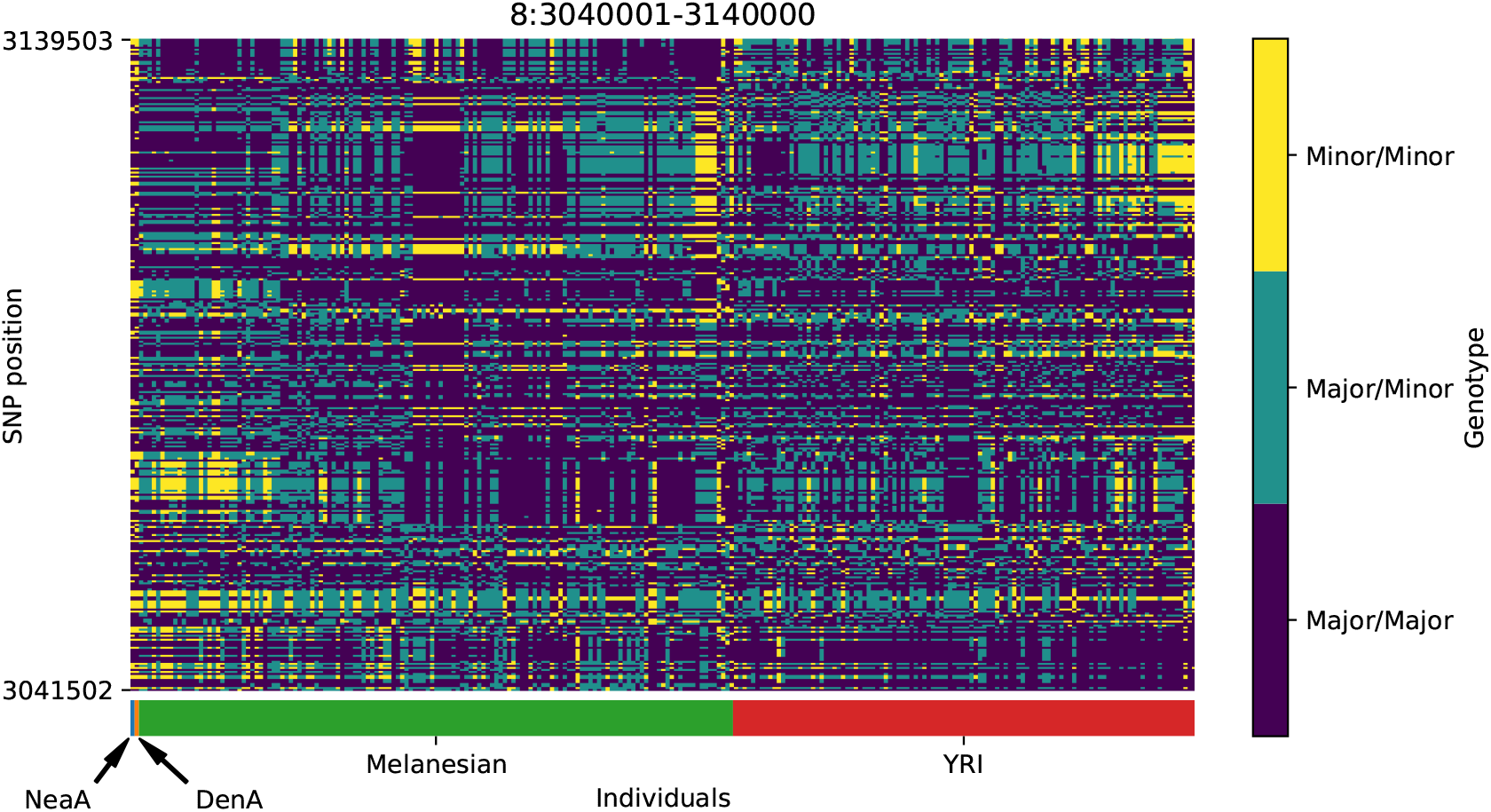
Genotype plot for the candidate region chr8:3040001-3140000 in the Denisovan-into-Melanesian AI scan. Dark blue = homozygote major allele, light blue = heterozygote, yellow = homozygote minor allele. Genotypes within populations are sorted left-to-right by similarity to the Denisovan.

**Figure S36:**
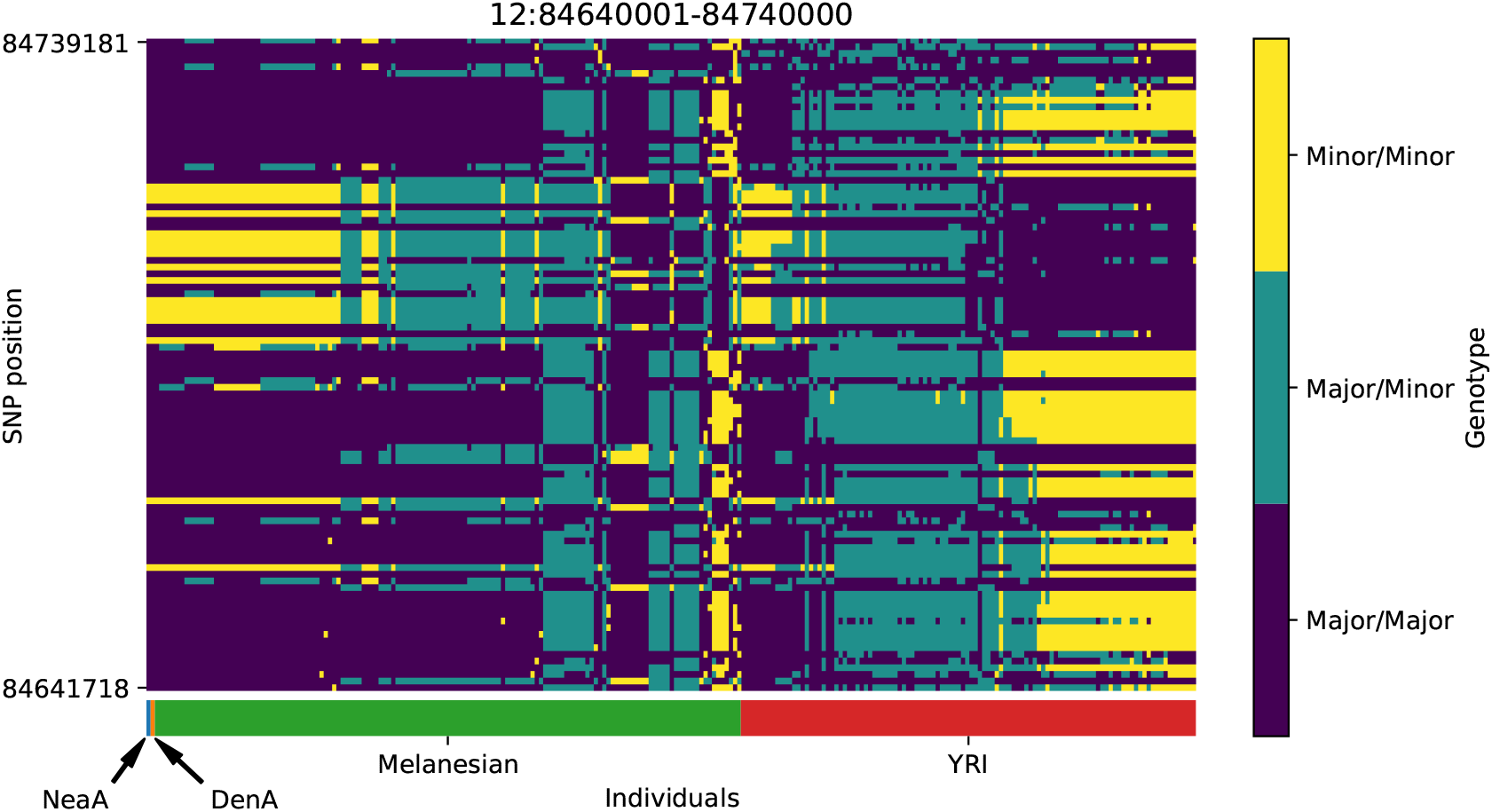
Genotype plot for the candidate region chr12:84640001-84740000 in the Denisovan-into-Melanesian AI scan. Dark blue = homozygote major allele, light blue = heterozygote, yellow = homozygote minor allele. Genotypes within populations are sorted left-to-right by similarity to the Denisovan.

**Figure S37:**
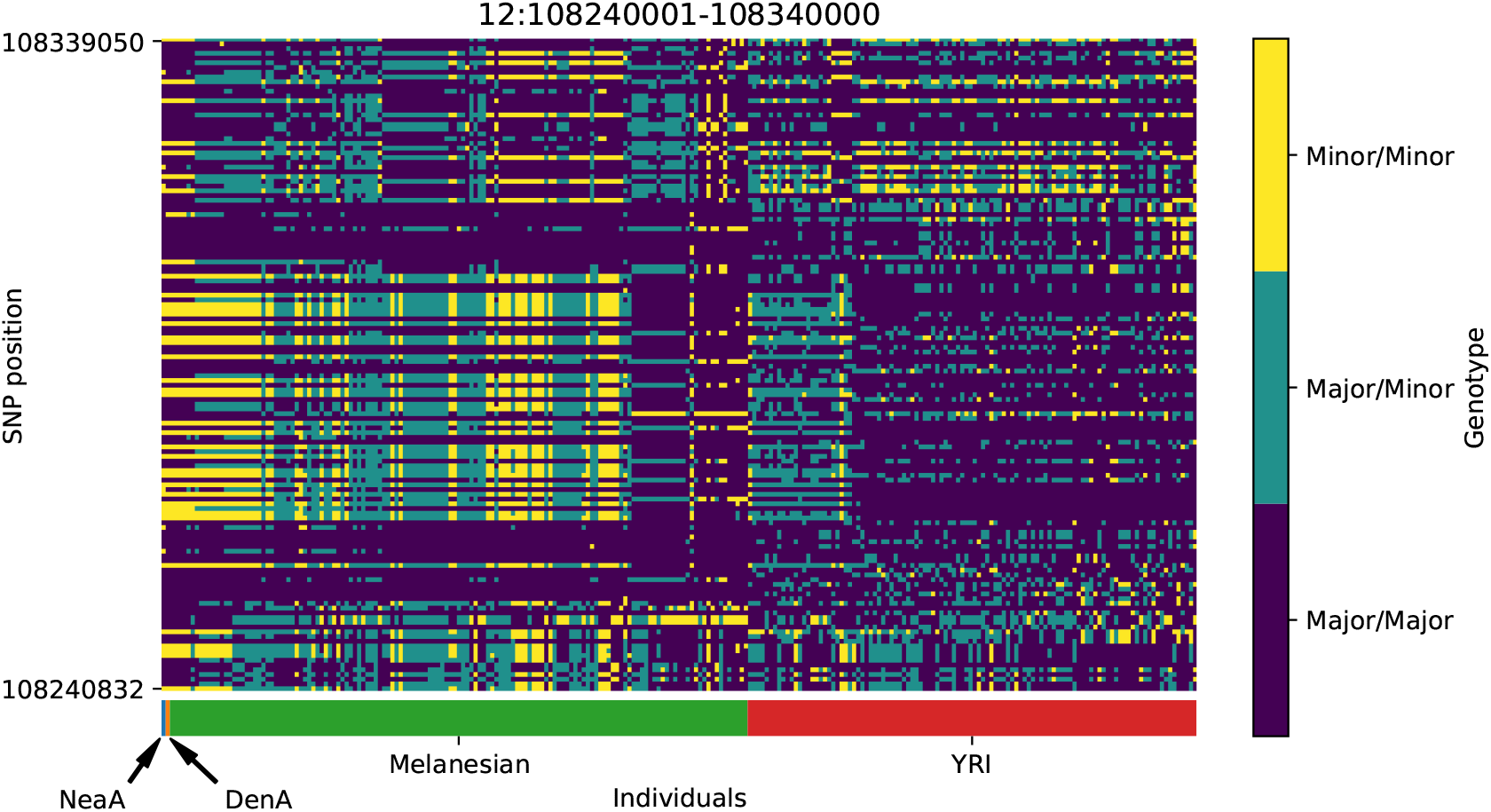
Genotype plot for the candidate region chr12:108240001-108340000 in the Denisovan-into-Melanesian AI scan. Dark blue = homozygote major allele, light blue = heterozygote, yellow = homozygote minor allele. Genotypes within populations are sorted left-to-right by similarity to the Denisovan.

**Figure S38:**
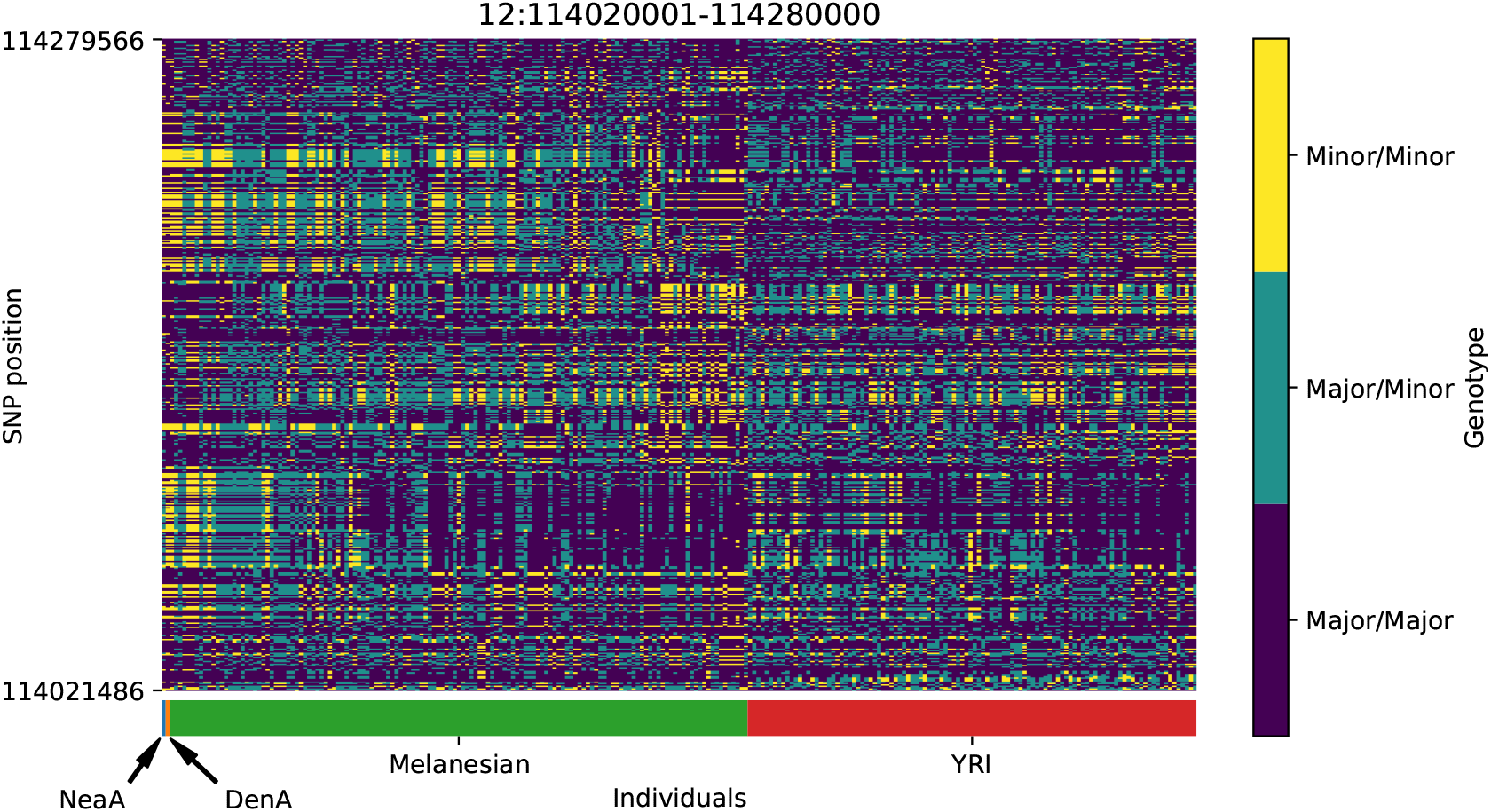
Genotype plot for the candidate region chr12:114020001-114280000 in the Denisovan-into-Melanesian AI scan. Dark blue = homozygote major allele, light blue = heterozygote, yellow = homozygote minor allele. Genotypes within populations are sorted left-to-right by similarity to the Denisovan.

**Figure S39:**
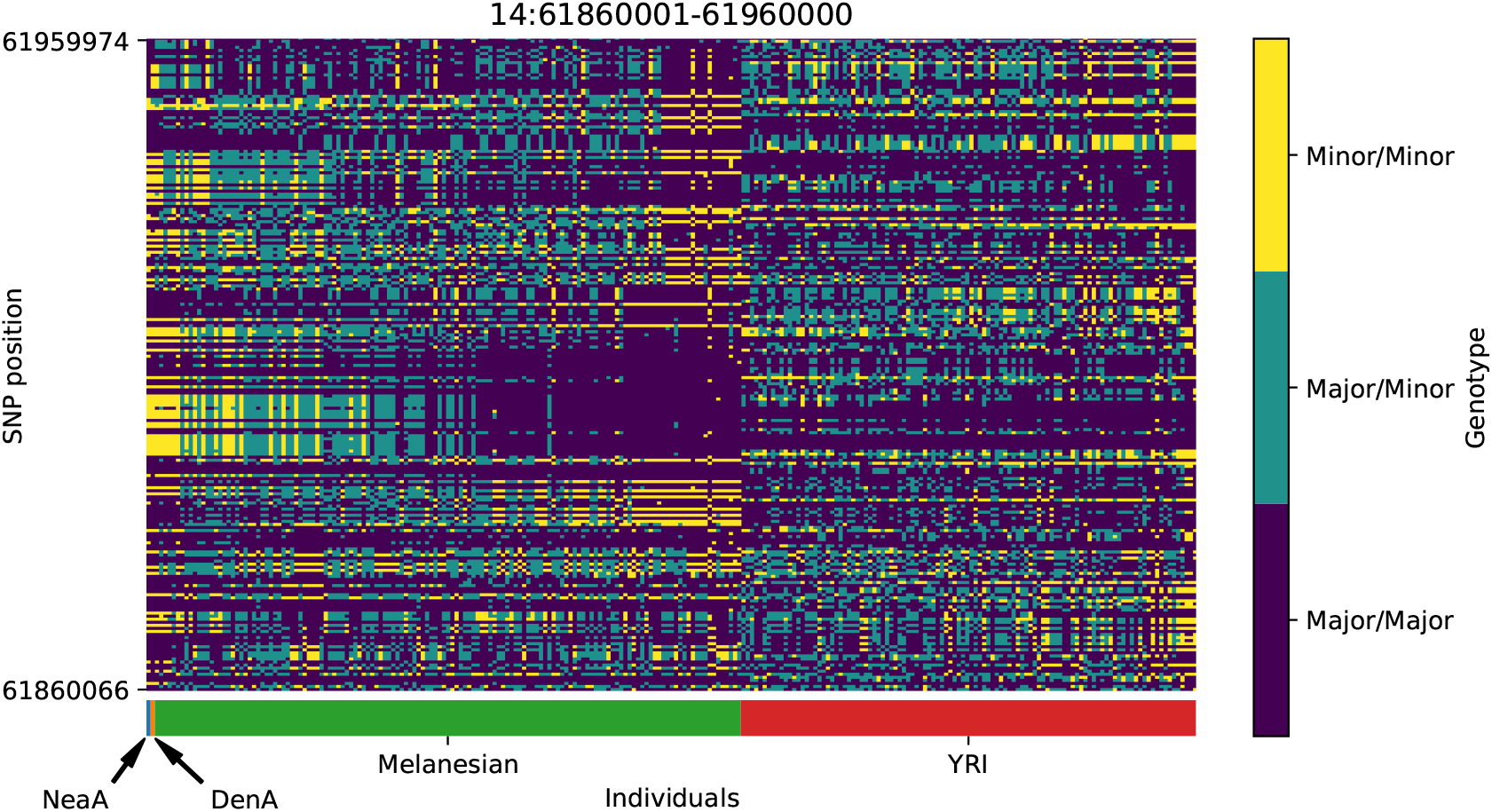
Genotype plot for the candidate region chr14:61860001-61960000 in the Denisovan-into-Melanesian AI scan. Dark blue = homozygote major allele, light blue = heterozygote, yellow = homozygote minor allele. Genotypes within populations are sorted left-to-right by similarity to the Denisovan.

**Figure S40:**
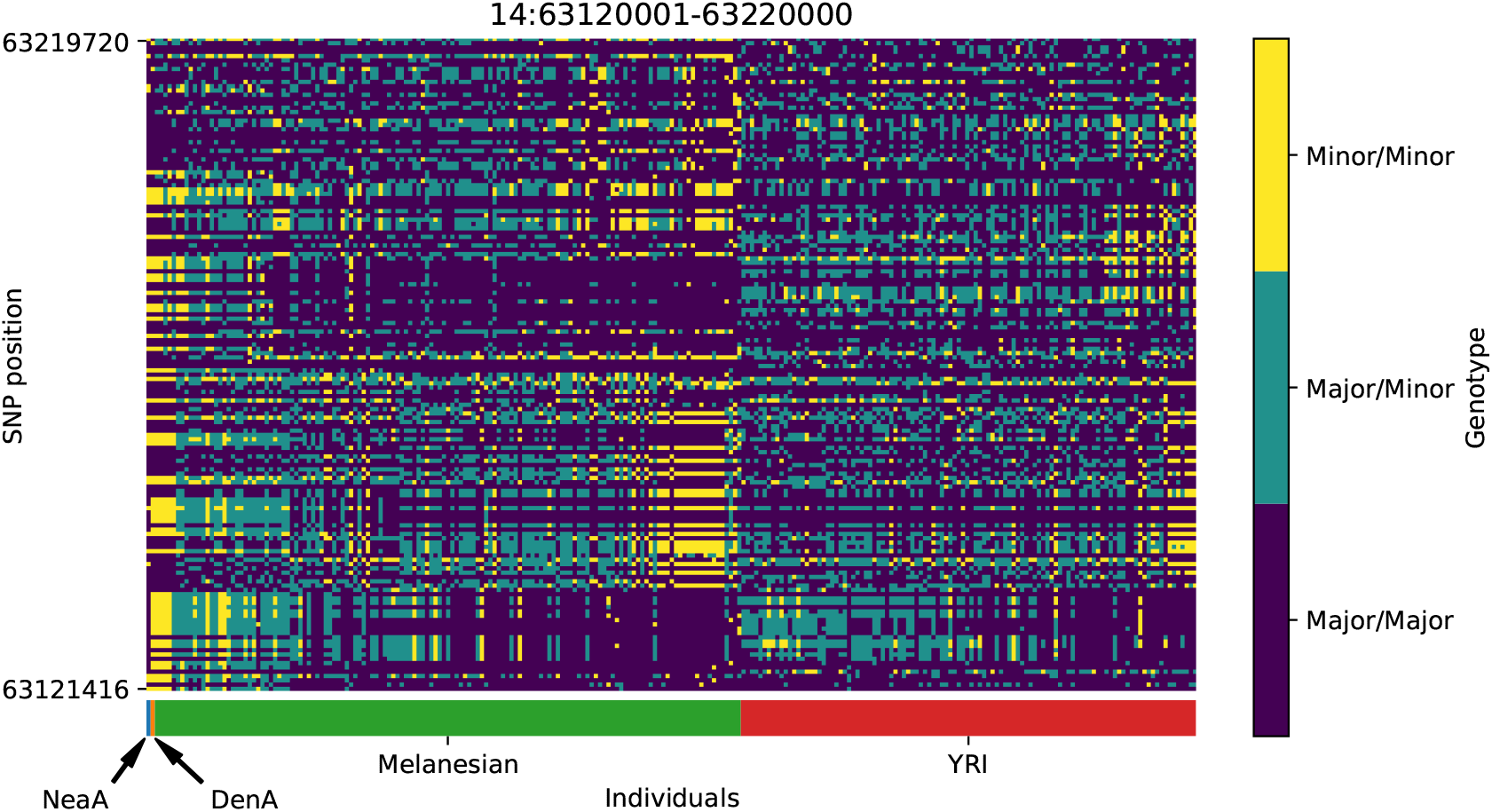
Genotype plot for the candidate region chr14:63120001-63220000 in the Denisovan-into-Melanesian AI scan. Dark blue = homozygote major allele, light blue = heterozygote, yellow = homozygote minor allele. Genotypes within populations are sorted left-to-right by similarity to the Denisovan.

**Figure S41:**
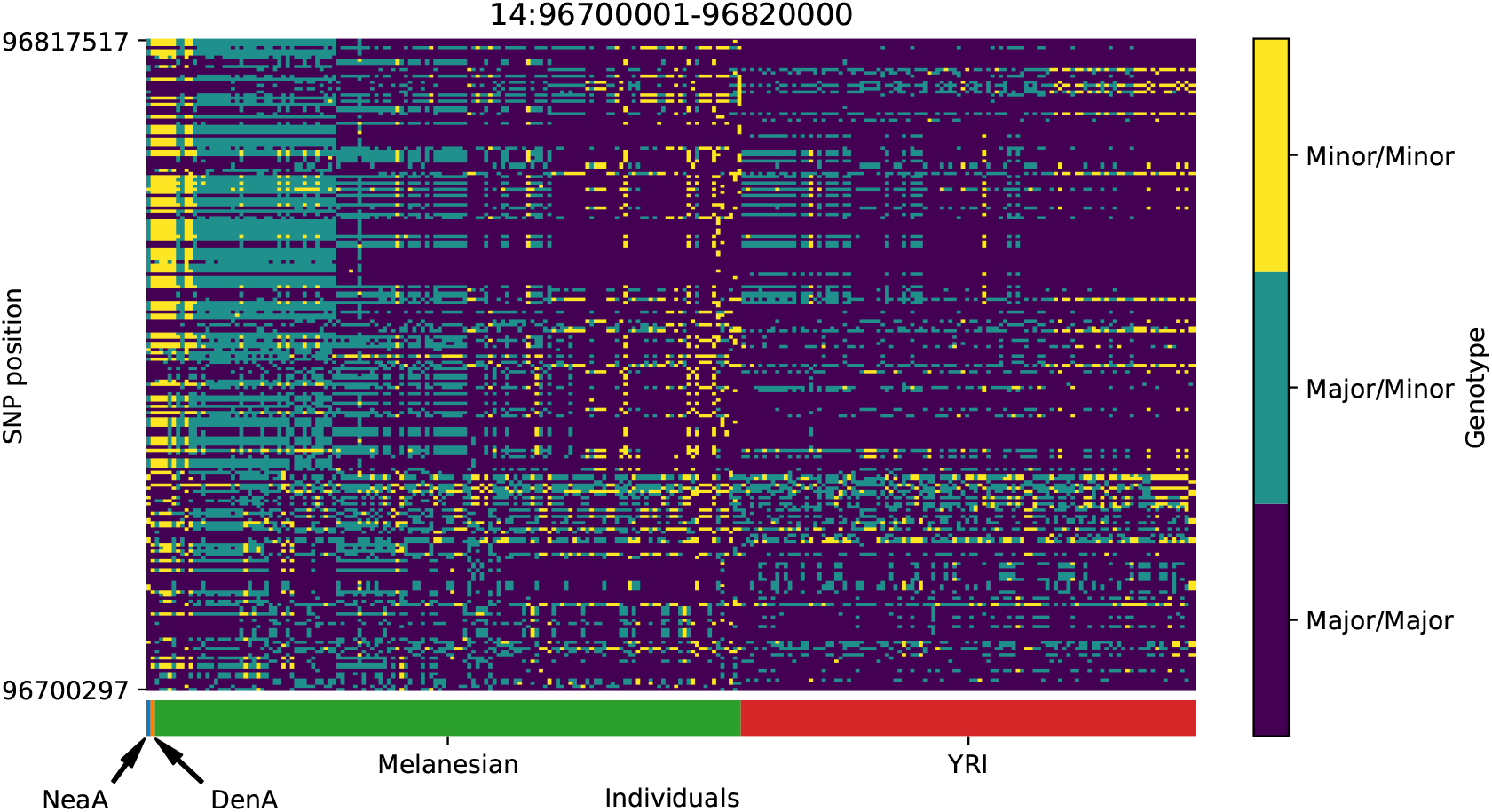
Genotype plot for the candidate region chr14:96700001-96820000 in the Denisovan-into-Melanesian AI scan. Dark blue = homozygote major allele, light blue = heterozygote, yellow = homozygote minor allele. Genotypes within populations are sorted left-to-right by similarity to the Denisovan.

**Figure S42:**
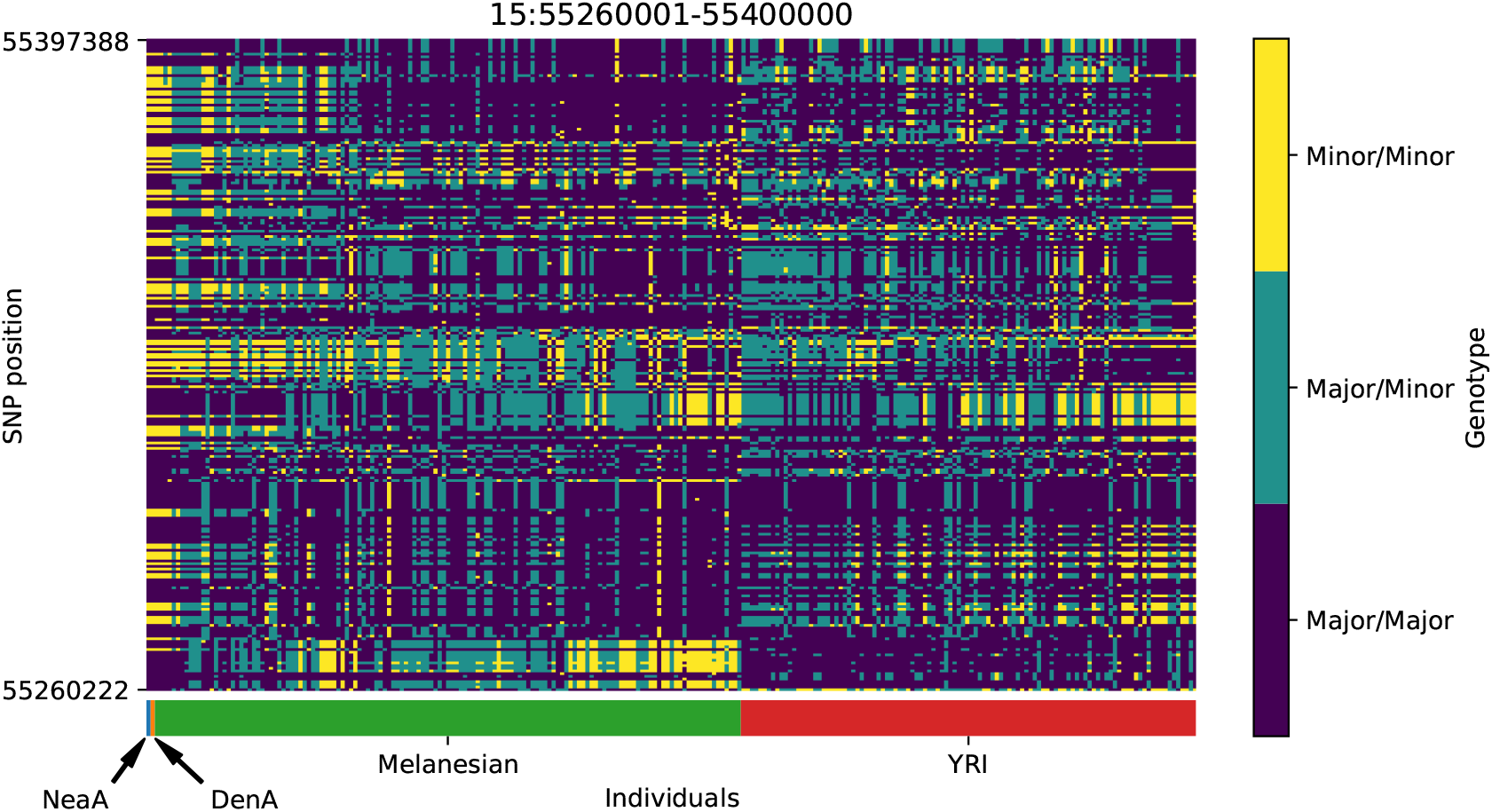
Genotype plot for the candidate region chr15:55260001-55400000 in the Denisovan-into-Melanesian AI scan. Dark blue = homozygote major allele, light blue = heterozygote, yellow = homozygote minor allele. Genotypes within populations are sorted left-to-right by similarity to the Denisovan.

**Figure S43:**
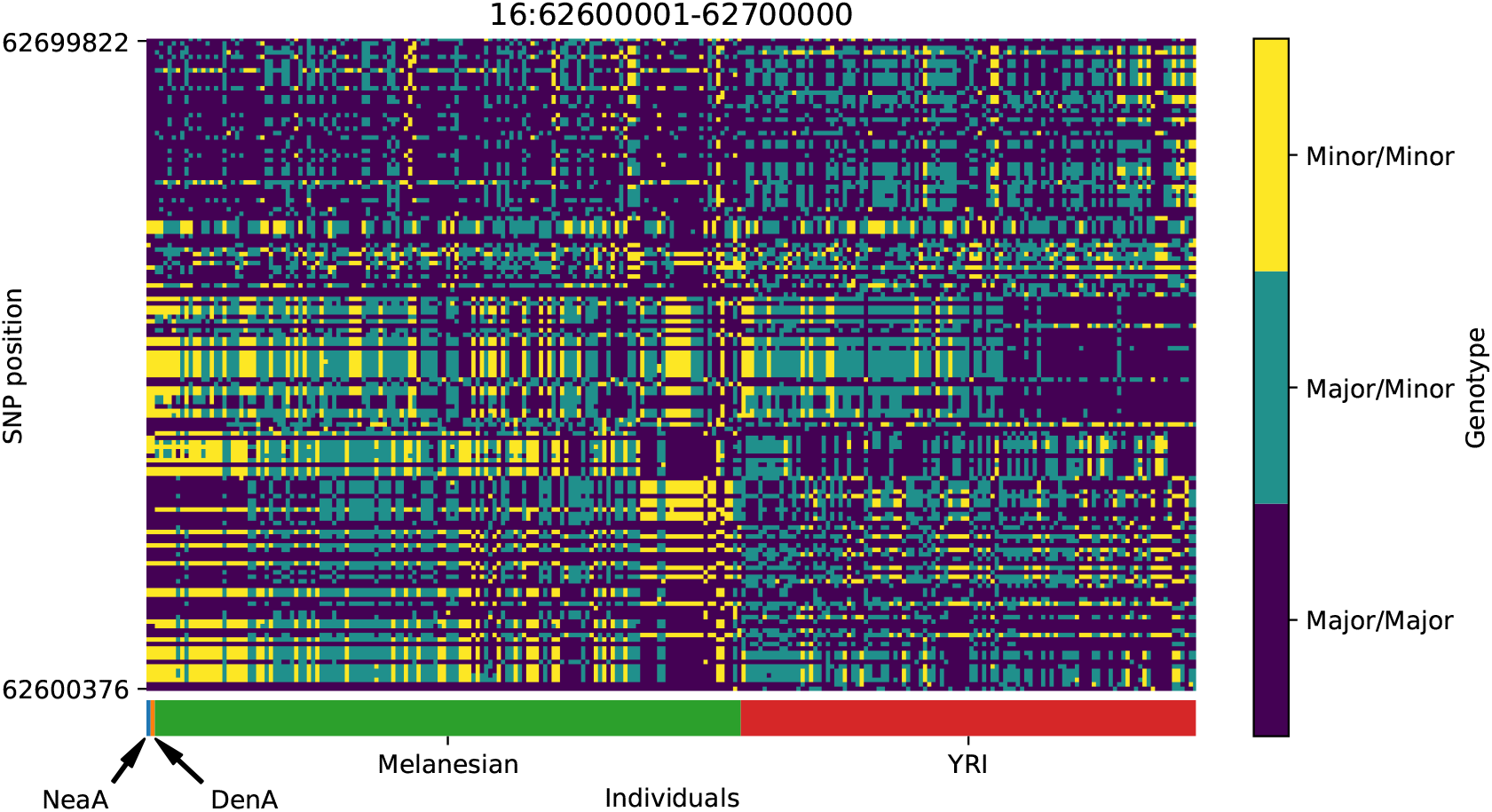
Genotype plot for the candidate region chr16:62600001-62700000 in the Denisovan-into-Melanesian AI scan. Dark blue = homozygote major allele, light blue = heterozygote, yellow = homozygote minor allele. Genotypes within populations are sorted left-to-right by similarity to the Denisovan.

**Figure S44:**
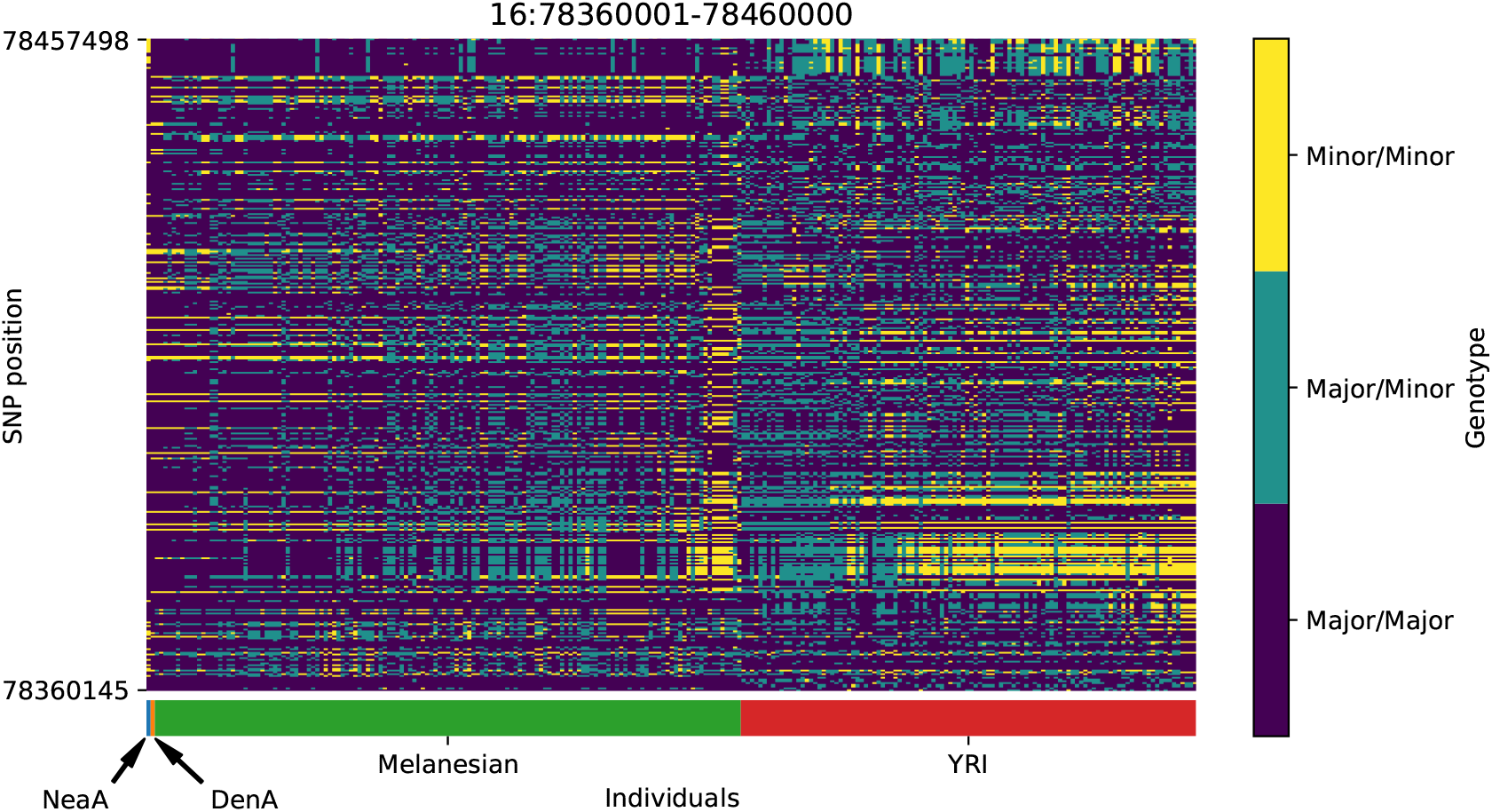
Genotype plot for the candidate region chr16:78360001-78460000 in the Denisovan-into-Melanesian AI scan. Dark blue = homozygote major allele, light blue = heterozygote, yellow = homozygote minor allele. Genotypes within populations are sorted left-to-right by similarity to the Denisovan.

**Figure S45:**
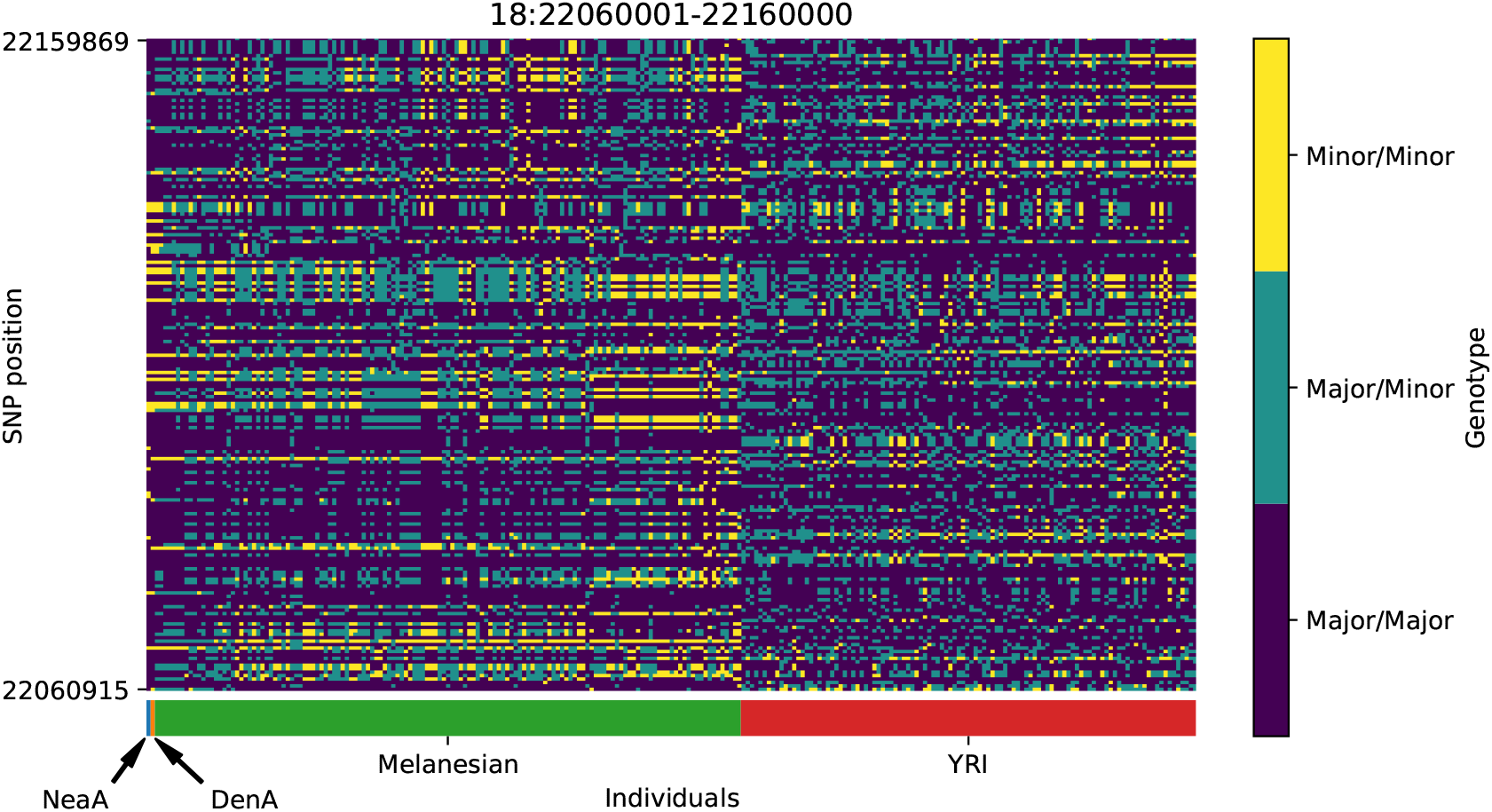
Genotype plot for the candidate region chr18:22060001-22160000 in the Denisovan-into-Melanesian AI scan. Dark blue = homozygote major allele, light blue = heterozygote, yellow = homozygote minor allele. Genotypes within populations are sorted left-to-right by similarity to the Denisovan.

**Figure S46:**
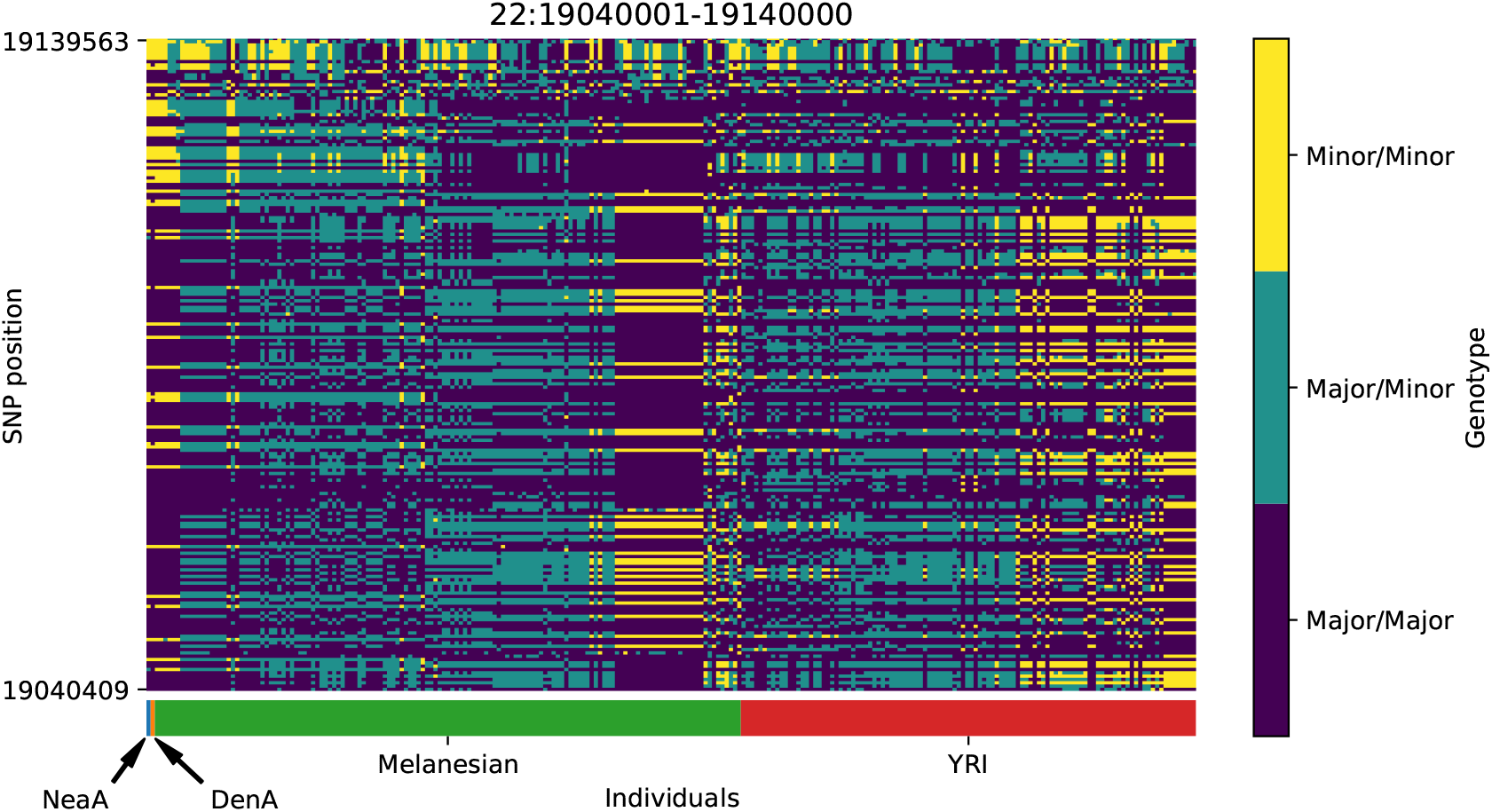
Genotype plot for the candidate region chr22:19040001-19140000 in the Denisovan-into-Melanesian AI scan. Dark blue = homozygote major allele, light blue = heterozygote, yellow = homozygote minor allele. Genotypes within populations are sorted left-to-right by similarity to the Denisovan.

